# Persistent DNA methylation changes associated with prenatal NO_2_ exposure in a Canadian prospective birth study

**DOI:** 10.1101/2023.03.02.530668

**Authors:** Samantha Lee, Hind Sbihi, Julia L. MacIsaac, Padmaja Subbarao, Piushkumar J. Mandhane, Theo J. Moraes, Stuart E. Turvey, Qingling Duan, Amirthagowri Ambalavanan, Michael Brauer, Jeffrey Brook, Michael S. Kobor, Meaghan J. Jones

## Abstract

**Background:** Accumulating evidence suggests prenatal air pollution exposure alters DNA methylation (DNAm), which could go on to affect long-term health. However, it remains unclear whether prenatal DNAm alterations persist through early life. Identifying DNAm changes that persist from birth into childhood would provide greater insight into the molecular mechanisms that most likely contribute to the association of prenatal air pollution exposure with health outcomes such as atopic disease.

**Objectives:** This study investigated the persistence of DNAm changes associated with prenatal NO_2_ exposure (a surrogate measure of traffic-related air pollution) at age one to begin characterizing which DNAm changes most likely to contribute to atopic disease.

**Methods:** We used an atopy-enriched subset of CHILD study participants (N=145) to identify individual and regional cord blood DNAm differences associated with prenatal NO_2_, followed by an investigation of persistence in age one peripheral blood. As we had repeated DNAm measures, we also isolated postnatal-specific DNAm changes and examined their association with NO_2_ exposure in the first year of life. MANOVA tests were used to examine the association between DNAm changes associated with NO_2_ and child wheeze and atopy.

**Results:** We identified 24 regions of altered cord blood DNAm, with several annotated to *HOX* genes. Two regions annotated to *MPDU1* and *C5orf63* were significantly associated with age one wheeze. Further, we found the effect of prenatal NO_2_ exposure across CpGs within all altered regions remained similar at age one. A single region of postnatal-specific DNAm annotated to *HOXB6* was associated with year one NO_2_ and age one atopy.

**Discussion:** Regional cord blood DNAm changes associated with prenatal NO_2_ exposure persist through at least the first year of life, and some of these changes are associated with age one wheeze. The early-postnatal period remains a sensitive window to DNAm perturbations that may also influence child health.

## Introduction

Prenatal and early-life air pollution exposure are associated with an increased risk of childhood asthma, which is characterized by wheezing, difficulty breathing, chest tightness, and bronchial hyper-responsiveness^1, 2^. Often, childhood asthma is preceded by asthma-like symptoms and immune disorders, including wheeze and allergies (atopy), in what is known as the atopic march^3^. This progression involves both genetic and environmental factors and is driven by type 2 inflammation^4^. While the exact mechanisms connecting prenatal air pollution exposure to childhood asthma and atopic disease remain unclear, accumulating evidence suggests that biological embedding of increased oxidative stress and concomitant inflammatory signalling may play a role^5^. Biological embedding refers to the notion that early-life experiences and exposures can “get under the skin” to influence health and development^6^. At the cellular level, this process may involve epigenetic alterations that regulate DNA packaging and expression and impact how cells respond to future stimuli^6^. One example of epigenetic modification is DNA methylation (DNAm), which refers to the addition of a methyl group on the 5’ position of a cytosine in a cytosine-guanine pair (termed “CpG”). Though DNAm patterns are established early in life, they remain malleable in response to environmental exposures. While some of these DNAm perturbations exist only transiently, others persist through development^7^. Given that persistent DNAm alterations are more likely to affect long-term cell function than transient changes, persistent DNAm changes should be of most interest as they are more likely to contribute to development of child asthma following prenatal air pollution exposure than transient DNAm changes^7^.

Despite the notion that persistent DNAm alterations are more likely to contribute to clinically-relevant health outcomes, most studies examining prenatal air pollution exposure only identify DNAm changes at a single time point, typically at birth. These studies suggest that prenatal air pollution exposure at least transiently alters cord blood DNAm in cellular pathways related to oxidative stress and inflammation. For example, a meta-analysis of prenatal NO_2_ exposure across several European cohorts revealed significant cord blood DNAm alterations in *LONP1* and *SLC25A28*, which are related to mitochondrial oxidative stress, as well as in *PVLAP,* which plays a role in leukocyte trafficking^8^. In the EARLI study, prenatal NO_2_ exposure is associated with lower *RNF39* DNAm^9^. The *RNF39* protein is part of Major Histocompatibility Complex (MHC) class I, thus it is biologically plausible that altered *RNF39* DNAm may be associated with changes in immune function^10^, including asthma. More recently, DNAm at *CASP7* was associated with prenatal NO_2_ exposure across all trimesters in a Korean cohort^11^. The *CASP7* gene is a critical mediator of mitochondria-induced apoptosis, thus this finding is consistent with previous observations of altered DNAm in mitochondrial oxidative stress pathways^8, 12^. Finally, prenatal NO_2_ exposure is associated with lower LINE DNAm at age 11 and is also correlated with higher blood pressure at age 11, suggesting that persistent changes in LINE DNAm induced by prenatal NO_2_ exposure may affect long-term child cardio-respiratory outcomes^13^. However, a major limitation of that study was the inability to discern the additional contribution of postnatal air pollution exposure on DNAm and blood pressure.

The postnatal period, especially the first year of life, is considered part of the developmental window that is especially sensitive to biological embedding as DNAm patterns, lungs, and the innate immune system are still developing, and there is an increasing diversity of exposures and experiences^14–16^. In an epidemiological cross-sectional analysis prenatal and postnatal passive cigarette smoke exposures were shown to have independent effects on the risk of negative respiratory health outcomes, supporting the notion that the postnatal period remains a sensitive window to exposures^17^. While several studies have identified positive correlations between postnatal air pollution exposure and child asthma, the role of postnatal-specific DNAm changes in this relationship has not been investigated^18–20^. This knowledge would help clarify the underlying molecular mechanisms connecting postnatal air pollution exposure to respiratory health and provide insight into how they differ from prenatal mechanisms. However, it is not possible to directly measure postnatal-specific changes in DNAm as DNAm measured in peripheral blood represents changes occurring across the prenatal and postnatal periods. Instead, postnatal-specific DNAm changes must be isolated *in silico* by subtracting measures of cord blood DNAm from measures of peripheral blood DNAm.

The purpose of this study was to examine the persistence of cord blood DNAm changes associated with prenatal NO_2_ exposure to identify the biological pathways that most likely to contribute to pollution-associated childhood asthma. To accomplish this goal, we first identified NO_2_-associated DNAm changes at individual sites and local regions using hypothesis-driven and discovery-based approaches. Then, we used linear mixed models to examine whether the effect of prenatal NO_2_ exposure on DNAm remained similar at birth and age one. As we had repeated measures of participant DNAm across time points, we were able to subtract cord blood DNAm measures from DNAm measured at age one, allowing us to isolate and examine postnatal-specific changes in DNAm occurring during the first year of life. This analysis is unique because it helps to demonstrate that the first year of life remains a critical window for the biological embedding of early air pollution exposures in addition to and possibly independent from the prenatal period. Finally, we examined the correlation between significant prenatal DNAm and postnatal-specific DNAm changes with predictors of childhood asthma at age one to provide greater insight into the affected pathways that most likely to impact child health outcomes.

## Methods

### Study population

The CHILD study (CHILD) is a prospective birth cohort designed to investigate the developmental origins of childhood allergic disease, including asthma^21, 22^. Between 2008 and 2012 a total of 3624 eligible mothers were recruited from four major Canadian cities: Vancouver, Edmonton, Winnipeg/Winkler, and Toronto. Eligible participants had to be ≥ 18 years of age (19 years in Vancouver), communicate in English, reside within reasonable proximity to a study centre and plan to give birth there, intend to continue living near a study centre, be willing to provide informed consent, and provide information for two alternate contacts^21^. Children born before 35 weeks gestation, born with major congenital abnormalities or respiratory distress syndrome, conceived by *in vitro* fertilization, resulting from multiple births, and/or who spend less than 80% of nights in the index home were excluded from the study^21^. Each study centre obtained ethics approval from their research ethics board, and CHILD was approved by the Hamilton Integrated Ethics Board (certificate number 07-2929). This study uses data from an atopy-enriched subset of 145 CHILD participants and was approved by the University of Manitoba Research Ethics Board (HS22880)^23^.

### Measurement of maternal characteristics

We included the *a priori* selected maternal variables education length (a surrogate for socioeconomic status) and smoking in our linear regressions to account for biological variation. Maternal characteristics were assessed by CHILD study health professionals at several time points throughout the prenatal period and first year of life^24^. Maternal education length was self-reported at initial intake and used a surrogate measure of socioeconomic status in all models. Prenatal maternal smoking was defined at 18 weeks gestation as an affirmative response to the question “At the present time, how often do you smoke” or to the question “During this pregnancy, did you cut down (but not completely stop) smoking?” or by any non-missing response to the questions “During this pregnancy, did you completely stop smoking?”, “At what week did you stop smoking?”, and “At what week of pregnancy did you cut down smoking”. Postnatal maternal smoking was defined at the end of year one as an affirmative response to the question “At the present time do you smoke?”.

### Air pollution exposure

Average prenatal exposure to outdoor NO_2_ was estimated separately for Vancouver, Edmonton, Winnipeg, and Toronto as previously described^25–27^. Briefly, city-specific land use regression (LUR) models were used to estimate outdoor NO_2_ concentration at the residential location of each participant for the prenatal period (estimated time of conception until birth) and the first year of life (birth until age one)^22, 28^. All models included information on land use, roads and traffic, population, physical geography, and meteorology; consequently, in addition to being related to combustion, NO_2_ is typically considered an indicator of traffic-related air pollution (TRAP)^28^. Thus, for CHILD subjects whose NO_2_ exposures are estimated to be high, NO_2_ exposure broadly represents exposure to a mixture of combustion-related emissions, including a range of particles and gases with varying potential toxicities. For participants that reported a change in primary residence during the study period (N=23), outdoor NO_2_ estimates were computed using a time-weighted average of exposure at each reported residence^28^. All outdoor NO_2_ estimates were temporally adjusted on a biweekly basis using local fixed-site ambient monitoring data collected by the National Air Pollution Surveillance (NAPS) network^28^. Participants with neither prenatal or year one outdoor NO_2_ estimates were omitted from the study (N = 2 participants)^28^.

### DNA methylation measurement and quality control

At each delivery site umbilical cord blood was collected into heparinized vacuum tubes. For each individual, heparinized blood samples were combined in a single 50 mL conical tube, which was processed to obtain cord blood mononuclear cells (CBMCs)^21^. CBMC pellets were resuspended in Buffer RLT-Plus (600 uL per mL; Qiagen) and stored at −80C for later nucleic acid extraction. Two peripheral blood aliquots (1.5 mL to 3 mL each) collected at age one were subjected to the same procedures to obtain peripheral blood mononuclear cells (PBMCs)^21^. CBMC and PBMC samples were used to generate DNAm data as previously described^21^. Briefly, 750 ng of extracted DNA (Qiagen DNAeasy kit) was bisulfite converted (Zymo EZ DNAm Gold kit), followed by application of 160 ng of bisulfite-converted DNA to the Illumina HumanMethylation 450k array. All nucleic acid procedures followed the manufacturer’s instructions.

Preprocessing and quality control of DNAm data was performed in *minfi*, except where noted^29^. Sample and probe quality were assessed using detection p-values. One cord blood sample with 20751 bad probes (detection p-value > 0.01) was removed prior to normalization. Background correction and dye-bias normalization were accomplished using *preprocessNoob*, followed by probe bias correction using *BMIQ* from the *wateRmelon* package^30, 31^. Next, we removed 8,482 probes with a detection P > 0.01 and/or with fewer than 3 beads contributing to signal, 15,333 probes with a known single nucleotide polymorphism (SNP), 10,860 probes localized to sex-chromosomes, and 26,183 cross-reactive probes leaving 424,644 of the initial 485,512 probes^32^. Quality control was performed to confirm sample labelling (by examining participant expected sex) and replicate quality. All sample replicates (n=3 cord blood samples with 3-4 replicates each) were of good quality based on strong and distinct methylated and unmethylated β-value peaks. Thus, one replicate from each set was randomly selected to include in DNAm analysis.

Measures of DNAm are quantified as the ratio of methylated probe intensities to intensities of all measured probes for a given CpG, known as β-values, or as M-values, which are log-transformed β-values^33^. Both β-values and M-values are commonly used in studies investigating epigenome-wide DNAm changes. M-values are often regarded as more statistically valid as they are homoscedastic, but they are not biologically interpretable^33^. In contrast, the 0-1 scale of β-values corresponds to the methylation percentage of a given CpG site^33^. Additionally, as β-values are linear they can be added or subtracted from one another without distorting DNAm measurements. Therefore, we used DNAm β-values in this study as they were compatible with our analyses of examining the occurrence of DNAm changes in the first year of life.

Specifically, we were interested in examining the effects of NO_2_ exposure during the first year of life on postnatal DNAm alterations; however, it is not possible to measure postnatal-specific DNAm changes directly. This is because peripheral blood DNAm interrogated in childhood represents DNAm changes occurring across both the prenatal and postnatal periods. To isolate postnatal-specific DNAm changes we used a novel method wherein we subtracted cord blood β-values from age one blood β-values and used the resulting differences in β-values in our analyses, termed “postnatal-specific” β-values. Postnatal-specific β-values represent DNAm changes occurring specifically in the first year of life.

### Estimation of cord blood monocyte proportions

DNAm is cell type specific and differences in cell type composition between samples may lead to spurious findings unless accounted for in linear models. Each cell type has a unique DNAm signature; therefore, DNAm patterns can be used to estimate cell type proportions in blood samples using a reference-based deconvolution method^34^. As cell types were not directly measured (e.g. by flow cytometry) for most CHILD participants, we estimated cord blood cell type proportions using a reference-based method^34^. Specifically, we used the *FlowSorted.CordBloodCombined.450k* cord blood reference set and the *estimateCellCounts2* function from the *FlowSorted.Blood.EPIC package* to estimate the relative proportions of 6 cell types (CD4 T-cells, CD8 T-cells, B-cells, NK cells, monocytes, and nucleated red blood cells) in CBMC samples either using default cord blood probe selection methods (“any” top 100 probes and “both” the 50 top hypomethylated and 50 top hypermethylated probes) or using a predefined set of *IDOL* probes selected to optimize cords blood deconvolution^35–37^. We based our probe selection method choice on whether the probe selection parameter produced cell types estimates within expected proportions and whether cell type estimates differed between cord blood between probe selection parameters based on Pearson correlation coefficients (r) of direct comparisons (**Supplementary Figure 1**). Based on these comparisons, we proceeded to estimate CBMC proportions (**Supplementary File 1**) using the default “any” parameter of *estimateCellCounts2* for deconvolution of cord blood.

The compositional properties of cell type estimates require they sum to 100% and can introduce multicollinearity into linear regressions^38^. To remove constraints imposed by cell type composition, we performed isometric-log ratio (ILR) transformations followed by robust principal component analysis (rPCA) on predicted CBMC proportions. We subsequently included the first four principal components (PCs) in linear regressions to account for ∼90% of the variance due to differences in estimate cord blood cell type proportions and removed 565 unique probes used to estimate cord blood cell proportions (**Supplementary File 2**) from the total number of cord blood DNAm probes, leaving 424079 probes remaining in our epigenome-wide analysis of prenatal NO_2_ exposure

### Estimation of age one peripheral blood monocyte proportions

There is no existing reference set for the estimation of cell proportions in age one blood. Thus, we conducted a preliminary analysis comparing cord blood and adult blood deconvolution procedures to determine which reference set (cord or adult) and probe selection parameters (“any”, “both”, or “IDOL”) are appropriate for age one PBMC prediction^36, 37^. We based our selection on whether each combination of reference set and probe selection parameter produced cell types estimates within expected proportions and whether cell type estimates differed between cord blood or adult blood reference sets based on Pearson correlation coefficients (r) of direct comparisons (**Supplementary Figure 2**). Cord blood and adult blood reference sets using the “any” probe selection parameter performed similarly (r≥0.92). Therefore, we proceeded with the *FlowSorted.CordBloodCombined.450k*^35^ reference set using the *estimateCellCounts2* function with the default cord blood probe selection parameter “any” to estimate the relative proportions of 5 cell types (CD4 T-cells, CD4 T-cells, B-cells, NK cells, monocytes) in age one PBMC samples (**Supplementary File 3**) for consistency with our CBMC deconvolution procedure. We performed ILR-transformation followed by rPCA on estimated PBMC proportions and used the first three PCs in linear regressions to account for >90% of variation related to differences in estimated cell type proportions. We removed the 565 unique probes used to estimate age one blood cell proportions (**Supplementary File 4**) from the total number of probes measuring age one blood DNAm for a total of 424079 remaining probes.

### Variation due to cell type in postnatal-specific measures of DNA methylation

The analysis of postnatal-specific DNAm changes requires consideration of variation in both CBMC and PBMC cell types. In this case, ILR-transformed CBMC and PBMC cell types were combined prior to rPCA and analyzed together. We included the first six PCs accounting for ∼90% of the variance due to combined CBMC and PBMC cell types in our analyses examining postnatal-specific DNAm changes. We removed the 565 unique probes shared between the probes used to estimate cord blood and age one blood cell proportions from the total number of probes passing quality control for a total of 424079 remaining probes interrogating postnatal-specific DNAm.

### Variation due to cell type in investigations of DNA methylation persistence

We previously generated cord blood and peripheral blood cell-type PCs separately to correct for differences in cell proportions at birth and age one. The top cord bloods PCs do not necessarily capture variation due to the same cell types as the top age one blood PCs, and thus it is not appropriate to model coefficients across cord blood PCs and age one blood PCs in linear mixed models using to investigate DNAm persistence. Therefore, we combined estimates of cord blood cell proportions with estimates of age one peripheral blood cell proportions in a row-wise manner. To facilitate combination of these data, we created a new variable in the estimates of age one peripheral blood cell proportions for nRBCs and set this value to zero as this cell type is not normally present in healthy children at age one^39^. Combining estimates of cord blood and peripheral blood cell proportion in this manner ensures the resulting cell type PCs reflect variation due to the same cell types across both time points. Afterwards, we conducted ILR and rPCA on the row-wise combined estimates of cell proportions and included the first three PCs accounting for >90% of the variance in our linear mixed models examining the persistence of cord blood DNAm at age one.

### Genotyping and population substructure

A total of 2967 CHILD participants were genotyped using the Illumina HumanCoreExome BeadChip, including the 145 participants used in this study. Quality control and imputation were performed on all genotyped participants together before selection of the subset of participants used in this study. Briefly, cord blood was collected at birth as described above. The QiaSymphony automated large sample nucleic acid purification system was used to extract DNA for genotyping (Genetic and Molecular Epidemiology Laboratory, McMaster University)^21^. Quality control was carried out at the subject and SNP levels using Plink (v1.9)^40^. Specifically, we omitted 39 participants with missing genotypes >10% and excess heterozygosity (±2 standard deviations), 56 participants with discordant sex, and 37 participants that were likely third-degree relatives as identified by pairwise Identify-By-Descent (IBD coefficient 0.185)^41^. At the SNP level QC, we selected for the autosomal variants, and omitted variants exceeding missingness of >0.05. A total of 2835 subjects remained with 515,033 variants passing QC. Imputations were performed using the HRC (r1.1 2016) reference panel implemented through the Michigan Imputation server^42^. Chromosomes were phased using ShapeIT (v2.r790), then SNPs were imputed via the Minimac algorithm^43, 44^. This resulted in 28 million SNPs, which included 5.4 million common SNPs (MAF >0.05) that passed the imputation quality score R2>0.3^44^. Quality control measures as above were applied to the imputed markers and no markers or subjects were excluded post-imputation. Afterwards, we performed a PCA of CHILD study genotype data (N=2835) to assess population substructure. We estimated the eigenvectors of CHILD participants using the EIGENSTRAT–SMARTPCA tool^45–47^. We estimated that the first 10 eigenvectors account for >99.9% of the total variance among subjects. We included the first three genotyping PCs in our linear regressions to account for differences related to genetic ancestry as the first three genotyping PCs resulted in clear separation of participants used in this study (**Supplementary Figure 3**).

### Candidate gene analyses of prenatal and year one NO_2_ exposure

The underlying mechanisms of prenatal air pollution exposure include increased oxidative stress and inflammatory signalling and the hypothesized biological embedding of these events in DNAm may contribute to atopic disease in childhood^48^. To investigate this notion, we examined prenatal and postnatal-specific DNAm changes associated with prenatal and year one NO_2_ exposure, respectively, across a list of 59 *a priori*-determined candidate genes hypothesized to participate in the cellular pathways activated by air pollution exposure or that were previously identified in epigenome-wide association studies of prenatal air pollution (**Supplementary Table 1**)^8, 9, 49–53, 53–59, 59–66^. We also examined whether two previously identified DNAm changes within the genes *LONP1* and *SLC25A2* could be replicated in our analysis^8^. Candidate gene analyses were conducted using multivariable linear models using the *lm* function. Our candidate gene analysis of prenatal NO_2_ exposure included the following covariates: sex, gestational age, prenatal maternal smoking, maternal education length, cell type, and genetic ancestry. We additionally included 17 surrogate variables (SVs) to account for technical variation, as determined by the *SVA* package (**Supplementary Figure 4**)^67^. When examining postnatal-specific DNAm changes in our candidate genes, we included variables to adjust for sex, year one maternal smoking, maternal education length, cell type, genetic ancestry, and technical variation using 15 SVs as determined by the SVA package (**Supplementary Figure 5**). We did not include study site as a covariate for either model, or in any of the models below, as most of the variation in prenatal and year one NO_2_ estimates occurred between study sites (i.e. including study site would limit our power to detect small DNAm changes; **Supplementary Figures 6-8**). The prenatal and postnatal-specific candidate gene analyses were corrected for multiple comparisons using the Bonferroni method. Significant DNAm changes within candidate genes were defined as having an adjusted p<0.05 and displaying an absolute effect size >0.025% DNAm.

### Epigenome-wide analyses of prenatal and year one NO_2_ exposure

We conducted epigenome-wide analyses to identify novel cord blood and postnatal-specific DNAm changes associated with prenatal and year one NO_2_ exposure, respectively. Both analyses used the *limma* R package to conduct multivariable linear regressions on DNAm β-values. Models investigating effects of prenatal NO_2_ exposure on cord blood DNAm included covariates to adjust for sex, gestational age, prenatal maternal smoking, maternal education length, cell type, and genetic ancestry, and technical variation using 17 SVs as suggested by the SVA package (**Supplementary Figure 4**)^67^. Models examining the effect of year one NO_2_ exposure on postnatal specific DNAm included covariates to adjust for sex, year one maternal smoking, maternal education length, cell type, genetic ancestry, and technical variation using 15 SVs as determined by the *SVA* package (**Supplementary Figure 5**)^67^. For each analysis the resulting p-values were adjusted for false discovery using the Benjamini-Hochberg method. Individual CpGs were considered significant if they surpassed FDR p<0.05 and exhibited an absolute effect size >0.025% DNAm.

### Identification and characterization of differentially methylated regions associated with prenatal and year one NO_2_ exposure

Differentially methylated regions (DMRs) refer to adjacent CpGs that display altered DNAm between phenotypes or exposures^68^. Region-based analyses are advantageous as they are typically more statistically powerful than epigenome-wide analyses as they can leverage information from adjacent CpGs. Additionally, DMRs can offer greater insight into biologically-relevant DNAm changes, as regions of altered DNAm are more likely to affect cell function than individual altered CpGs. We used the *combp* function from the *ENmix* R package to identify cord blood and postnatal-specific DMRs associated with prenatal and year one NO_2_ exposure, respectively^69, 70^. The *combp* method spatially adjusts p-values obtained from an epigenome-wide investigation to identify DMRs^69^. We considered *combp* DMRs to be statistically significant if they surpassed FDR p<0.05 and exhibited a mean absolute effect size >0.025% DNAm.

We were interested in whether observed DNAm changes represented gene-by-environment effects, but lacked power to test whether identified CpGs represented methylation trait quantitative loci (mQTLs)^71^. Instead, we cross-checked all CpGs contained with cord blood (N=151 CpGs) and postnatal-specific (N=6 CpG) DMRs with mQTLdb, an online database of methylation quantitative trait loci (mQTLs)^72^. We compared cord blood DMR CpGs identified in this study with mQTLs surpassing p<1E-14 and exhibiting an effect size >0.02% (**Supplementary File 5**)^72^. Postnatal-specific DMR CpGs represent changes that occur in the first year of life; however, the average age of childhood mQTLs reported on mQTLdb is 7.5 ± 0.2 years, which represents a different developmental stage^72^. Therefore, we compared our postnatal-specific DMR CpGs with mQTLs reported at both birth and childhood on mQTLdb (**Supplementary Files 6 and 7**).

Gene ontology analysis can provide insight into the biological pathways that are affected by regional changes in DNAm. We performed gene ontology analysis on cord blood DMRs using *goRegion* from the *missMethyl* R package with default settings^73^. We considered gene ontology terms to be significant when FDR p<0.05. We did not perform gene ontology analysis on postnatal-specific DMRs as we identified only one DMR at this time point.

### Persistence of individual and regional cord blood DNA methylation changes at age one

Previous studies have not examined whether cord blood DNAm changes associated with prenatal air pollution exposures persist through early childhood development. Persistent DNAm changes are of interest because they are more likely to impact child health than transient changes^7^. Therefore, we aimed to conduct a novel analysis examining whether DNAm changes present at birth persisted at age one in CHILD participants. To determine persistence of cord blood DNAm changes at age one, we used linear mixed effects models to examine whether the coefficient of prenatal NO_2_ exposure significantly differed in cord blood at birth and peripheral blood at age one). We applied these linear mixed models to the top 100 CpGs identified in our epigenome-wide analysis of prenatal NO_2_ exposure that exhibited an absolute effect size >0.025% DNAm and FDR p <0.05. In a separate analysis we applied linear mixed models to all individual CpGs (N= 151) contained within cord blood DMRs associated with prenatal NO_2_ exposure. In both analyses, linear models were run using the *lme4* package. Specifically, linear mixed models included fixed effects to account for prenatal NO_2_ exposure, time point (birth or age one), biological sex, gestational age, maternal education status, maternal smoking status, and cell type PCs. Additionally, we included SVs to account for technical variation across both the birth and age one time points. We combined cord blood β-values and age one β-values in a row-wise manner before conducting surrogate variable analysis to generate SVs that capture variation due to the same measured and unmeasured variables across both time points and included the first 16 SVs in our linear mixed models (**Supplementary Figure 9**). Linear mixed models also included interaction terms between time and NO_2_ exposure, and time and gestational age to reflect that the effect of these variables likely changes over time. Finally, a random intercept term was included to account for repeated measures across participants. We considered the effect of prenatal NO_2_ exposure on DNAm to persist if the p-value of interaction term between time and NO_2_ failed to reach significance at p<0.05 after Bonferroni adjustment.

### Global DNA methylation analyses

Global DNAm refers to the average methylation level across a genome. Often, DNAm of long interspersed elements (LINEs) is used as a surrogate measure of global DNAm since LINEs are tightly regulated by DNAm and sensitive to environmental exposures^13, 74, 75^. Here, we examined both average DNAm across all measured CpGs in a sample as well as average DNAm across all probes mapping to L1 LINEs using the Data Integrator function of the University of California Santa Cruz with annotation from Repeat Masker and the ENCODE HAIB Methyl450 track^76–78^. We identified 7302 probes mapping to L1 LINEs in cord blood and postnatal-specific DNAm, respectively. We assessed the correlation between both measures of “global” DNAm in cord blood and postnatal-specific DNAm using linear regressions with covariates to correct for biological and technical variation as outlined in our candidate gene and epigenome-wide analyses. We defined a significant alteration in either measure of “global” DNAm as p<0.05. We did not include an effect size cut-off for biological relevance as even a small change in genome-wide or L1 LINE DNAm during development may have a large impact on cell function and health outcome^13, 79^.

### Linkage of NO_2_-associated differentially methylated regions to asthma phenotype

We examined whether any of the identified altered regions of prenatal or postnatal-specific DNAm were associated with predictors of child asthma, including age one wheeze and atopy, to provide greater insight into their biological relevance^3^. We used previously defined classifications of atopy and wheeze as primary asthma-related outcomes^3^. Briefly, age one atopy was defined as a positive skin prick test (wheal diameter ≥2 mm compared to glycerin control) to any of 10 tested allergens including Alternaria tenuis, cat hair, dog epithelium, Dermatophagoides pteronyssinus, Dermatophagoides farinae, German cockroach, peanut, soybean, egg white, and cow’s milk^3^. Age one wheeze was defined as a caregiver reporting child wheezing with or without a cold in the first year of life via questionnaires administered at three, six, and 12 months, or if a CHILD clinician reported wheezing during the age one clinical visit^80^. The relationship between prenatal and postnatal-specific DMRs and age one atopy or wheeze was assessed using MANOVA tests (with the Pillai–Bartlett test statistic) that included covariates to correct for biological and technical variation as specified in our epigenome-wide analyses. Resulting p-values were adjusted for multiple test comparisons using the Bonferroni method with significance defined as p<0.05.

### Effect of birth month on the relationship between NO_2_ exposure and DNA methylation

We hypothesized that the effect of NO_2_ exposure on DNAm may vary by birth month as levels of traffic-related air pollution, including NO_2_, vary annually with higher levels occurring during colder months^81^. Based on previous findings investigating prenatal air pollution exposure timing and asthma risk, we postulated that participants who were born in a month that exposed them to the highest levels of NO_2_ during a critical period of prenatal development (e.g. those born in warmer summer months are hypothesized to be exposed to higher levels of outdoor NO_2_ during the first trimester of pregnancy in colder winter months) would exhibit a greater magnitude of DNAm change^82^. To investigate this hypothesis, we conducted a sensitivity analysis to investigate whether the effect of prenatal or year one NO_2_ on DNAm varied by birth month at the top 10 CpGs identified in our prenatal and postnatal specific analyses, respectively. Specifically, we re-ran multivariable linear regressions with an interaction term between NO_2_ exposure and birth month and used ANOVA likelihood ratio tests to examine whether modelling the effect of prenatal NO2 exposure on DNAm by birth month significantly reduced the residual sum of squares compared to models that did not include birth month. We considered the interaction model fit to be significantly different when p<0.05.

## Results

A total of 144 participants with DNAm data available at birth and age one remained after preprocessing and normalization. Of these, 128 participants had complete prenatal NO_2_ exposure data, with a median exposure of 12.0 parts per billion (ppb; **Table 1**). These 128 participants were included in our analysis of prenatal NO_2_ exposure in cord blood DNAm. We used a subset (N=124) of these 128 participants that also had available age one peripheral blood DNAm data to examine the persistence of cord blood DNAm changes at age one. This subset exhibited the same distribution of prenatal NO_2_ exposure as the 128 participants included in our analyses of prenatal NO_2_ exposure (**Table 1**). In contrast, participants (N=125) that were included in our postnatal analysis were exposed to slightly lower levels of NO_2_ air pollution during the first year of life, with a median postnatal NO_2_ exposure of 10.2 ppb (**Table 1**). Paired t-tests indicated that the difference in prenatal and year one NO_2_ exposure means was statistically significant for participants (N=122) that had estimates available for both time points (p= 1.8e-05; **Supplementary Figure 10**). However, it is important to note that medians of both prenatal and postnatal NO_2_ exposure fall within the same air quality standard defined by the Canadian Council of Ministers of the Environment, but above the most current recommendations from the World Health Organization, and are therefore expected to have similar effects on health^83, 84^. Participants included in our analyses of cord blood DNAm, persistence of cord blood DNAm changes, and postnatal-specific DNAm changes exhibited similar demographic profiles, including maternal ethnicity and maternal education, to one another and the full CHILD cohort^24^. However, a slightly higher percentage of participants in our analysis of prenatal NO_2_ exposure (9.4%; **Table 1**) reported maternal smoking during the prenatal period compared to full CHILD cohort (5.4%)^24^.

**Table 1.**
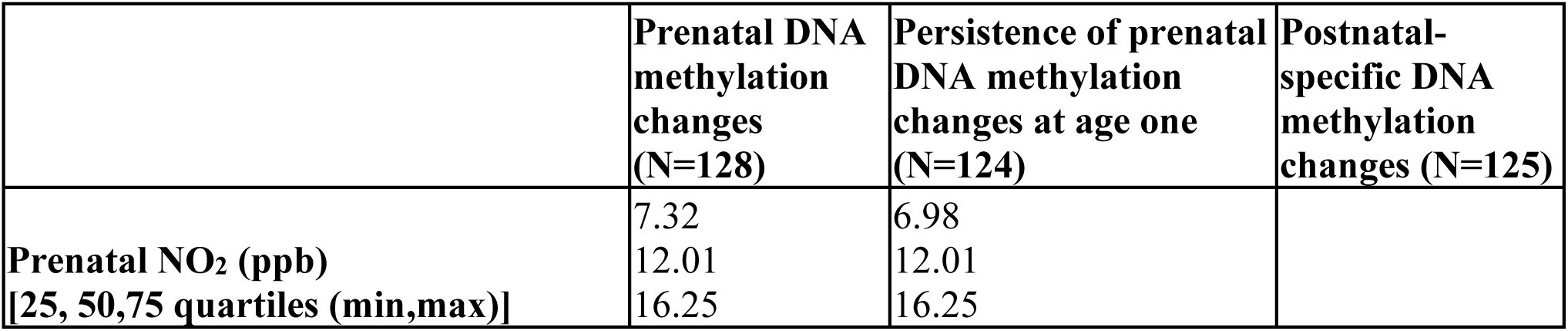
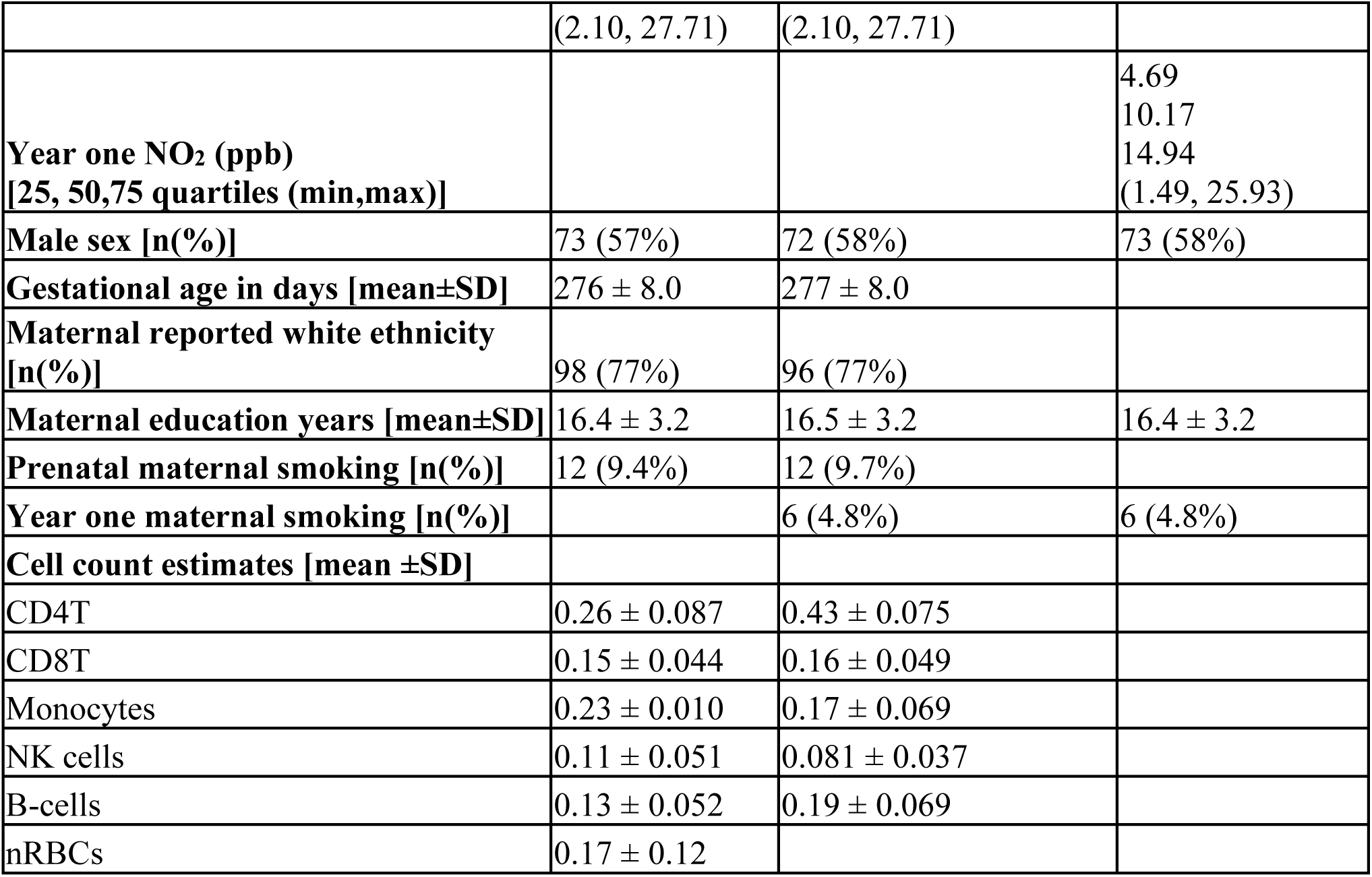
Study population characteristics

As NO_2_ air pollution exposure induces oxidative stress and inflammation, we began our analyses by investigating DNAm across 59 candidate genes encompassing 877 CpGs related to these cellular pathways. This *a priori* list included two previously identified genes, *LONP1* and *SLC25A28*, for replication^8^. No CpGs annotated to the candidate genes were significantly associated with prenatal NO_2_ exposure in cord blood (**Supplementary File 8**), or with year one NO_2_ exposure in postnatal-specific DNAm (**Supplementary File 9**). Further, we found no associations between cord blood DNAm and prenatal NO_2_ exposure at cg12283362 (*LONP1*) and cg08973675 (*SLC25A28*), nor did we observe a relationship between year one NO_2_ exposure and postnatal-specific DNAm at these CpGs (**Figure 1A and B**).

**Figure 1.**
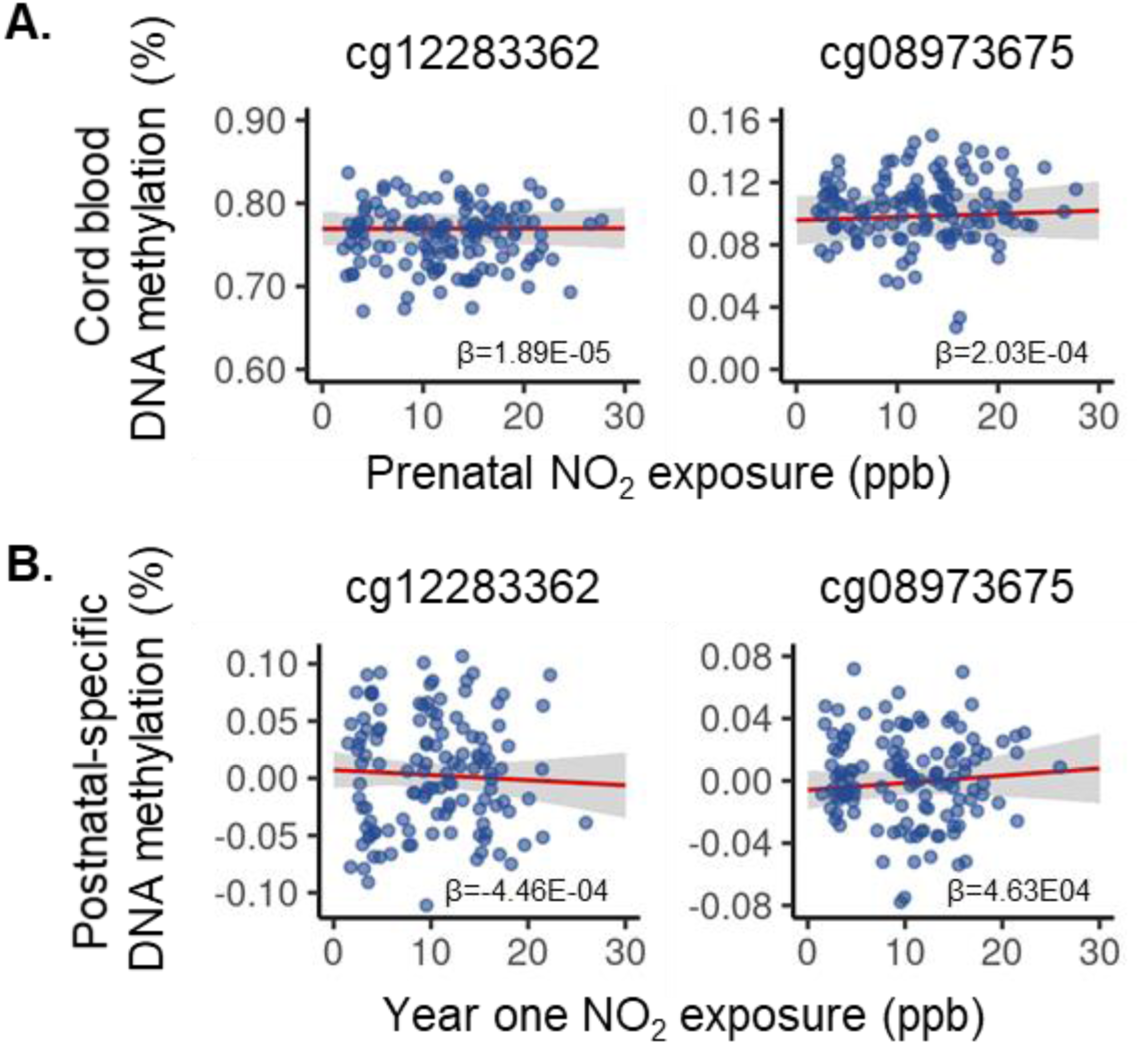
Previously identified cord blood DNA methylation (DNAm) changes in *LONP1* (cg12283362) and *SLC25A28* (cg08973675) do not replicate in CHILD. (**A**) In cord blood, neither DNAm at cg12283362 (adjusted p=1; effect size=4.83e-4; β=1.89e-5) or cg08973675 (adjusted p=1; effect size=5.19E-3; β=2.03E-4) was significantly associated with prenatal NO_2_ exposure in CHILD participants (N=128). (**B**) Postnatal-specific measures of DNAm at cg12283362 (adjusted p=1; effect size=-1.10E-2; β=-4.46E-5) and cg08973675 (adjusted p=1; effect size=1.12E-2; β=4.63E-4) were not significantly associated with year one NO_2_ exposure in CHILD participants (N=125). Red lines represent the estimated marginal effect of NO_2_ exposure on DNAm with a slope equal to coefficient of NO_2_ exposure (β). The surrounding shaded area represents the 95% confidence interval of the marginal effect.

Epigenome wide analyses can provide insight into novel DNAm changes that may not be captured in a hypothesis-driven candidate gene approach. Therefore, we next investigated epigenome-wide DNAm changes associated with prenatal NO_2_ exposure in cord blood (N=128), and with year one NO_2_ exposure in postnatal-specific DNAm (N=125). We did not observe any significant DNAm alterations associated with prenatal NO_2_ exposure in our epigenome-wide analysis of cord blood at 5% FDR and using a biological cut-off of an absolute effect size >0.025% DNAm (**Supplementary Figure 11; Supplementary File 10**). Similarly, we did not observe any significantly DNAm changes in our epigenome-wide analysis of year one NO_2_ exposure and postnatal-specific DNAm at 5% FDR and using an absolute effect size cut-off of >0.025% DNAm (**Supplementary Figure 12; Supplementary File 11**).

The small number of available participants and relatively small range of NO_2_ exposure in our epigenome wide analyses likely limited our power to detect individual cord blood DNAm changes. Analysis of DMRs is often more statistically powerful than epigenome-wide investigations as it can leverage information from adjacent CpGs. Additionally, DMRs are more likely to have a biological effect than individual CpGs. Thus, we examined whether prenatal NO_2_ exposure and year one NO_2_ exposure were associated with regional DNAm changes in cord blood DNAm and postnatal-specific DNAm measures, respectively. At 5% FDR we identified 24 cord blood DMRs exhibiting an absolute mean effect size of prenatal NO_2_ exposure >0.025% DNAm (**Figure 2; Supplementary Table 2**, **Supplementary Figures 13-36**). These regions were not enriched for gene ontology terms; however, we observed several cord blood DMRs containing CpGs annotated to developmentally relevant genes, such as those in the HOX family (**Supplementary Table 2**). We crossed checked the 151 CpG probes contained within the 24 cord blood DMRs with previously reported mQTLs (**Supplementary File 5**) present at birth in the ARIES cohort to assess gene-by-environment effects^72^. We observed that >50% of CpGs were reported as mQTLs in nine of the 24 cord blood DMRs (**Table 2; Supplementary File 12**). In contrast, we identified only a single postnatal-specific DMR associated with year one NO_2_ exposure. This DMR contained 6 CpG probes annotated to *HOXB6* (**Figure 3; Table 3; Supplementary File 13; Supplementary Figure 37**). We examined whether CpGs within this postnatal-specific DMR were previously reported as mQTLs at birth or childhood on mQTLdb (**Supplementary Files 6 and 7**). We observed no CpGs within this DMR were previously reported as mQTLs at birth or during childhood using statistical and biological cut-offs of p<1E-14 and an effect size >0.02% to filter CpGs on mQTLdb.

**Figure 2.**
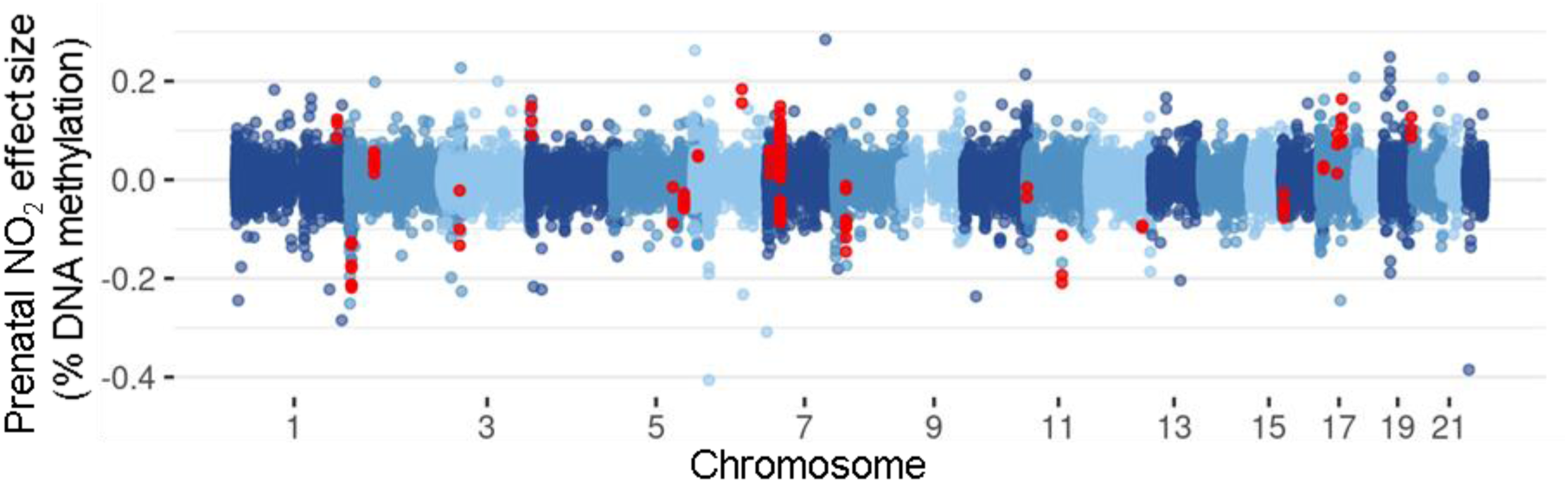
A total of 24 cord blood differentially methylated regions (DMRs) were significantly associated with prenatal NO_2_ exposure. Cord blood DMRs were identified using *combp* and considered significant if they surpassed 5% FDR and exhibited a mean absolute effect size of prenatal NO_2_ exposure >0.025% DNAm. The probes (N=151) contained within these DMRs are highlighted in red.

**Figure 3.**
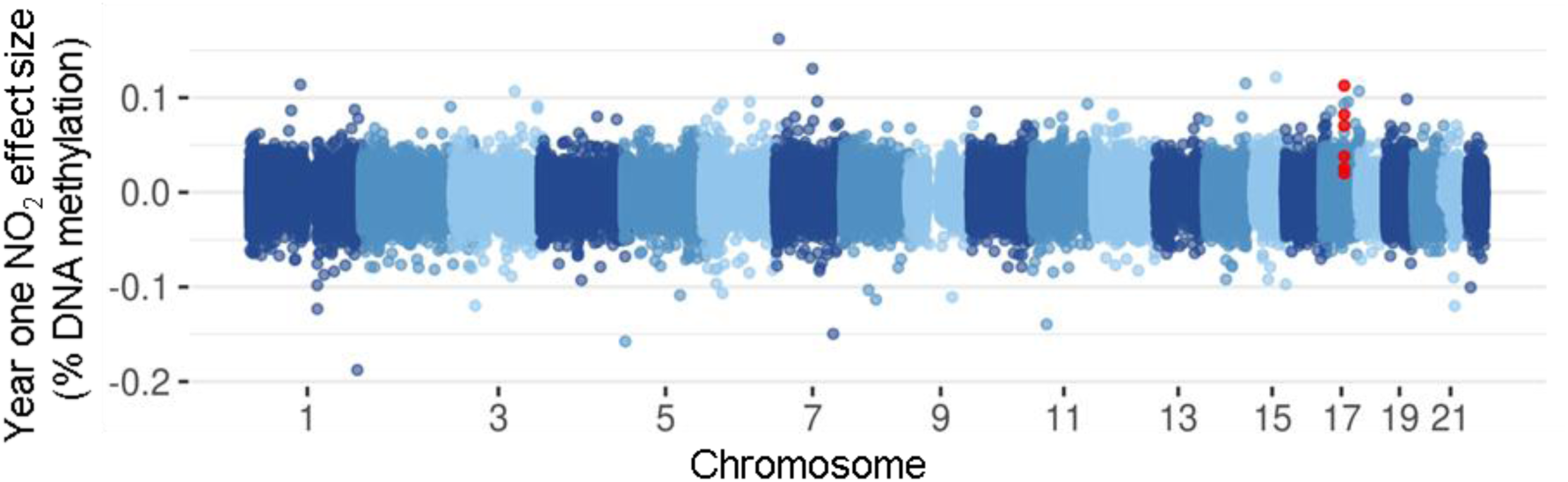
A single postnatal-specific differentially methylated region (DMR) was associated with year one NO2 exposure. Postnatal-specific DMRs were identified using *combp* and considered significant if they surpassed 5% FDR and exhibited a mean absolute effect size of prenatal NO_2_ exposure >0.025% DNAm. The probes (N=6) contained within the postnatal-specific DMR we identified are highlighted in red.

**Table 3.**
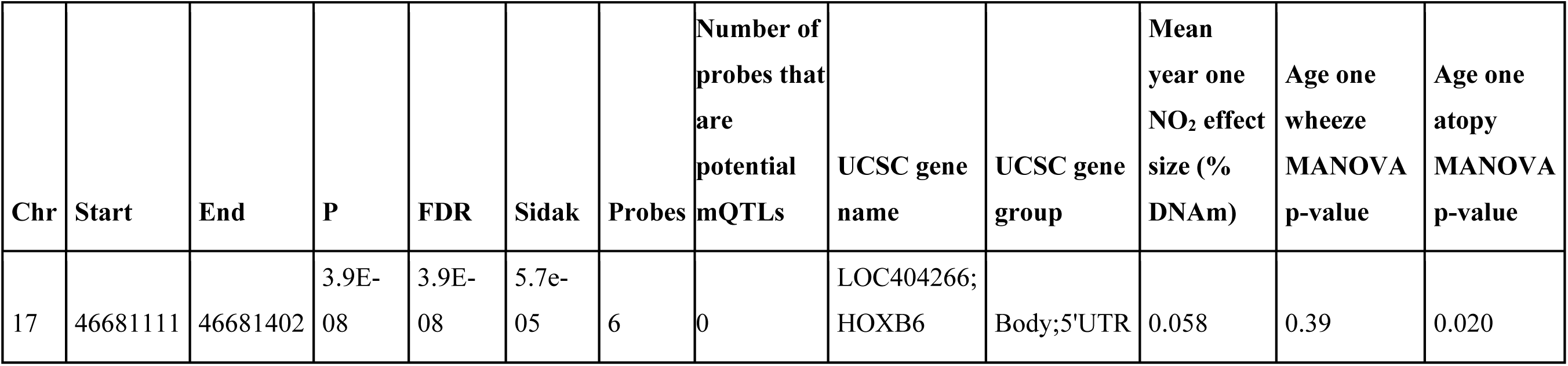
Genomic information of the identified postnatal-specific differentially methylated region significantly associated with year one NO2 exposure.

Persistent DNAm changes are hypothesized to have a greater impact on long term health outcomes than transient DNAm changes^7^. Therefore, we examined whether the cord blood DNAm changes we observed persisted at age one. Specifically, we used linear mixed models to investigate whether the absolute effect size magnitude of prenatal NO_2_ exposure remained similar between birth and age one at the top 100 cord blood CpGs identified in our epigenome-wide analysis of prenatal NO_2_ exposure and at the 151 CpGs contained within significant cord blood DMRs. We investigated individual and regional DNAm changes separately to examine differences in persistence across these perturbations. For discussion purposes we stratified CpGs into those exhibiting lower DNAm at birth and those exhibiting higher DNAm at birth to more readily convey how DNAm is evolving between birth and age one. As linear mixed models were specified in a different manner than multivariable linear models used to investigate epigenome-wide DNAm changes, we first conducted a sensitivity analysis to ensure linear mixed model results were consistent with those from multivariable linear models used in our epigenome-wide analysis and could therefore be used to investigate persistence of cord blood DNAm changes. Specifically, we compared whether effect sizes returned from linear mixed models and multivariable linear models were in the same direction and of similar magnitude. Linear mixed model effect size magnitudes were consistently slightly smaller than those returned from multivariable linear models (**Supplementary File 12 and 14**). We hypothesize the decrease in effect size magnitude displayed in linear mixed models compared to multivariable linear models is likely due to differences in how we accounted for variation related to estimated cell type proportions and technical effects between the two models. Importantly, we noted that effect sizes always (except for 1 CpG) exhibited the same directionality between linear mixed models and multivariable linear models (**Supplementary File 12 and 14**). Based on these findings, we proceeded analyzing the persistence of cord blood DNAm changes using linear mixed models. When examining the persistence of DNAm changes at the top 100 individual CpGs identified in our epigenome-wide analysis of prenatal NO_2_ exposure, we found that two CpGs (cg16118259, cg18063278) displaying lower DNAm at birth and two CpGs displaying higher DNAm at birth (cg05686950, cg08033031) exhibited significant decreases in the absolute effect size magnitude of prenatal NO_2_ exposure between birth and age one (**Figure 4; Supplementary File 14**). In other words, we found that the absolute effect size magnitude of prenatal NO_2_ exposure at these four CpGs was closer to zero at age one than at birth. All four of these CpGs exhibiting an absolute effect size magnitude of prenatal NO_2_ exposure >0.025% DNAm at birth, but not at age one, suggesting the effect of prenatal NO_2_ exposure did not persist at these loci (**Figure 4; Supplementary File 14**). The remaining 96 CpGs from our epigenome-wide analysis exhibited nonsignificant changes in the absolute effect size magnitude of prenatal NO_2_ exposure between birth and age one. Of these 96 CpGs, 54 exhibited lower DNAm at birth and 42 displayed higher DNAm at birth. Almost all (N=50) CpGs exhibiting lower DNAm displayed a decrease in the absolute effect size of prenatal NO_2_ exposure between birth and age one, with the remaining minority (N=4) displaying an increase absolute effect size magnitude between birth and age one. Similarly, most (N=38) CpGs exhibiting higher DNAm at birth also displayed a decrease in the absolute effect size magnitude of prenatal NO_2_ exposure between birth and age one. We noted that 43 of the 54 CpGs exhibiting lower DNAm at birth and 32 of the 42 CpGs exhibiting higher DNAm at birth met the biological cut-off of an absolute effect size magnitude of prenatal NO_2_ exposure >0.025% DNAm at the birth time point. Only 21 CpGs displaying lower DNAm at birth and 11 CpGs displaying higher DNAm at birth continued to meet our biological cut-off of >2.5% DNAm at age one, suggesting the effect of prenatal NO_2_ exposure persisted at these loci (**Figure 4; Supplementary File 14**). Of the 151 CpGs contained within cord blood DMRs, 55 exhibited lower DNAm at birth while the remaining 96 displayed higher DNAm at birth (**Figure 5, Supplementary File 12**). A total of 38 CpGs exhibiting lower DNAm at birth exhibited a decrease in the absolute effect size magnitude of prenatal NO_2_ exposure between birth and age one. Similarly, the absolute effect size magnitude of prenatal NO_2_ exposure decreased between birth and age one at 72 of the 96 DMR CpGs exhibiting higher DNAm at birth. These changes failed to reach statistical significance at all 151 DMR CpGs. We noted that 48 of the 55 DMR CpGs exhibiting lower DNAm at birth and 71 of 96 DMR CpGs displaying higher DNAm at birth met the biological cut-off of an absolute effect size magnitude of prenatal NO_2_ exposure >0.025% DNAm at the birth time point. In contrast to 100 CpGs selected from our epigenome-wide analysis, almost all DMR CpGs meeting biological cut-offs at the birth time continued to meet biological cut-offs at age one. Specifically, 41 of the 48 DMR CpGs exhibiting lower DNAm at birth and 56 of the 71 DMR CpGs exhibiting higher DNAm at birth displayed an absolute effect size magnitude of prenatal NO_2_ exposure >0.025% DNAm at birth and age one, suggesting the effect of prenatal NO_2_ exposure persists at these loci until at least age one. Investigating effect size changes across individual CpGs located within DMRs does not clarify which DMRs are persistently or transiently affected by prenatal NO_2_ exposure. Therefore, we also examined which DMRs continued to display a mean absolute effect size magnitude of prenatal NO_2_ exposure >2.5% DNAm at age one. We noted that 23 of 24 cord blood DMRs displayed a mean absolute effect size magnitude of prenatal NO_2_ exposure >0.025% at birth based on linear mixed models (**Supplementary Table 2**). At age one, 19 DMRs continued to exhibit mean absolute effect size magnitude of prenatal NO_2_ exposure >2.5% DNAm (**Supplementary Table 2**). The DMRs exhibiting persistent effects of prenatal NO_2_ exposure included those annotated to developmentally relevant *HOX* genes (**Supplementary Table 2**).

**Figure 4.**
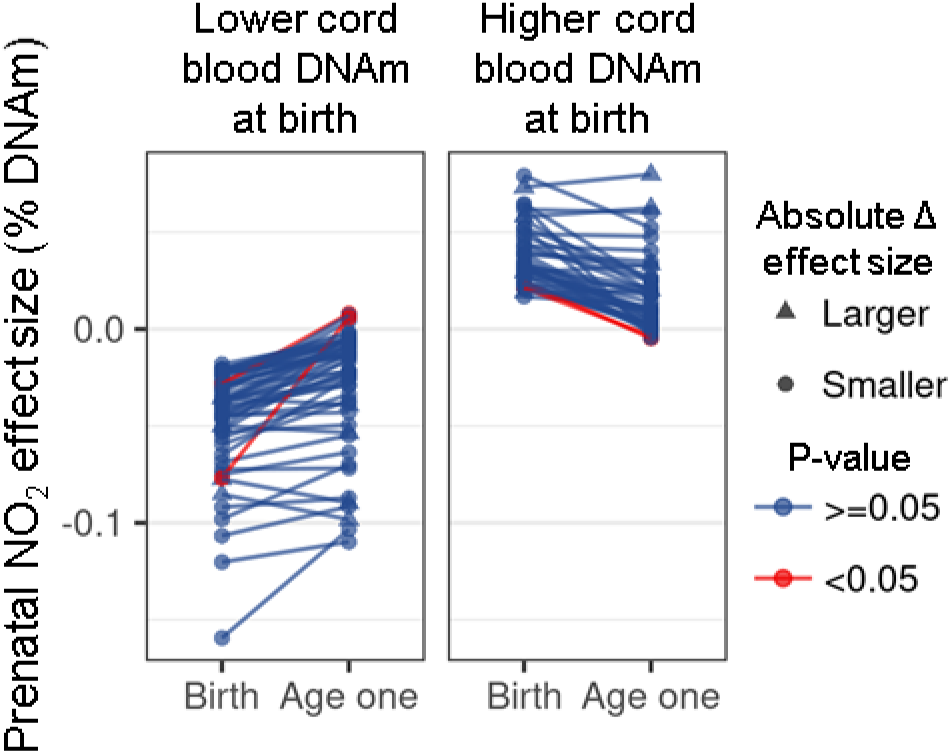
The effect size of prenatal NO_2_ exposure at the top 100 cord blood CpGs identified in an epigenome-wide analysis did not significantly differ between birth and age one in CHILD participants (N=124). The CpGs that exhibited lower DNAm at birth are plotted separately from those that displayed higher DNAm at birth to display the change in absolute effect size magnitude of prenatal NO_2_ exposure between birth and age one more readily. We observed that the absolute effect size magnitude of prenatal NO_2_ exposure decreased between birth and age one at 92 CpGs, with four of these CpGs exhibiting a statistically significant decrease. The remaining 8 CpGs exhibited non-significant increases in the absolute effect size magnitude of prenatal NO_2_ exposure between birth and age one. The CpGs exhibiting a smaller absolute effect size magnitude of prenatal NO_2_ exposure at age one are indicated by circles, while those displaying a larger absolute effect size magnitude of prenatal NO_2_ exposure at age one are indicated by triangles. The CpGs that displayed a significant (Bonferroni p<0.05) change in the absolute effect size of prenatal NO_2_ exposure between birth and age one are highlighted in red.

**Figure 5.**
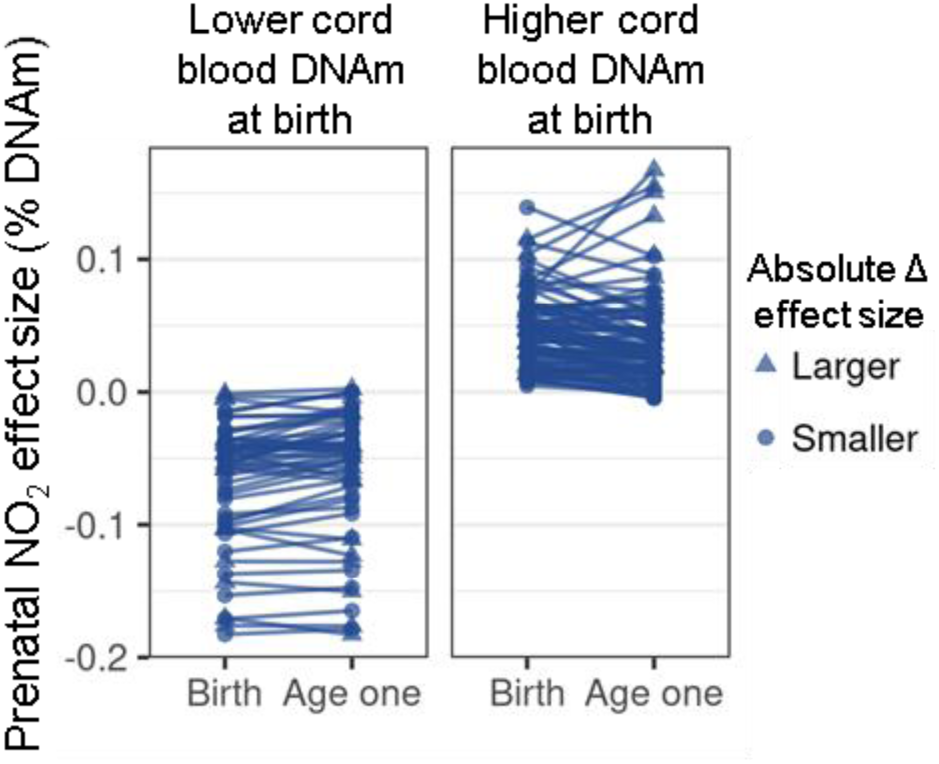
The effect size of prenatal NO_2_ exposure on CpG (N=151) DNAm within cord blood differentially methylated regions does not significantly differ between birth and age one in CHILD participants (N=124). The CpGs that exhibited lower DNAm at birth are plotted separately from those that displayed higher DNAm at birth to display the change in absolute effect size magnitude of prenatal NO_2_ exposure between birth and age one more readily. The 24 identified cord blood DMRs contained a total of 151 CpGs. Of these, 109 CpGs exhibited non-significant decreases in the absolute effect size magnitude of prenatal NO_2_ exposure between birth and age one. The remaining 42 CpGs exhibited non-significant increases in the absolute effect size magnitude of prenatal NO_2_ exposure at age one. The CpGs exhibiting a smaller absolute effect size magnitude of prenatal NO_2_ exposure at age one compared to birth are indicated by circles, and those displaying a larger absolute effect size of prenatal NO_2_ exposure at age one compared to birth are indicated by triangles.

Air pollution is hypothesized to reduce global DNAm by potentiating DNA demethylation and altering the availability of enzymes and methyl donors^48^. LINE DNAm can be used as a surrogate measure for whole genome DNAm and previous studies have shown LINE DNAm is affected by air pollution exposure^75^. Therefore, we investigated whether CHILD participants exhibited average genome-wide or LINE-specific DNAm alterations at birth or age one related to prenatal NO_2_ exposure or postnatal NO_2_ exposure, respectively. Genome-wide DNAm was measured by averaging DNAm across all CpGs for each individual. We obtained LINE-specific DNAm measures by averaging DNAm across all CpGs annotated to L1 LINEs on the Illumina HumanMethylation 450k array. We observed no significant associations between prenatal NO_2_ exposure and genome-wide DNAm alterations at birth (**Figure 6A**), nor did we find any correlation between year one NO_2_ exposure and genome-wide postnatal-specific DNAm changes (**Figure 6B**). Further, neither prenatal NO_2_ exposure (**Figure 6A**) or year one NO_2_ exposure (**Figure 6B**) was associated with significant changes to prenatal or postnatal-specific L1 LINE DNAm, respectively.

**Figure 6.**
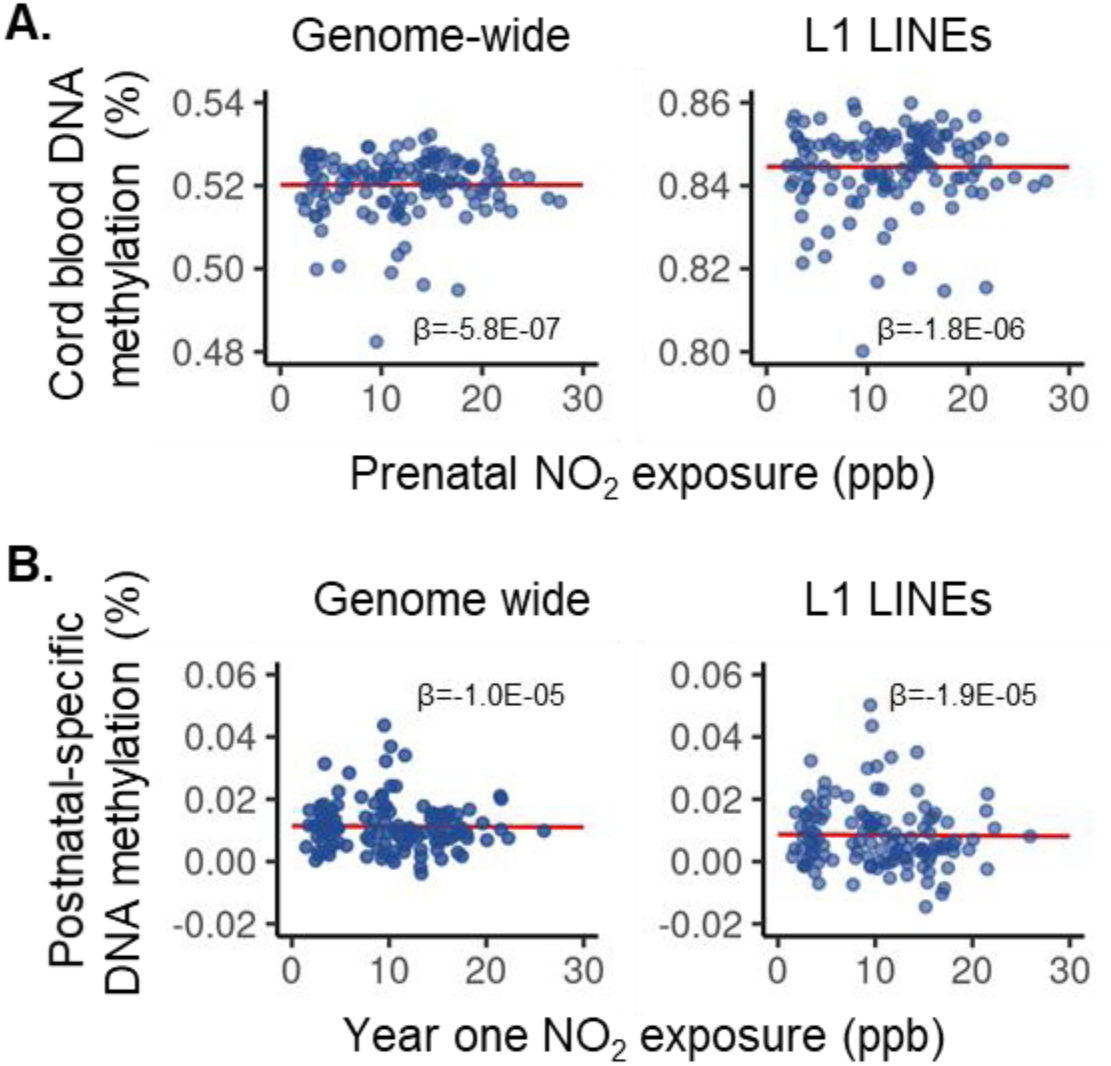
Prenatal and year one NO_2_ exposures are not associated with genome average or L1 LINE-specific measures of global DNA methylation in CHILD participants. (**A**) Associations between global measures of cord blood DNAm and prenatal NO_2_ exposure. Neither genome-wide DNAm (p=0.94; effect size=-1.5E-05; β=-5.8E-07) or LINE-specific DNAm (p=0.89; effect size=-4.7E-05; β=-1.8E-06) was significantly associated with prenatal NO2 exposure. (**B**) Associations between global measures of postnatal-specific DNAm and year one NO_2_ exposure. Neither postnatal-specific changes in genome-wide DNAm (p=0.25; effect size=-4.6E-4; β=-1.0e-05) or LINE-specific DNAm (p=0.33; effect size=-2.5E-4; β=-1.0e-05) were significantly associated with year one NO_2_ exposure. Red lines represent the estimated marginal effect of NO_2_ exposure on DNAm with a slope equal to coefficient of NO_2_ exposure (β). The surrounding shaded area represents the 95% confidence interval of the marginal effect.

Currently, researchers believe the biological embedding of early life exposures in DNAm patterns contributes to altered health outcomes. Using MANOVA tests we examined the relationship between DNAm across cord blood and postnatal-specific DMRs identified in this study with markers of childhood asthma, including age one atopy and wheeze. In cord blood, DNAm across Chr17:7486551-7486616 (adjusted p = 0.03) and Chr5:126409007-126409062 (adjusted p = 0.002) significantly differed between CHILD participants with (N=33) and without wheeze (N=86; **Figure 7A**). We also identified that DNAm across the Chr19:58879022-58879060 in cord blood was significantly (adjusted p=0.001) different between CHILD participants with (N=25) and without (N=103) atopy at age one (**Figure 7B**). We next examined whether the postnatal-specific DMR was associated with age one health outcomes. We observed that altered postnatal-specific DNAm across Chr17:76681111-46681402 was significantly (p=0.02) different between participants with (N=23) and without (N=102) age one atopy (**Figure 8**) but did not different between participants with (N=33) and without (N=87) wheeze at age one.

**Figure 7.**
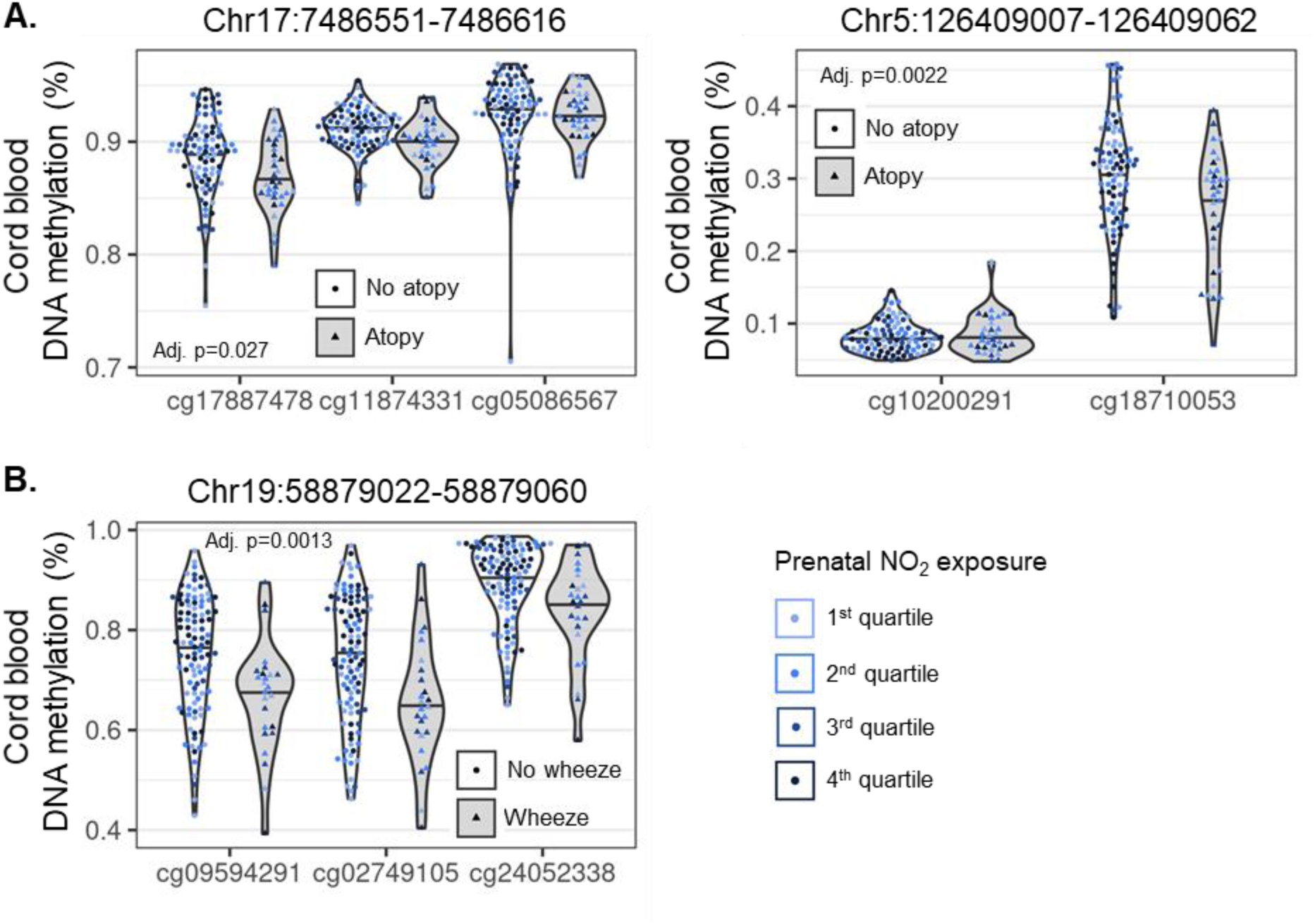
Three cord blood differentially methylated regions (DMRs) are associated with age one wheeze or age one atopy in CHILD participants. (**A**) Two cord blood DMRs are associated with age on wheeze in CHILD participants (N=119). MANOVA tests suggested that DNA methylation (DNAm) across Chr17:7486551-7486616 (adjusted p=0.03) and Chr5:126409007-126409062 (adjusted p=0.002) was significantly different between participants with (N=33) and without (N=86) wheeze at age one. (**B**) One cord blood DMR was associated with age on atopy in CHILD participants (N=128). MANOVA tests also suggests that DNAm across Chr19:58879022-58879060 (adjusted p=0.02) was significantly different between CHILD participants with and without atopy at age one. Shaded violin plots with triangular points represent DNAm measures for participants with wheeze (panel A) or atopy (panel B). Unshaded plots with circular points represent DNAm measures for participants without wheeze (panel A) or atopy (panel B). Points are coloured based on quartiles of prenatal NO_2_ exposure as specified in Supplementary Table 2 to display the relationship between prenatal NO_2_ exposure and altered DNAm most clearly.

**Figure 8.**
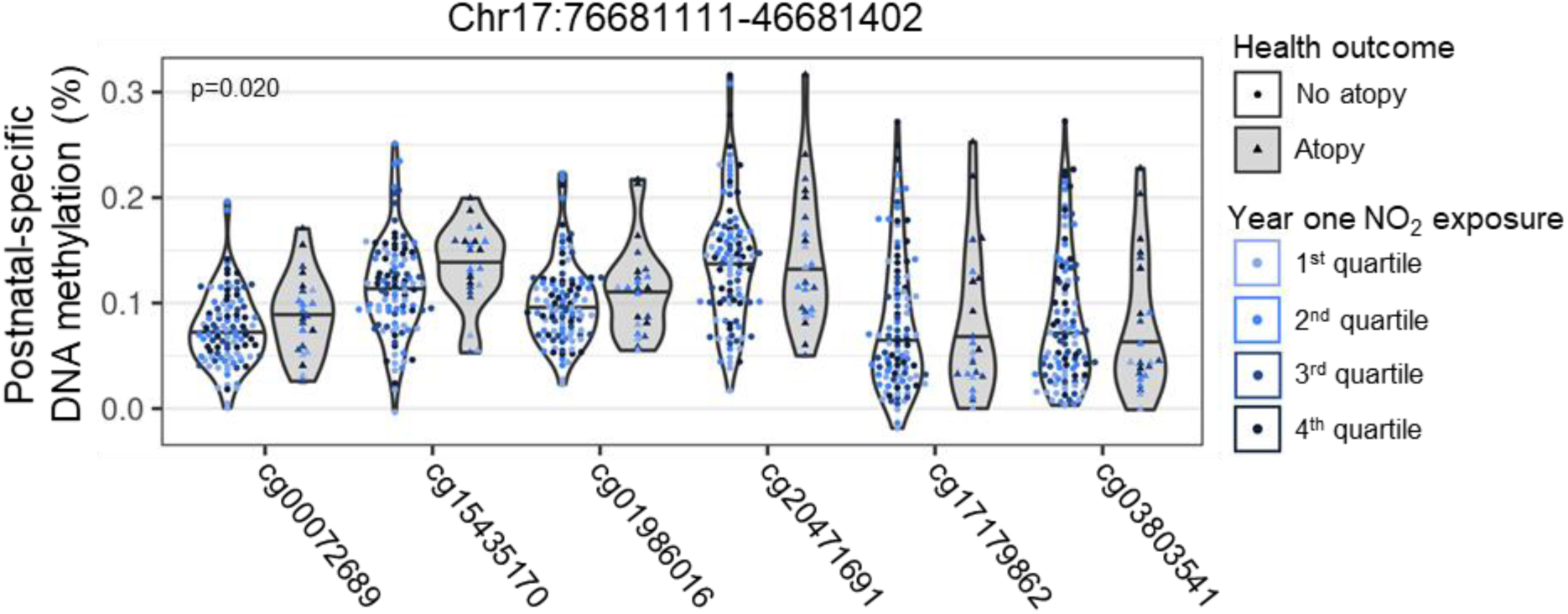
Altered postnatal-specific DNA methylation (DNAm) across Chr17:76681111-46681402 is associated with one age atopy in CHILD participants (N=125). MANOVA tests suggested that altered postnatal-specific DNAm across Chr17:76681111-46681402 significantly (p=0.020) differed between CHILD participants with (N=23) and without atopy (N=102). Shaded violin plots with triangular points represent DNAm measures for participants with atopy. Unshaded plots with circular points represent DNAm measures for participants without atopy. Points are coloured based on quartiles of year one NO_2_ exposure as specified in Supplementary Table 2 to display the relationship between prenatal NO_2_ exposure and altered DNAm most clearly.

Levels of ambient NO_2_ vary annually in Canada, with the highest levels typically occurring during colder periods^81^. We hypothesized that children exposed to higher levels of air pollution during critical developmental periods may experience larger magnitudes of DNAm alterations. Therefore, we conducted a sensitivity analysis to examine whether effect of NO_2_ exposure on DNAm varies by birth month at the top 10 CpGs identified in our prenatal and postnatal-specific epigenome-wide analyses. We found that modelling the effect of prenatal NO_2_ exposure on cord blood DNAm by birth month did not significantly improve model fit based on ANOVA likelihood ratio tests (**Supplementary Figure 38**). Similarly, ANOVA likelihood tests suggested that modelling the effect of year one NO_2_ exposure on postnatal-specific DNAm did not improve model fit (**Supplementary Figure 39**), suggesting the effects of prenatal and year one NO_2_ exposure on DNAm do not vary by birth month.

## Discussion

In the present study, we investigated the persistence of cord blood DNAm changes associated with prenatal NO_2_ exposure to identify epigenetic alterations with the greatest potential to influence child health outcome. Additionally, longitudinal measures of DNAm facilitated the investigation of postnatal-specific DNAm changes associated with NO_2_ exposure in the first year of life. Our results show that prenatal NO_2_ exposure is associated with regional DNAm changes, and that the effect of prenatal NO_2_ exposure across CpGs within these regions remains similar at age one. These findings suggest the effects of prenatal NO_2_ exposure on DNAm persist through early life and are therefore more likely to affect child respiratory health outcomes. This notion is consistent with our observations that several regional cord blood DMRs were also associated with wheeze or atopy at age one. Additionally, our observations that postnatal-specific DNAm alterations are associated with atopy contribute to our understanding that the postnatal period remains a sensitive window for the biological embedding of air pollution exposure. Together, these findings provide greater insight into the putative molecular pathways underlying the association between prenatal and early life NO_2_ exposure and childhood atopic disease.

We observed several cord blood DMRs localized to developmentally relevant genes. Additionally, linear mixed models showed that the effect of prenatal NO_2_ exposure on CpG DNAm contained within these DMRs did not significantly vary at age one, and that the majority of DMR CpGs meeting our biological cut-off of a mean absolute effect size of prenatal NO_2_ exposure at birth (N=109 CpGs) continued to meet this cut-off at age one (N=98 CpGs). In contrast, only 32 of 75 individual CpGs from our epigenome-wide analysis met the biologically cut-off for prenatal NO2 exposure effect size at birth and age one. Together, this implies the effect of prenatal NO2 exposure is more persistent across altered regions than individual CpGs. We observed persistently altered regions of DNAm in *HOXA4* and *HOXA5*, which are strongly implicated in lung development and also have roles in immune cell differentiation^85, 86^. Regional DNAm changes associated with air pollution exposure have previously been reported in *HOXA4* and *HOXA5*. For example, Gruzieva *et al.* (2019) reported a *HOXA4* DMR in cord blood associated with prenatal PM_10_ exposure^87^ and Chi *et al.*(2016) reported a *HOX5A* DMR in adult peripheral blood associated with PM_2.5_^76, 87^. Further, altered *HOX5A* DNAm reported by Chi *et al.*(2016) was associated with gene expression of *HOX5A*, as well as *HOXA9* and *HOXA10* gene expression, demonstrating that altered *HOX5A* DNAm has functional consequences^76^. Both *HOXA4* and *HOXA5* DMRs identified in this study overlap with those previously reported by Gruzieva *et al.* (2019) and Chi *et al.* (2016)^76, 87^. It is important to note that in this study we examined the effect of prenatal NO_2_ exposure while the aforementioned studies investigated particulate matter exposures. NO_2_ exposure is considered a putative causative agent of DNAm change as well as a surrogate measure of traffic-related air pollution, which also contains particulate matter^28^. While it is not possible to disentangle whether the changes we observed are due to the direct of effect of prenatal NO_2_ exposure or potentially other air pollutants NO_2_ may be correlated with, the similarities between DMRs identified in this study and those observed previously strongly implicate air pollution exposure in altered DNAm of HOX genes. Given the roles HOX genes play in prenatal and early life lung development, persistently altered HOX gene DNAm observed in this study may represent a molecular mechanism linking prenatal air pollution exposure to changes in child lung structure observed in asthma^88^.

Most studies investigating the impact of early-life NO_2_ exposure have focused on the prenatal period based on the notion that disruption of DNAm patterns during epigenetic reprogramming, especially those related to immune function and lung development, will have the largest impact on cell function and long term respiratory health^89, 90^. The early postnatal period, especially the first year of life, is often overlooked despite the continued development of the lungs, immune system, and DNAm patterns during this time^14, 15, 91^. In this study we observed that year one NO_2_ exposure was associated with a postnatal-specific DMR annotated to *HOXB6*. Further, we identified that altered DNAm across this region was associated with age one atopy, suggesting that this DNAm change may underlying altered child health outcomes. This is consistent with results from a large cross-sectional epidemiological showing prenatal and postnatal passive cigarette smoke exposures independently affect the risk of negative respiratory health outcomes in childhood^17^. Together, these findings support the first year of life as also a critical window for the biological embedding of air pollution exposure. Of note, we also observed a cord blood DMR localizing to *HOXB6*, although at an upstream location, suggesting *HOX6B* DNAm appears to be sensitive to air pollution exposure throughout early-life, and that prenatal and postnatal-specific changes in DNAm may play different roles in the regulation of *HOXB6* expression^92^. The *HOXB6* gene is one of the most highly expressed HOX family members in human fetal and adult lung, playing an important role in early lung development, and also in hematopoiesis^88^. Thus, prenatal and postnatal *HOXB6* DNAm changes may work together to affect lung development across early-life, leading to long-term changes in lung structure and/or function that contribute to the pathology of childhood asthma^93^. Alternatively, changes to *HOXB6* DNAm in hematopoietic stem cells could potentiate the expansion of myeloid precursor cells and their progenitors, leading to changes in cell abundances (such as increased mast cells) observed in asthma^94^. In this study, we investigated DNAm changes under the hypothesis of cellular reprogramming^7^. Additional investigation is needed to determine if *HOXB6* DNAm is affected in a similar manner within other tissues and how the DNAm alterations we observed affect cell function and differentiation.

In CHILD study participants, concurrent atopy and wheeze at age one is a stronger predictor of age three asthma than either condition occurring alone^3^. We were limited in our ability to examine associations between DNAm across significant DMRs and atopy and wheeze at age one together due to a small number (N=9 in our analysis of prenatal NO_2_ exposure) of participants in our analyses displaying both conditions at age one. Instead, we used MANOVA tests to separately examine the association between DNAm across significant cord blood and postnatal-specific DMRs and age one wheeze or age one atopy. We observed two cord blood DMRs annotated to *MPDU1* and *C5orf63* (*FLJ44606*) that were associated with age one wheeze. The association between *MPDU1* DNAm and age one wheeze represents a novel observation. *MPDU1* codes for an endoplasm reticulum protein that utilizes mannose-P-dichol in the synthesis of lipid-linked oligosaccharides and glycosylphosphatidylinositols, which are used downstream in post-translational modification of proteins^95^. It remains possible that alterations in protein post-translational modifications arising from changes in *MPDU1* DNAm may contribute to childhood wheeze and asthma. For example, glycosylation of the Fc region of IgG1 can impact antibody function and decreases in IgG1 glycosylation are associated with immune activation and have been observed in asthmatic compared to non-asthmatic children, as well as in murine models of allergic asthma^96–98^. Two previous studies have reported DMRs in *C5orf63* associated with diesel exhaust inhalation in bronchial epithelial cells and with smoking in peripheral blood samples obtained from a Korean cohort^99, 100^. The *C5orf63* DMR observed in this study overlaps with both previously identified *C5orf63* DMRs, implying this DMR represents a robust DNAm change across development stages, tissues, and human populations in response to inhaled pollutant exposure. One of the mechanisms by which inhaled air pollutants induce DNAm changes is through oxidative stress. Given that regional *C5orf63* DNAm changes are robust across studies and that *C5orf63* encodes a glutaredoxin-like protein, these findings support a role for *C5orf63* in oxidative stress following inhaled air pollutant exposure that is regulated by DNAm^101^. To our knowledge, this is the first study to identify an association between *C5orf63* DNAm and childhood wheeze. Increased oxidative stress contributes to the underlying pathology of asthma^102^. The observed changes in *C5orf63* DNAm may represent an underlying molecular mechanism of this pathology. In cord blood, we also observed that altered DNAm in *ZNF837* was associated with age one atopy. The *ZNF837* protein is predicted to help regulate nuclear transcription by RNA polymerase II. There exists a previous report of transcription factor binding sites for AHR and ARNT in the promoter of *ZNF387*^103, 104^. These proteins represent transcription factors which enter the nucleus and modulate gene expression following inhaled air pollution exposure, suggesting that the *ZNF837* product plays a role in this pathway^105^. Though the genes which the *ZNF837* protein regulates remain to be elucidated, we envision a role for *ZNF837* in regulating expression and DNA binding of atopy and asthma-relevant gene products and transcription factors, respectively^106^. Replication of these results followed by further experiments is needed to confirm the exact role *ZNF837* plays in gene expression following air pollution exposure.

This subset of CHILD study participants was over-sampled for atopic conditions including allergenic sensitization to inhaled or food allergens. Atopy often occurs together with asthma and atopic dermatitis in what is known as the “atopic triad”^107^. There is an underlying genetic component to the atopic triad; however, early life environmental exposures such as air pollution exposure increase disease predisposition^107^. It is possible that genetic variants that increase atopy risk may also increase the susceptibility of nearby CpGs to DNAm changes following air pollution exposure, known as methylation trait quantitative loci (mQTLs)^71^. Our small sample size limited our ability to investigate the role of mQTLs in the DNAm changes we observed. When we cross checked cord blood DMR CpGs identified in this study with an online database constructed using data from the European ARIES cohort^72^, we found that 55 of these cord blood DMR CpGs represented potential mQTLs with effect sizes ranging from 0.02 to 0.27. Interestingly, no CpGs contained within the cord blood DMRs associated with wheeze or atopy were identified as potential mQTLs, suggesting that DNAm within these regions plays a primary role in determining child respiratory outcomes. Investigation of mQTLs in future studies will provide insight to which populations may be most susceptible to DNAm changes and adverse health outcomes following early-life air pollution exposure.

The longitudinal design of the CHILD study is a major advantage in investigating persistent DNAm changes as we were able follow DNAm within the same group of individuals through the first year of life. Additionally, this study design allowed us to isolate and examine postnatal-specific DNAm alterations associated with NO_2_ exposure. To our knowledge, this is the first time postnatal-specific DNAm changes have been investigated in the context of early-life air pollution exposure. The biological context of our findings in relation to childhood atopic disease is limited by the sample type (blood mononuclear cells) used. White blood cells are of interest as changes to immune cell function are thought to play a role in the underlying pathology of child atopic disease^108^. While some DNAm patterns in blood appear to be surrogate markers for less accessible tissues, such as brain^109^, the correlation between DNAm in blood and DNAm in tissues related to respiratory health such as lung remains unclear. Thus, it remains to be determined whether the DNAm changes we identified also occur in respiratory tissue following early life NO_2_ exposure. This investigation was also limited by the relatively small sample size of CHILD participants and a relatively low variation in NO_2_ compared to previous studies^8^, likely decreasing our ability to detect additional DNAm alterations. Finally, the CHILD cohort is mainly comprised of white participants with higher socioeconomic status. Replication of our results in larger cohorts with more diverse populations is required before they can be generalized.

## Summary

Together, these findings suggest prenatal exposure to NO_2_ air pollution alters regions of DNAm and that DNAm changes across most of these regions persist into early childhood. For the first time we show that the first year of life remains a sensitive period to NO_2_ exposure, resulting in postnatal-specific DNAm alterations. Our observations provide further support these changes are occurring in cellular pathways related to early lung development. This knowledge contributes to our understanding of the molecular pathways that may be altered by early life air pollution exposure and affect child health outcome.

## Supporting information

Supplementary tables excluding 10 and 11

## Acknowledgements

This study was supported by a fellowship from the Canadian Association of Asthma, Allergy, and Immunology Fellowship (S.L.). We thank the CHILD Cohort Study participant families for their dedication and commitment to advancing health research. CHILD was initially funded by CIHR and AllerGen NCE Inc.

## Code availability

Code used to conduct the analyses reported in this study can be accessed at https://github.com/mjoneslab/CHILD_prenatal_no2_analysis.

---

**Supplementary Figure 1.**
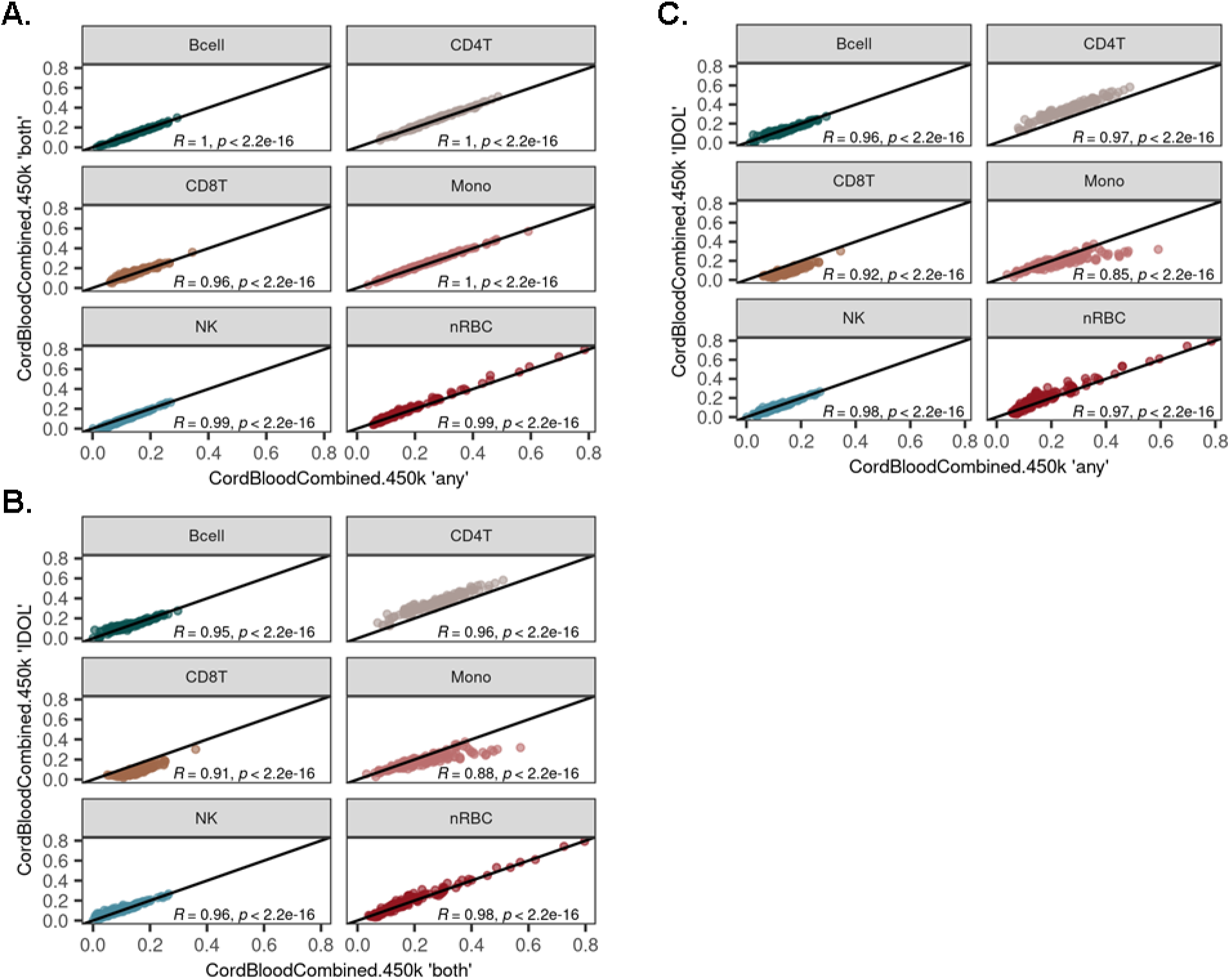
Umbilical cord white blood cell deconvolution methods perform similarly in estimating monocyte cell proportions. (**A**) Deconvolution of umbilical cord white blood cells with the FlowSorted.CordBloodCombined.450k package using “any”, which selects the top100 probes with the greatest magnitude of difference between cell types, or “both”, which selects equal numbers of probes with the greatest magnitude of effect in either the positive or negative direction. (**B**) Deconvolution of umbilical cord white blood cells with the FlowSorted.CordBloodCombined.450k package using “any” of the top100 probes, or “IDOL”, which using a pre-determined set of optimized probes for deconvolution of cord blood white cells. (**C**) Deconvolution of umbilical cord white blood cells with the FlowSorted.CordBloodCombined.450k package using “both” to select the equal numbers of probes with the greatest magnitude of effect in the positive (50) or negative (50) direction, or using optimized “IDOL” probes.

**Supplementary Figure 2.**
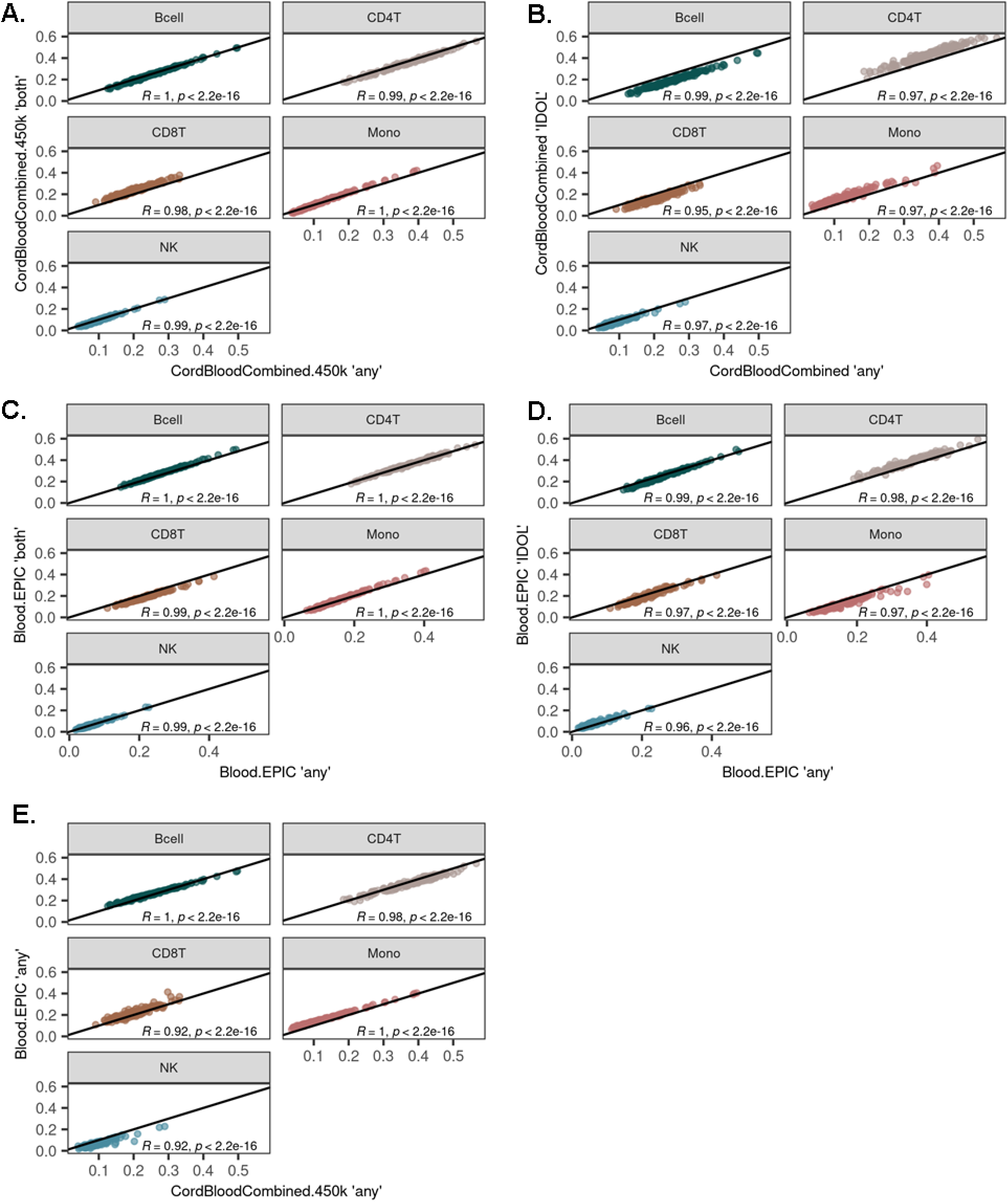
Age one one peripheral white blood cell deconvolution methods perform similarly in estimating monocyte cell type proportions. (**A**) Deconvolution of age one peripheral white blood cells using the FlowSorted.CordBloodCombined.450k package (cord blood references set) using “any” probe select, which returns the top100 probes with the greatest magnitude of difference between cell types, or “both” probe select, which returns equal numbers of probes with the greatest magnitude of effect in either the positive or negative direction. (**B**) Deconvolution of age one peripheral white blood cells with the FlowSorted.CordBloodCombined.450k package selecting “any” of the top100 probes, or selecting cord blood “IDOL” probes, which comprises a set of predetermined probes optimized for deconvolution of cord blood white cells. (**C**) Deconvolution of age one peripheral white blood cells with the FlowSorted.Blood.EPIC package (adult peripheral blood reference set) and selecting “any” probes, or “both” probes. (**D**) Deconvolution of age one peripheral white blood cells with the FlowSorted.Blood.EPIC package and selecting “any” probes, or using adult “IDOL” probes, which comprises a set of predetermined probes optimized for deconvolution of adult peripheral white blood cells. (**E**) Deconvolution of age one peripheral white blood cells using the FlowSorted.CordBloodCombined.450k package using “any” probe select, or deconvolution using the FlowSorted.Blood.EPIC package and selecting “any” probes.

**Supplementary Figure 3.**
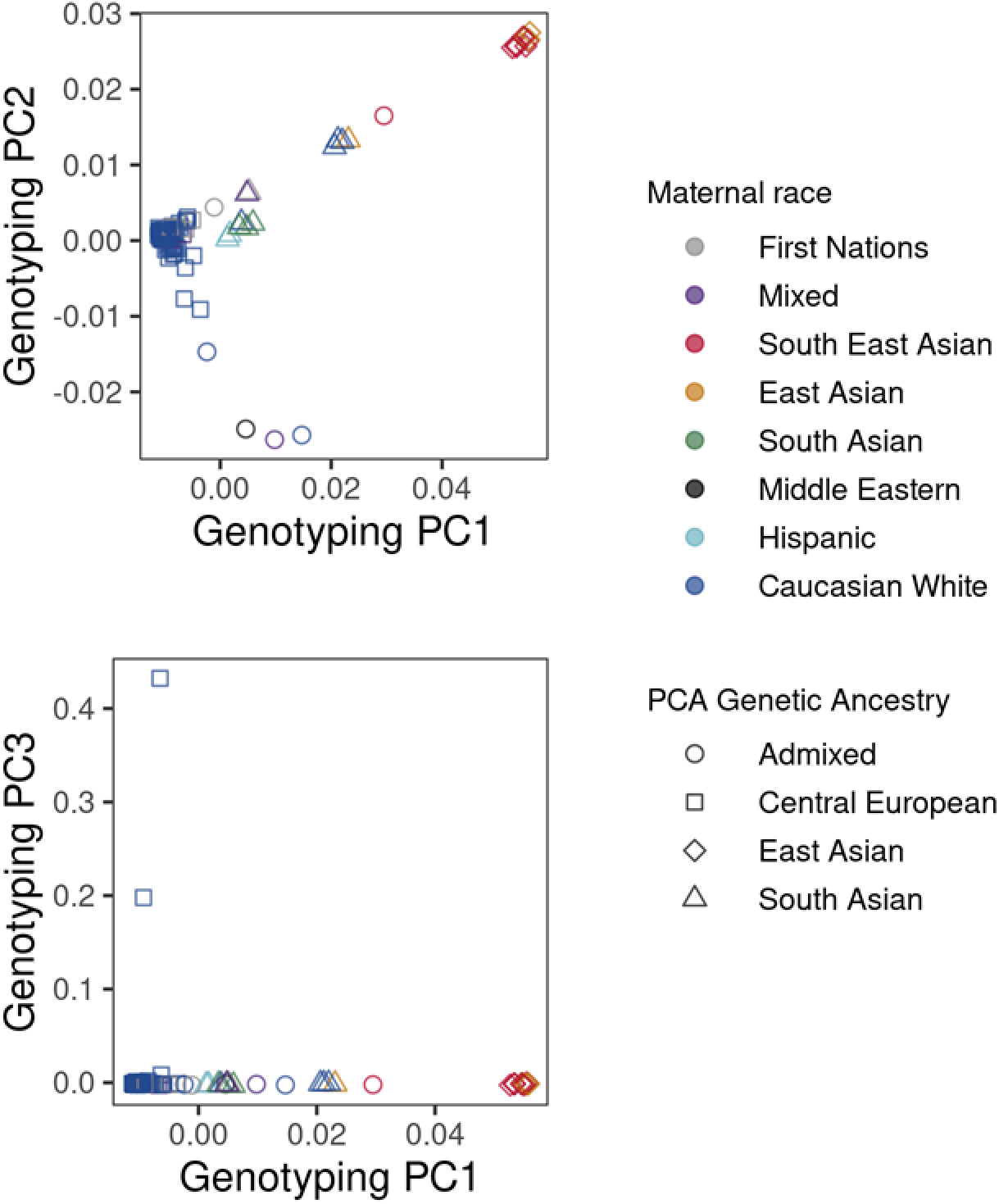
The first three genetic ancestry principle components (PCs) separate CHILD participants (N=131) by self-identified race. Point colour represents maternal-self identified race and point shape represents genetic ancestry as determined by principle component analysis.

**Supplementary Figure 4.**
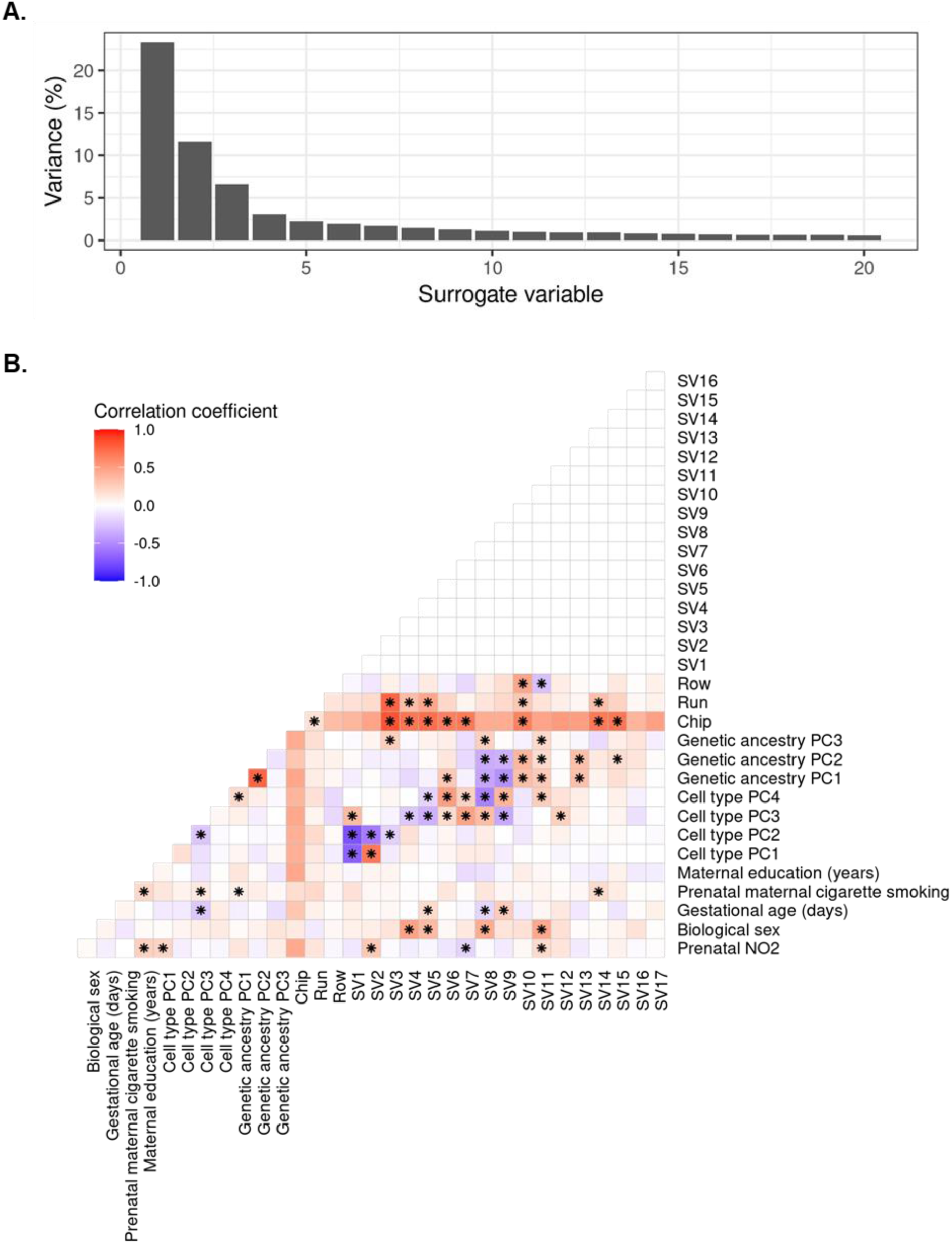
Surrogate variable analysis (SVA) captures technical variation in cord blood DNA methylation (N=128). (**A**) Variation captured by the first 20 surrogate variables. Surrogate variables capture variation that is not accounted for by covariates specified in the model of prenatal NO_2_ exposure. The first 17 surrogate variables were included in the prenatal model of NO_2_ exposure based on recommendation of the SVA package. (**B**) Correlations between covariates in the prenatal NO_2_ model and the first 17 surrogate variables returned by SVA. Note that the surrogate variables are strongly correlated with chip and run. *Represent a significant correlation between two variables.

**Supplementary Figure 5.**
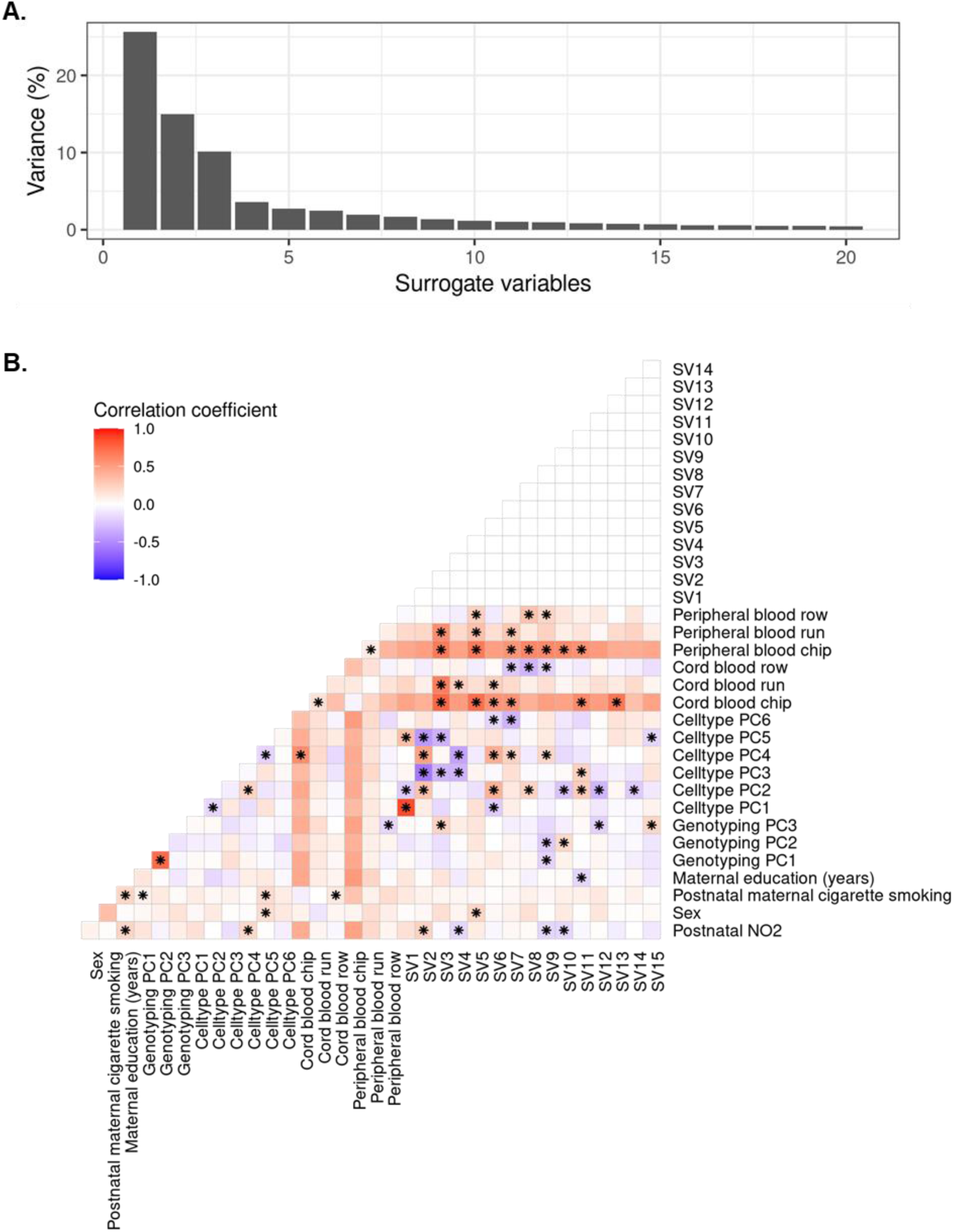
Surrogate variable analysis (SVA) captures technical variation in postnatal-specific DNA methylation (N=125). (**A**) Variation captured by the first 20 surrogate variables. Surrogate variables capture variation that is not accounted for by covariates specified in the model examining postnatal-specific DNA methylation changes associated with year one NO_2_ exposure. The first 15 surrogate variables were included in models examining year one NO_2_ exposure and postnatal-specific DNA methylation. (**B**) Correlations between covariates included in models examining postnatal-specific DNA methylation changes associated with year one NO2 exposure and the first 15 surrogate variables returned by SVA. Note that surrogate variables are strongly and correlated with chip and run. *Represent a significant correlation between two variables.

**Supplementary Figure 6.**
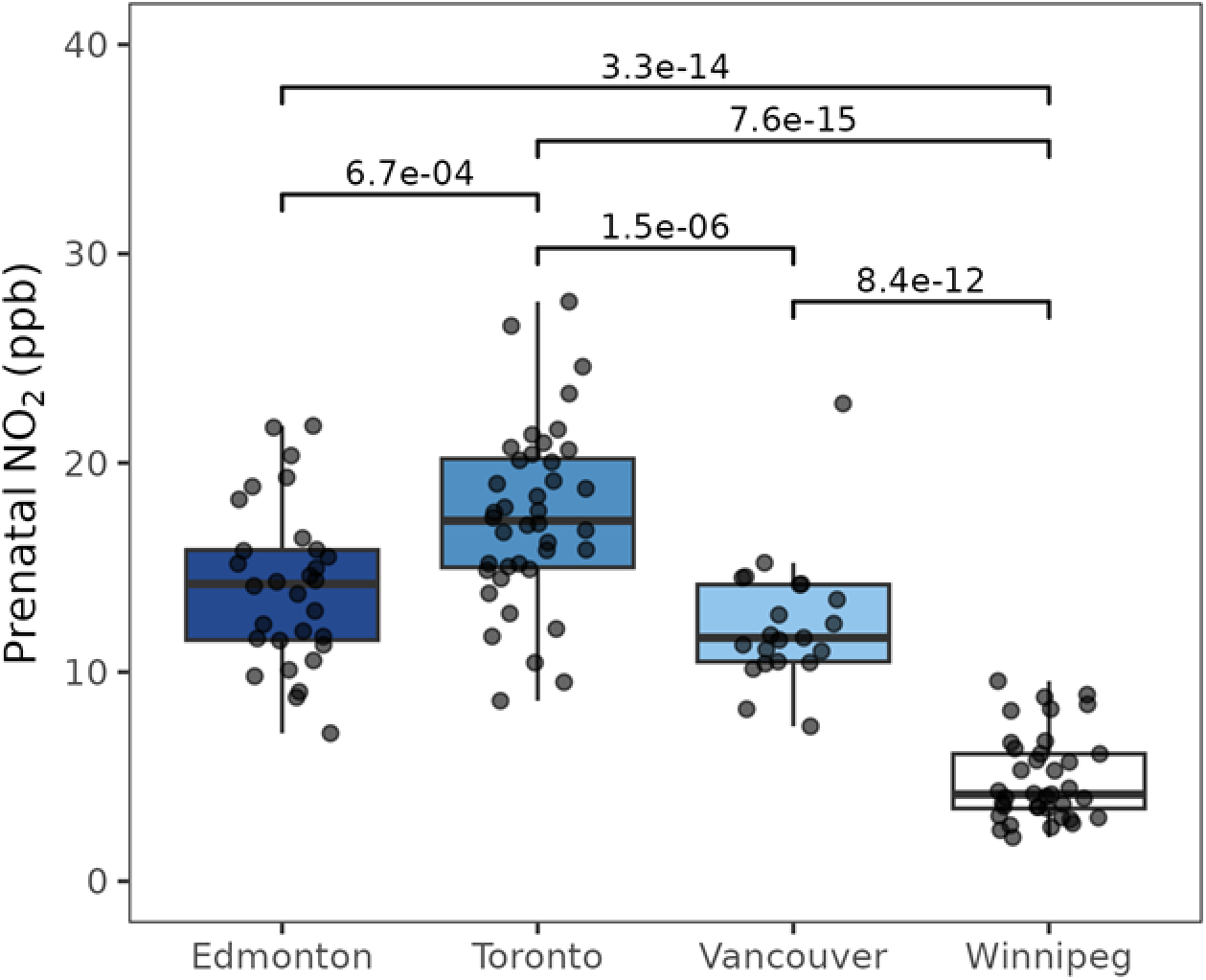
Variation in prenatal NO_2_ exposure estimates is greater between study sites compared within study sites for participants included in analysis of prenatal DNA methylation changes (N=128). The four CHILD study sites are located in the Canadian cities of Edmonton, Toronto, Vancouver, and Winnipeg. An ANOVA test indicated that prenatal NO_2_ exposure estimates were significantly (p = 1.0e-different between study sites. Post-hoc Tukey tests using the Honest Significant Difference method revealed that mean prenatal NO_2_ exposure estimates were significantly (adjusted p < 0.05) different between all study sites except for Edmonton and Toronto. The mean NO_2_ exposure estimate across all study sites was 12.2 ± 6.1 ppb. The mean and standard deviation of NO_2_ exposure estimates for Edmonton, Toronto, Vancouver, and Winnipeg were 14.1 ± 3.8 ppb, 17.4 ± 4.3 ppb, 12.4 ± 3.2 ppb, and 4.9 ± 2.1 ppb, respectively.

**Supplementary Figure 7.**
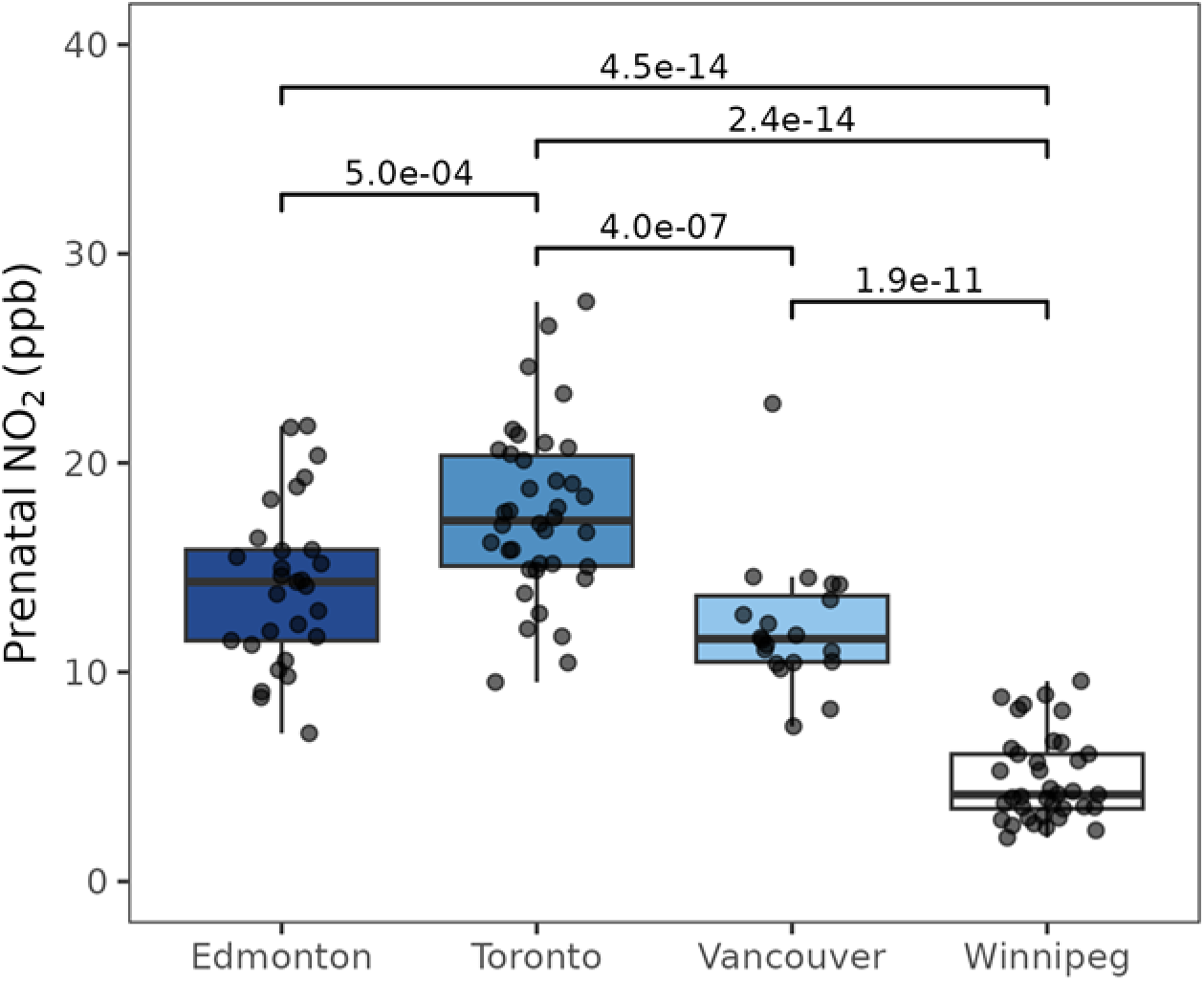
Variation in prenatal NO_2_ exposure estimates is greater between study sites compared to within study sites for participants included in analysis of persistent DNA methylation changes (N=124). The four CHILD study sites are located in the Canadian cities of Edmonton, Toronto, Vancouver, and Winnipeg. An ANOVA test indicated that prenatal NO_2_ exposure estimates were significantly (p = 4.5e-different between study sites. Post-hoc Tukey tests using the Honest Significant Difference method revealed that mean prenatal NO_2_ exposure estimates were significantly (adjusted p < 0.05) different between all study sites except for Edmonton and Toronto. The mean NO_2_ exposure estimate across all study sites was 12.2 ± 6.1 ppb. The mean and standard deviation of NO_2_ exposure estimates for Edmonton, Toronto, Vancouver, and Winnipeg were 14.3 ± 3.9 ppb, 17.6 ± 4.1 ppb, 12.2 ± 3.2 ppb, and 4.9 ± 2.1 ppb, respectively.

**Supplementary Figure 8.**
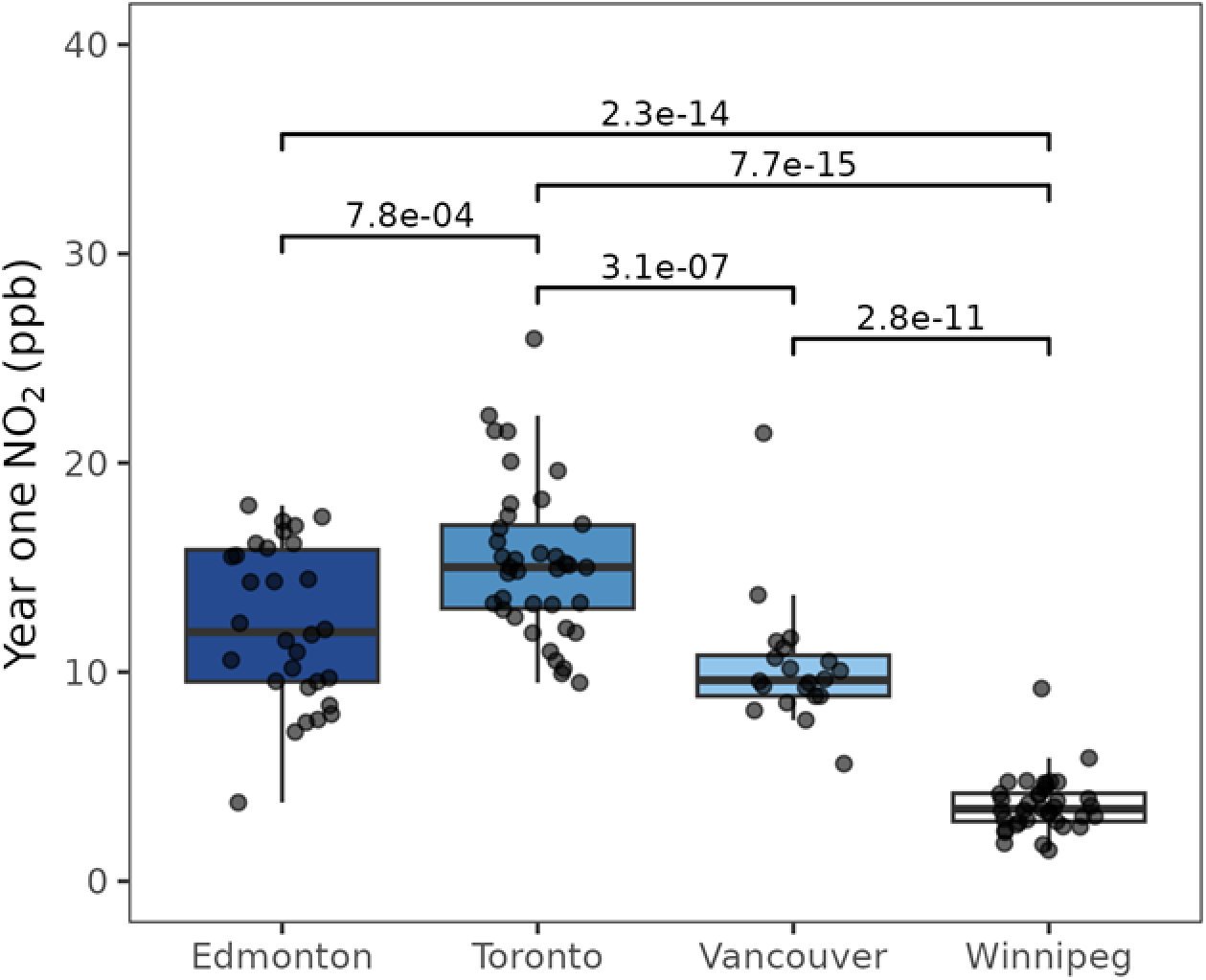
Variation in year one NO_2_ exposure estimates is greater between study sites compared to within study sites for participants included in analysis of postnatal-specific DNA methylation changes (N=124). The four CHILD study sites are located in the Canadian cities of Edmonton, Toronto, Vancouver, and Winnipeg. An ANOVA test indicated that year one NO_2_ exposure estimates were significantly (p = 3.1e-31) different between study sites. Post-hoc Tukey tests using the Honest Significant Difference method revealed that mean prenatal NO_2_ exposure estimates were significantly (adjusted p < 0.05) different between all study sites except for Edmonton and Toronto. The mean NO_2_ exposure estimate across all study sites was 10.3 ± 5.6 ppb. The mean and standard deviation of NO_2_ exposure estimates for Edmonton, Toronto, Vancouver, and Winnipeg were 12.3 ± 3.8 ppb, 15.3 ± 3.7 ppb, 10.3 ± 3.2 ppb, and 3.7 ± 1.4 ppb, respectively.

**Supplementary Figure 9.**
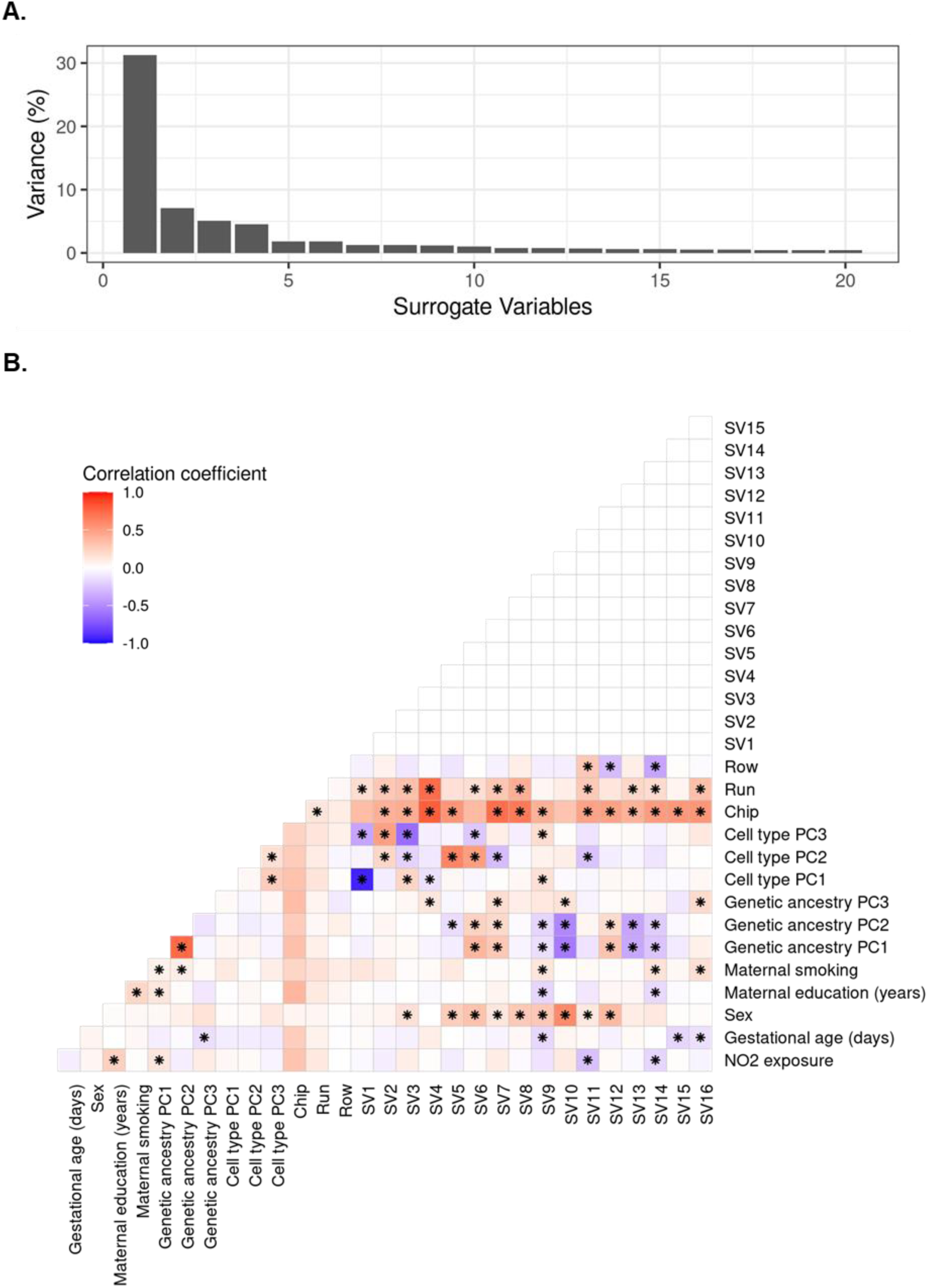
Surrogate variable analysis (SVA) captures technical variation associated across cord blood and peripheral blood DNA methylation used in the investigation of DNA methylation persistence (N=124). (**A**) Variation captured by the first 20 surrogate variables. Surrogate variables capture variation that is not accounted for by covariates specified in the model. The SVA package suggested inclusion of the first 96 surrogate variables, which would significantly reduce the power available to detect DNA methylation changes. Therefore, we included the first 16 surrogate variables in linear mixed models examining the persistence of DNA methylation changes as these surrogate variables accounted for ∼60% of the captured variance, which was similar to variance captured by the first 17 cord blood surrogate variables (∼60%) and the first 15 postnatal-specific surrogate variables (∼69%). (**B**) Correlations between covariates included in linear mixed models investigating the persistence of cord blood DNA methylation changes and the first 16 surrogate variables returned by SVA. Note that the surrogate variables are strongly and positively correlated with chip and run. *Represent a significant correlation between two variables.

**Supplementary Figure 10.**
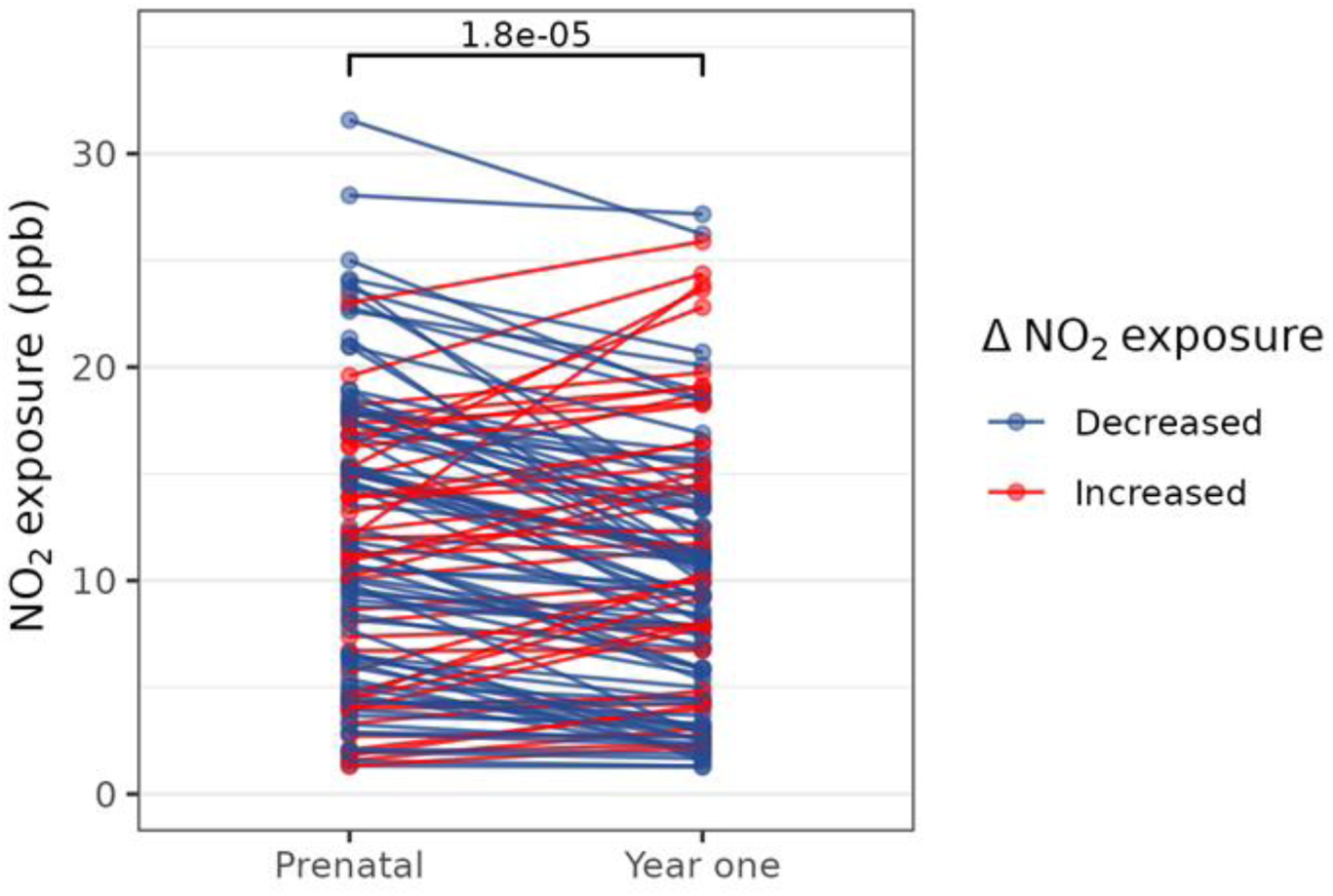
Year one NO_2_ exposure estimates was slightly lower than prenatal NO_2_ exposure estimates for CHILD participants. Paired t-tests indicated this difference was significant (p=1.8e-05) for participants (N=122) that had available both prenatal and year one NO_2_ exposure estimates. For the majority of participants (N=82; shown in blue) year one NO_2_ exposure estimates were lower than prenatal NO_2_ exposure estimates, while year one NO_2_ exposure estimates were higher than prenatal NO_2_ exposure estimates for the remaining participants (N=40; shown in red).

**Supplementary Figure 11.**
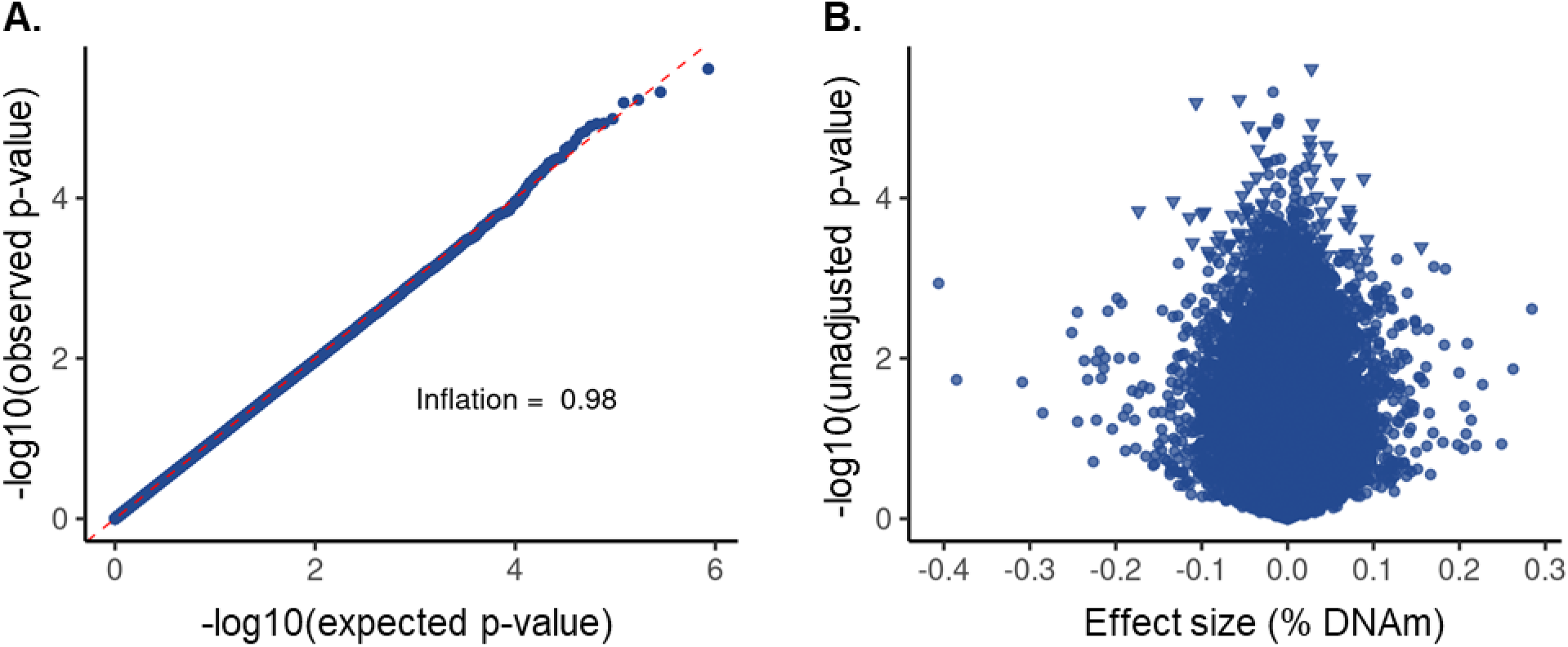
No individual CpGs were significant in the epigenome-wide analysis of prenatal NO_2_ exposure in CHILD participants (N=128). (**A**) The qqplot did not exhibit significant inflation or deflation suggesting adequate model fit. (**B**) No p-values surpassed our biological cut-off of an absolute effect size of prenatal NO2 exposure >0.025% DNA methylation at a false discovery rate of 5%. Triangular points represent the top 100 CpGs used in our investigation of persistence of individual cord blood DNA methylation changes.

**Supplementary Figure 12.**
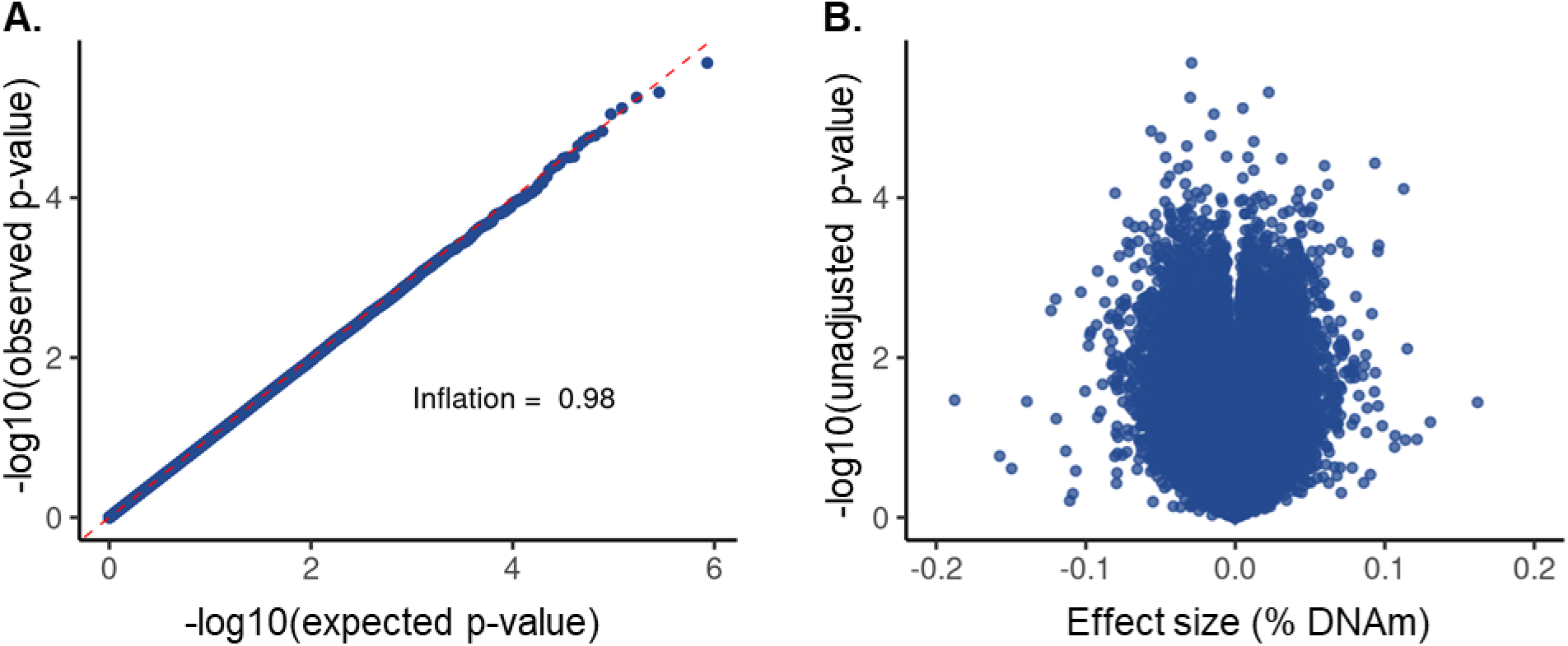
No individual CpGs were significant in the postnatal-specific epigenome-wide analysis year one NO_2_ exposure in CHILD participants (N=125). (**A**) The qqplot did not exhibit significant inflation or deflation suggesting adequate model fit. (**B**) No p-values surpassed our biological cut-off of an absolute effect size of year one NO_2_ exposure >0.025% DNA methylation at a false discovery rate of 5%. Triangular points represent the top 100 CpGs used in our investigation of persistence of individual cord blood DNA methylation changes.

**Supplementary Figure 13.**
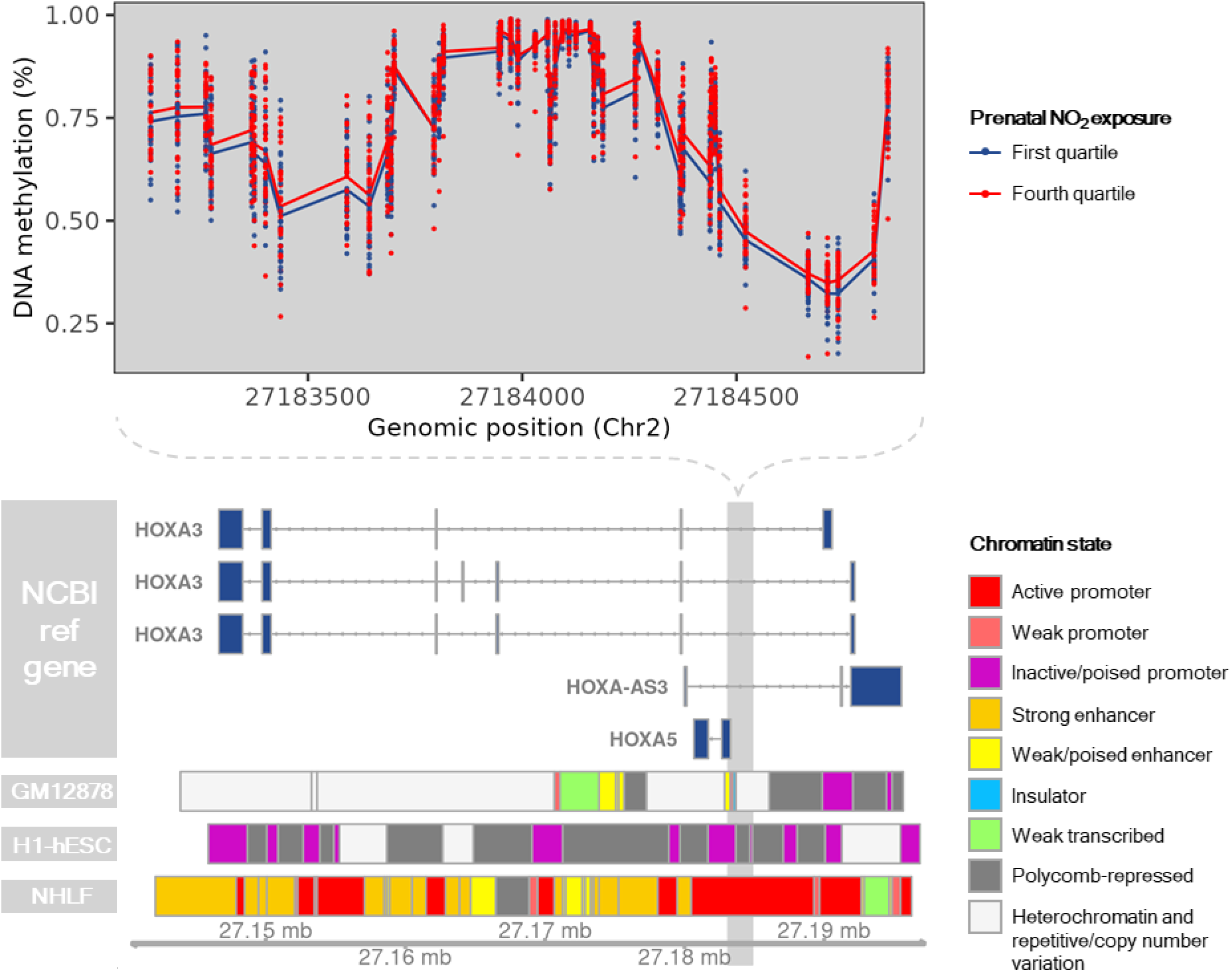
DNA methylation and chromatin state across the cord blood differentially methylated region (DMR) located on Chr2: 27183133-27184854. DNA methylation at CpGs contained within Chr2: 27183133-27184854 is shown for individuals with prenatal NO_2_ exposure < 7.3 ppb (first quartile; N=32; blue) or prenatal NO_2_ exposure > 16.3 ppb (fourth quartile; N=32; red) is displayed. Annotation of known genes and chromatin state were obtained from the University of California Santa Cruz using the *UcscTrack()* function from the *Gviz* R package. Chromatin states of GM12878, H1-hESC, and NHLF cell lines are displayed based on their relevance to blood monocytes, prenatal development, and lung health and function, respectively.

**Supplementary Figure 14.**
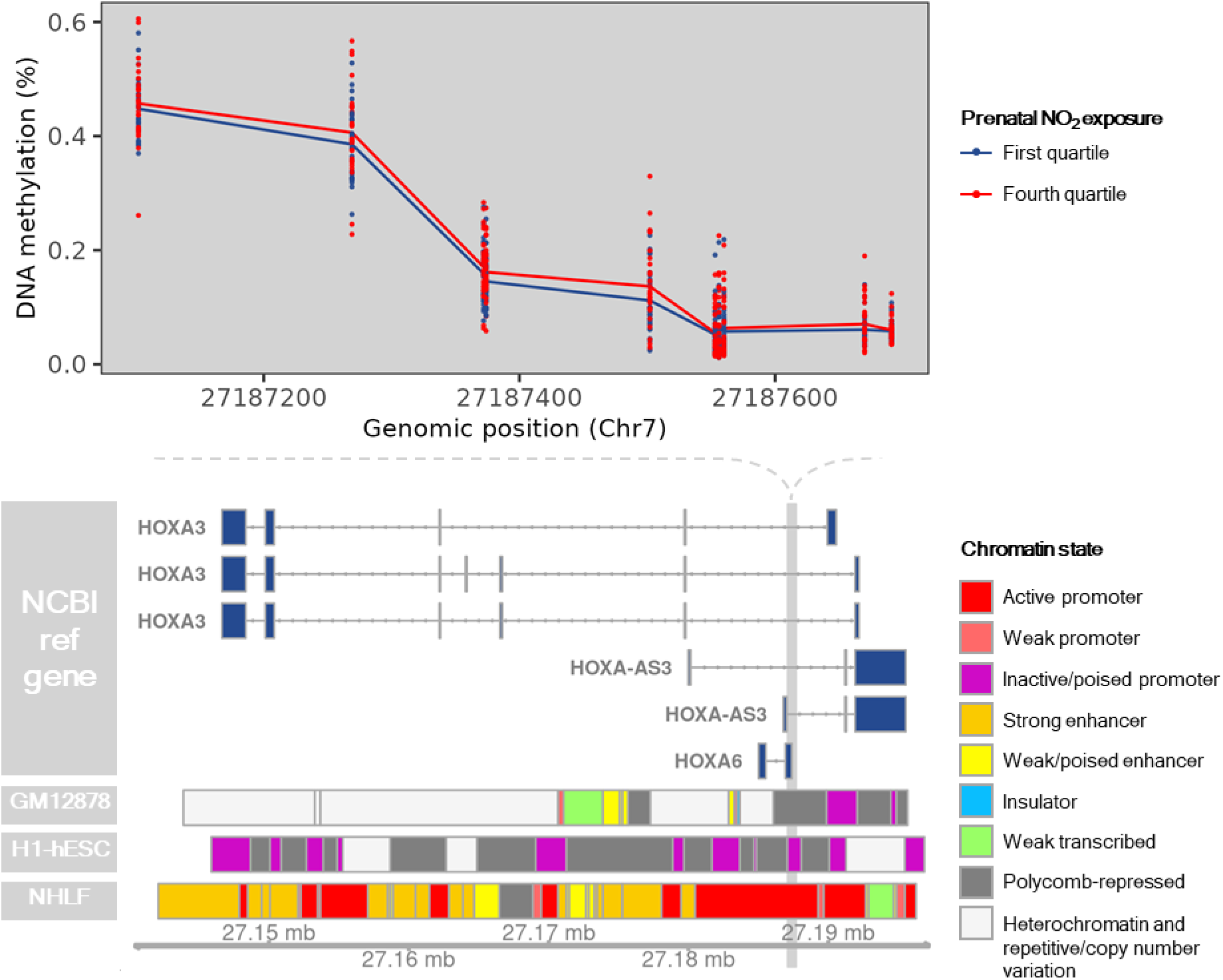
DNA methylation and chromatin state across the cord blood differentially methylated region (DMR) located on Chr7: 27187102-27187692. DNA methylation at CpGs contained within Chr7: 27187102-27187692 is shown for individuals with prenatal NO_2_ exposure < 7.3 ppb (first quartile; N=32; blue) or prenatal NO_2_ exposure > 16.3 ppb (fourth quartile; N=32; red) is displayed. Annotation of known genes and chromatin state were obtained from the University of California Santa Cruz using the *UcscTrack()* function from the *Gviz* R package. Chromatin states of GM12878, H1-hESC, and NHLF cell lines are displayed based on their relevance to blood monocytes, prenatal development, and lung health and function, respectively.

**Supplementary Figure 15.**
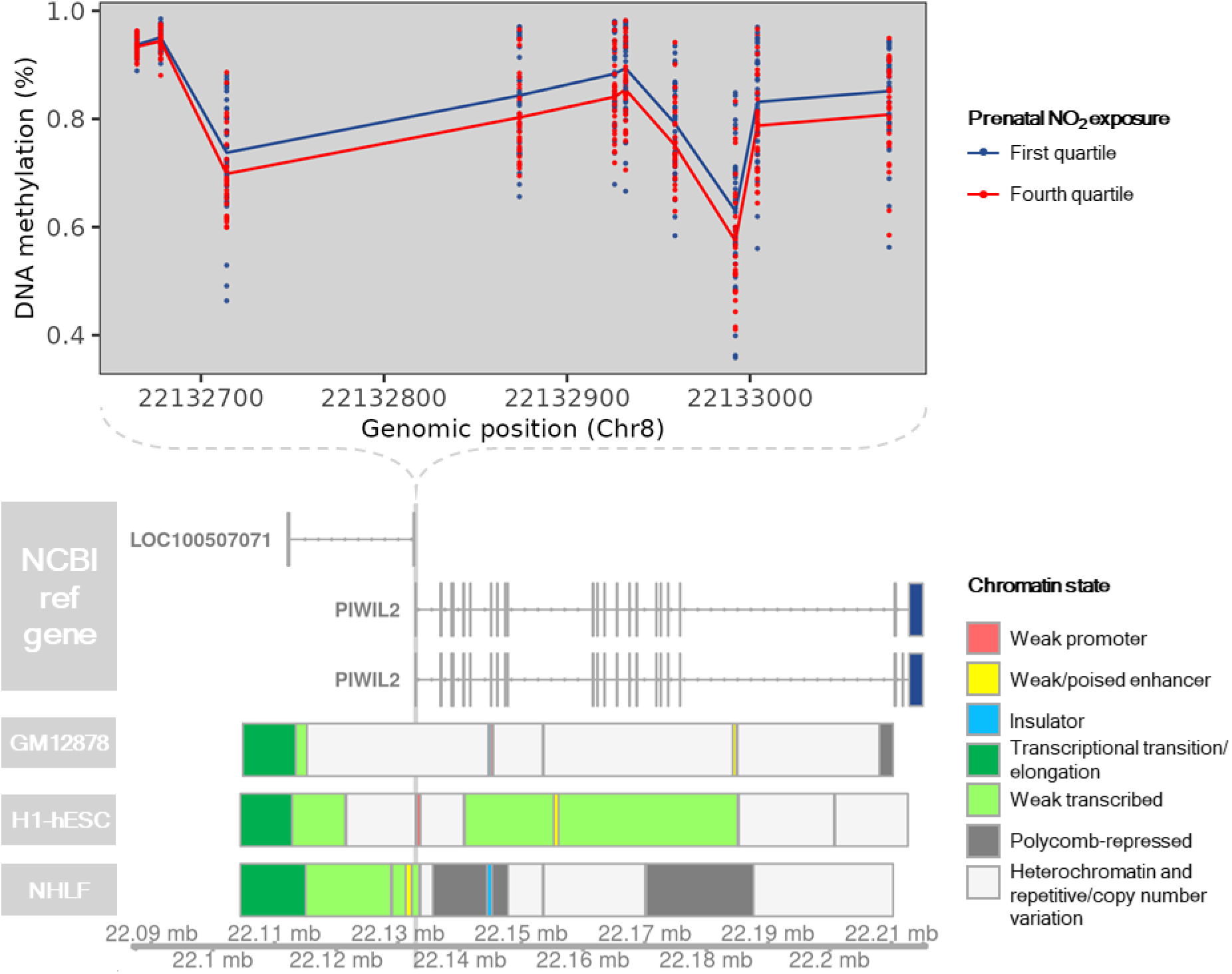
DNA methylation and chromatin state across the cord blood differentially methylated region (DMR) located on Chr8: 22132665-22133077. DNA methylation at CpGs contained within Chr8:22132665-22133077 is shown for individuals with prenatal NO_2_ exposure < 7.3 ppb (first quartile; N=32; blue) or prenatal NO_2_ exposure > 16.3 ppb (fourth quartile; N=32; red) is displayed. Annotation of known genes and chromatin state were obtained from the University of California Santa Cruz using the *UcscTrack()* function from the *Gviz* R package. Chromatin states of GM12878, H1-hESC, and NHLF cell lines are displayed based on their relevance to blood monocytes, prenatal development, and lung health and function, respectively.

**Supplementary Figure 16.**
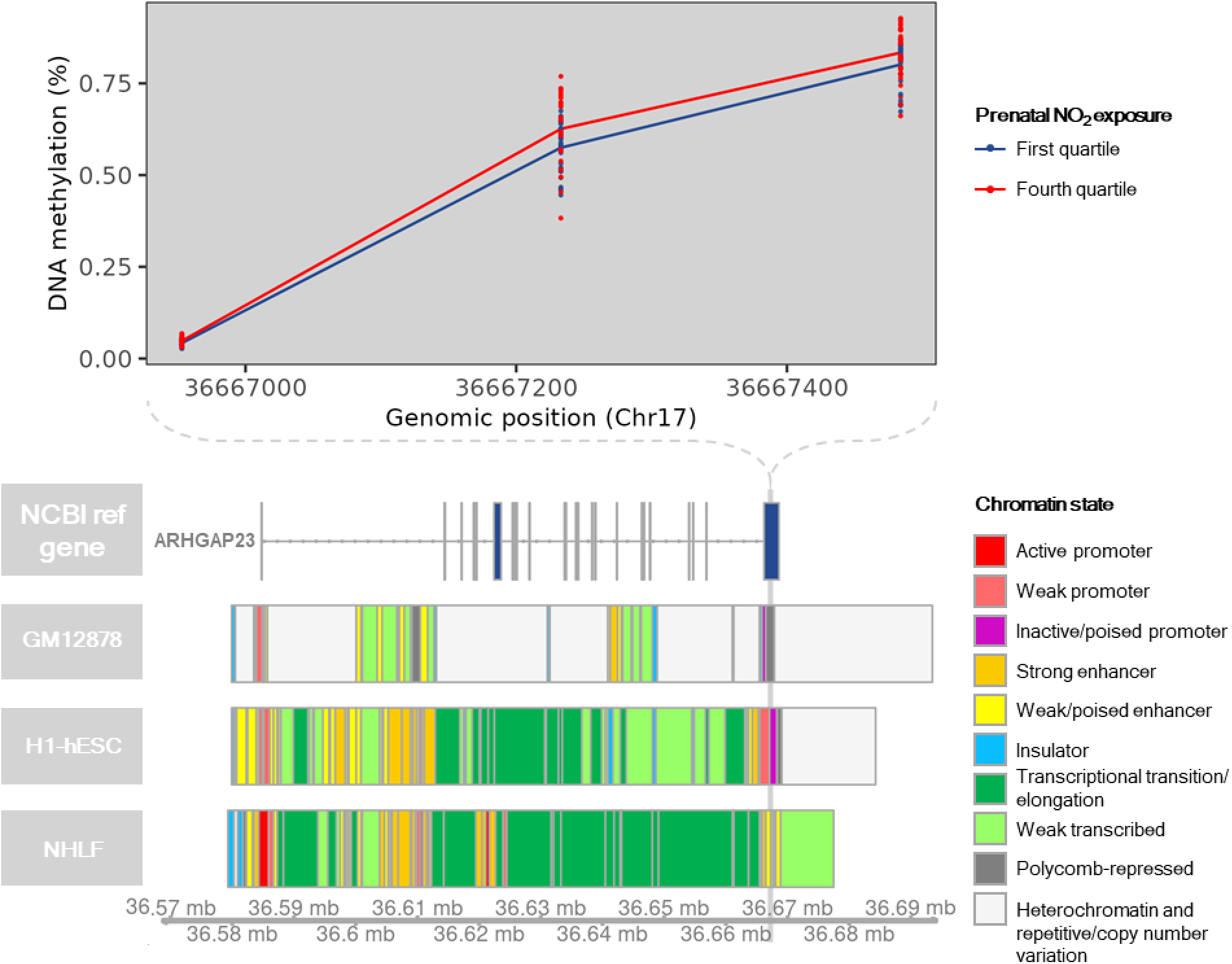
DNA methylation and chromatin state across the cord blood differentially methylated region (DMR) located on Chr17:36666953-36667485. DNA methylation at CpGs contained within Chr17:36666953-36667485 is shown for individuals with prenatal NO_2_ exposure < 7.3 ppb (first quartile; N=32; blue) or prenatal NO_2_ exposure > 16.3 ppb (fourth quartile; N=32; red) is displayed. Annotation of known genes and chromatin state were obtained from the University of California Santa Cruz using the *UcscTrack()* function from the *Gviz* R package. Chromatin states of GM12878, H1-hESC, and NHLF cell lines are displayed based on their relevance to blood monocytes, prenatal development, and lung health and function, respectively.

**Supplementary Figure 17.**
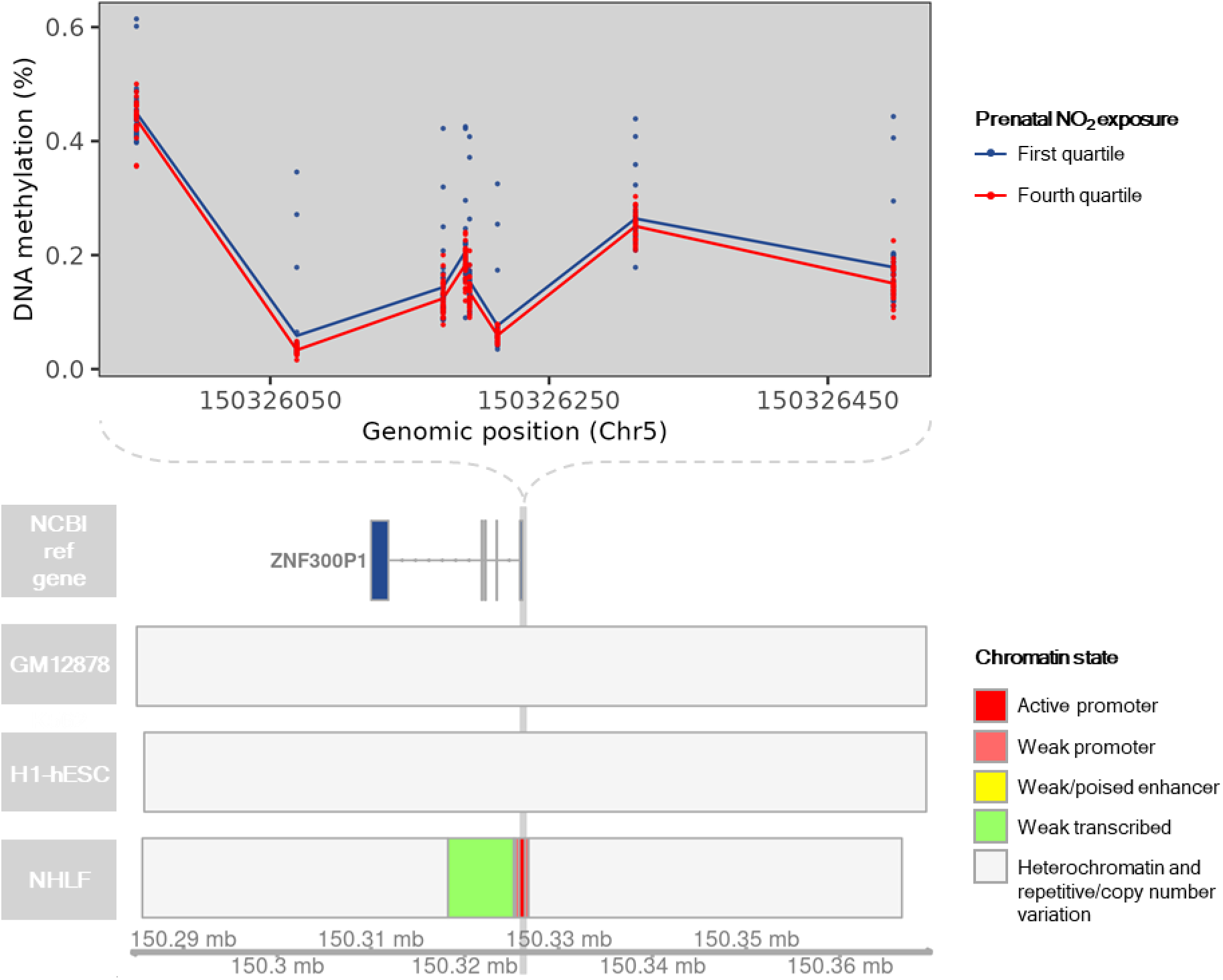
DNA methylation and chromatin state across the cord blood differentially methylated region (DMR) located on Chr5:150325954-150326498. DNA methylation at CpGs contained within Chr5:150325954-150326498 is shown for individuals with prenatal NO_2_ exposure < 7.3 ppb (first quartile; N=32; blue) or prenatal NO_2_ exposure > 16.3 ppb (fourth quartile; N=32; red) is displayed. Annotation of known genes and chromatin state were obtained from the University of California Santa Cruz using the *UcscTrack()* function from the *Gviz* R package. Chromatin states of GM12878, H1-hESC, and NHLF cell lines are displayed based on their relevance to blood monocytes, prenatal development, and lung health and function, respectively.

**Supplementary Figure 18.**
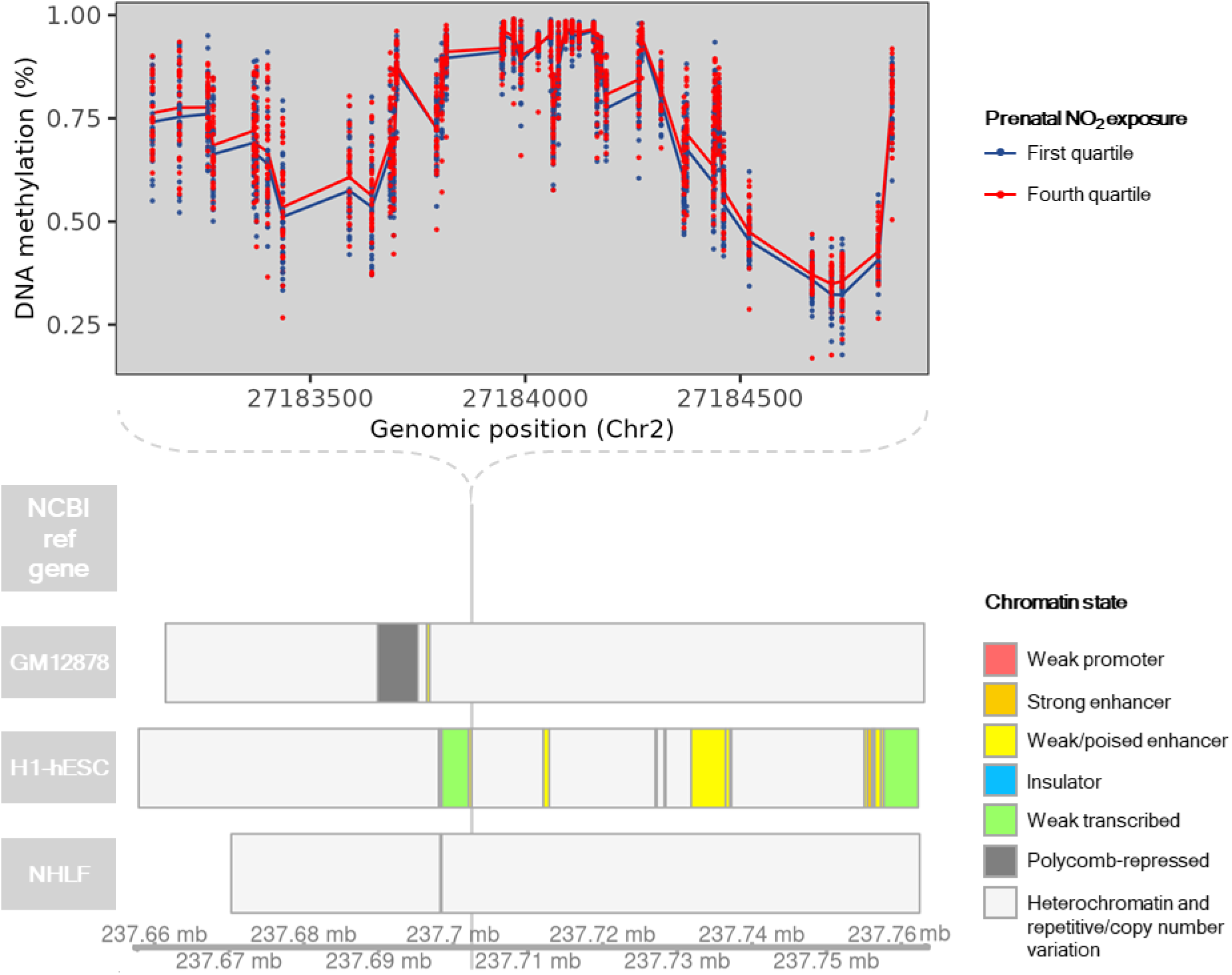
DNA methylation and chromatin state across the cord blood differentially methylated region (DMR) located on Chr2:237702390-237702581. DNA methylation at CpGs contained within Chr2:237702390-237702581 is shown for individuals with prenatal NO_2_ exposure < 7.3 ppb (first quartile; N=32; blue) or prenatal NO_2_ exposure > 16.3 ppb (fourth quartile; N=32; red) is displayed. Annotation of known genes and chromatin state were obtained from the University of California Santa Cruz using the *UcscTrack()* function from the *Gviz* R package. Chromatin states of GM12878, H1-hESC, and NHLF cell lines are displayed based on their relevance to blood monocytes, prenatal development, and lung health and function, respectively.

**Supplementary Figure 19.**
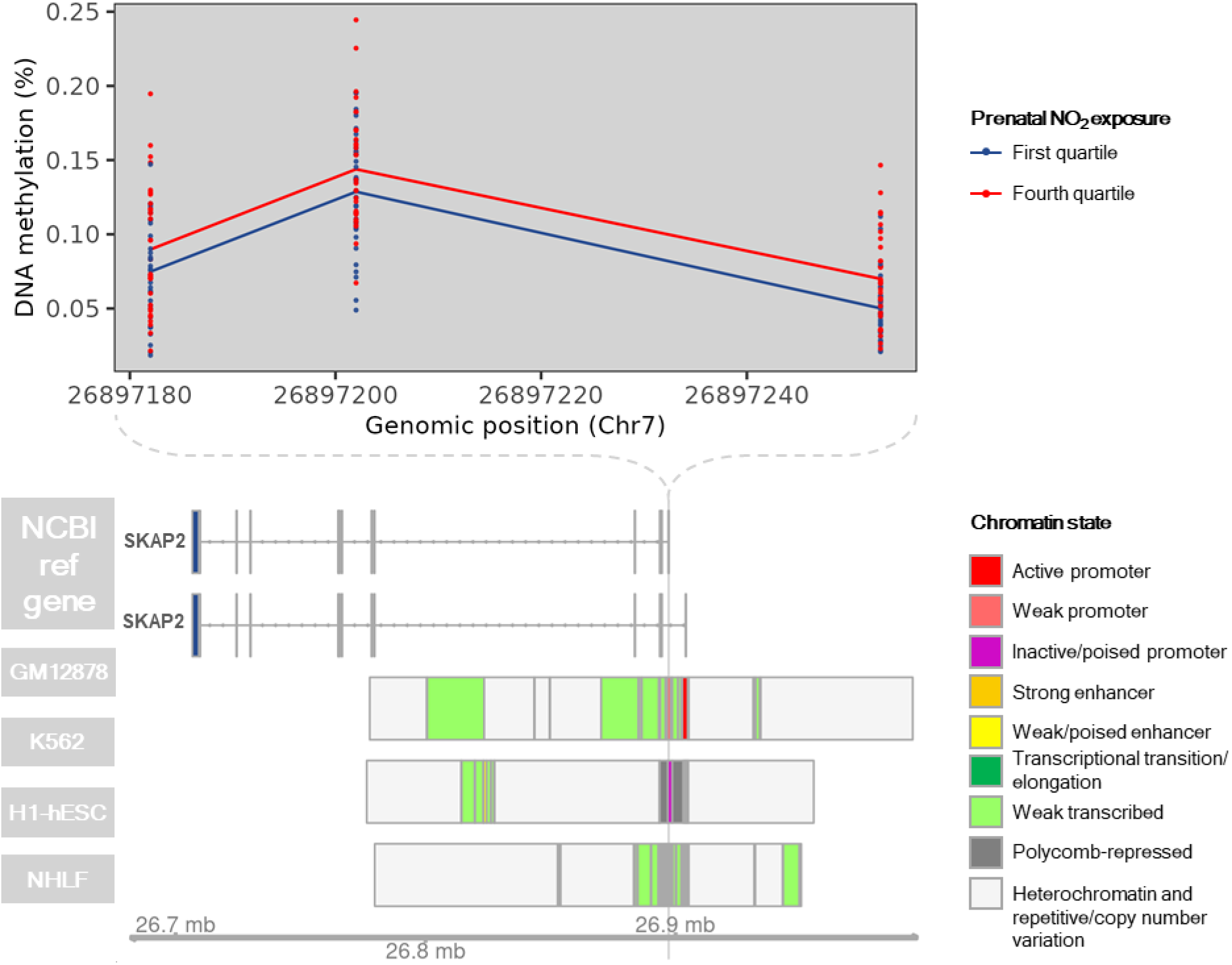
DNA methylation and chromatin state across the cord blood differentially methylated region (DMR) located on Chr7:26897182-26897254. DNA methylation at CpGs contained within Chr7:26897182-26897254 is shown for individuals with prenatal NO_2_ exposure < 7.3 ppb (first quartile; N=32; blue) or prenatal NO_2_ exposure > 16.3 ppb (fourth quartile; N=32; red) is displayed. Annotation of known genes and chromatin state were obtained from the University of California Santa Cruz using the *UcscTrack()* function from the *Gviz* R package. Chromatin states of GM12878, H1-hESC, and NHLF cell lines are displayed based on their relevance to blood monocytes, prenatal development, and lung health and function, respectively.

**Supplementary Figure 20.**
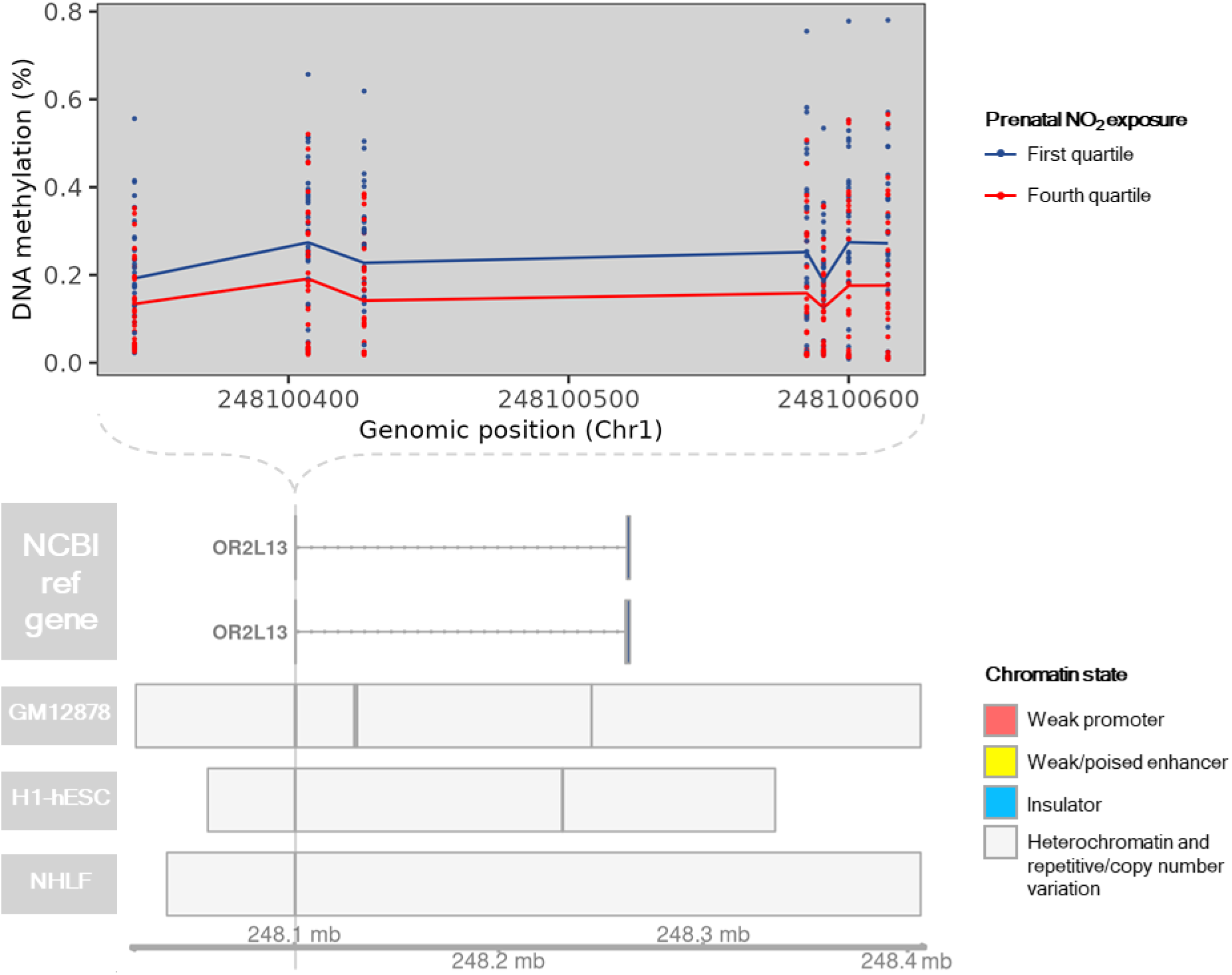
DNA methylation and chromatin state across the cord blood differentially methylated region (DMR) located on Chr1:248100345-248100615. DNA methylation at CpGs contained within Chr1:248100345-248100615 is shown for individuals with prenatal NO_2_ exposure < 7.3 ppb (first quartile; N=32; blue) or prenatal NO_2_ exposure > 16.3 ppb (fourth quartile; N=32; red) is displayed. Annotation of known genes and chromatin state were obtained from the University of California Santa Cruz using the *UcscTrack()* function from the *Gviz* R package. Chromatin states of GM12878, H1-hESC, and NHLF cell lines are displayed based on their relevance to blood monocytes, prenatal development, and lung health and function, respectively.

**Supplementary Figure 21.**
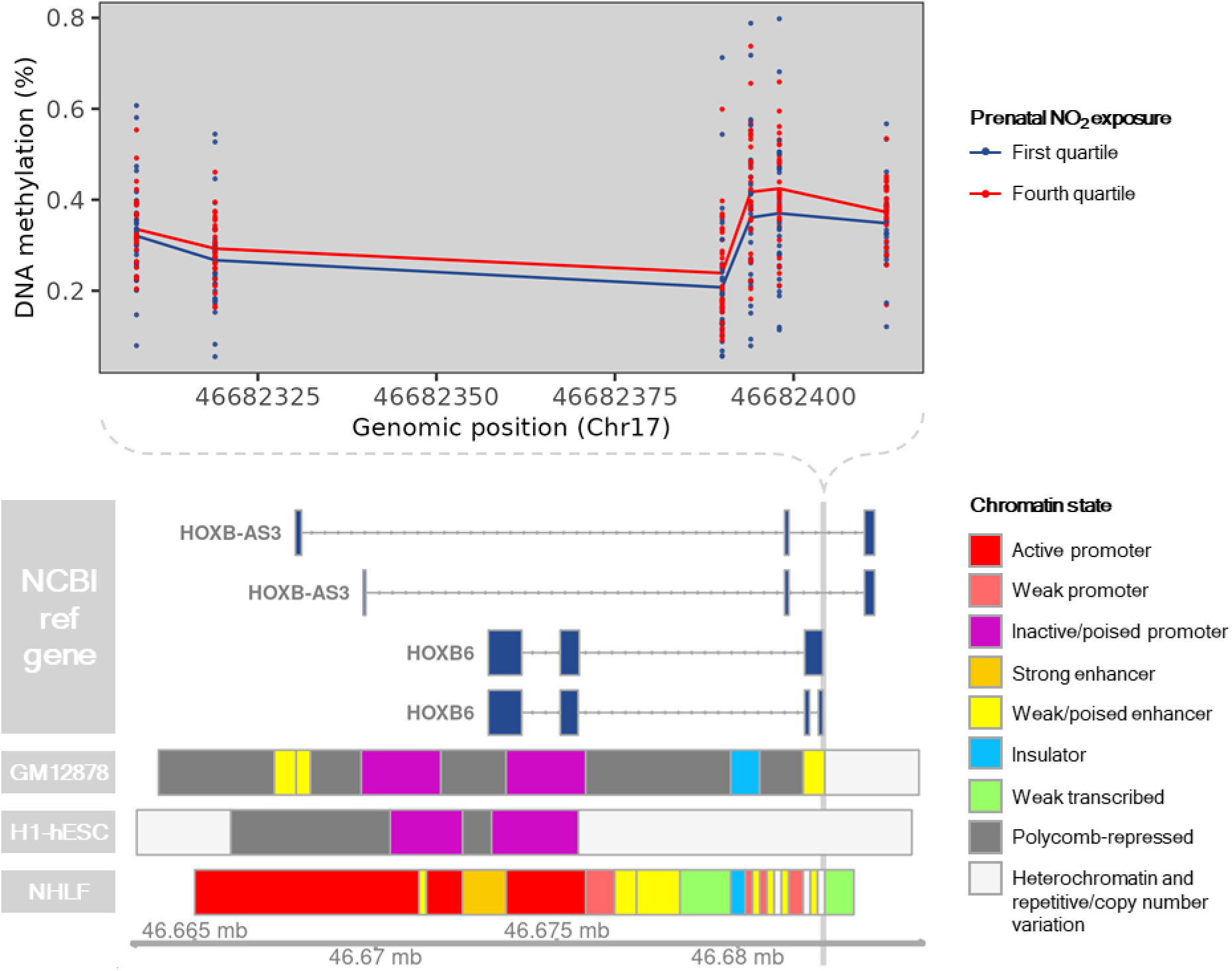
DNA methylation and chromatin state across the cord blood differentially methylated region (DMR) located on Chr17:46682308-46682414. DNA methylation at CpGs contained within Chr17:46682308-46682414 is shown for individuals with prenatal NO_2_ exposure < 7.3 ppb (first quartile; N=32; blue) or prenatal NO_2_ exposure > 16.3 ppb (fourth quartile; N=32; red) is displayed. Annotation of known genes and chromatin state were obtained from the University of California Santa Cruz using the *UcscTrack()* function from the *Gviz* R package. Chromatin states of GM12878, H1-hESC, and NHLF cell lines are displayed based on their relevance to blood monocytes, prenatal development, and lung health and function, respectively.

**Supplementary Figure 22.**
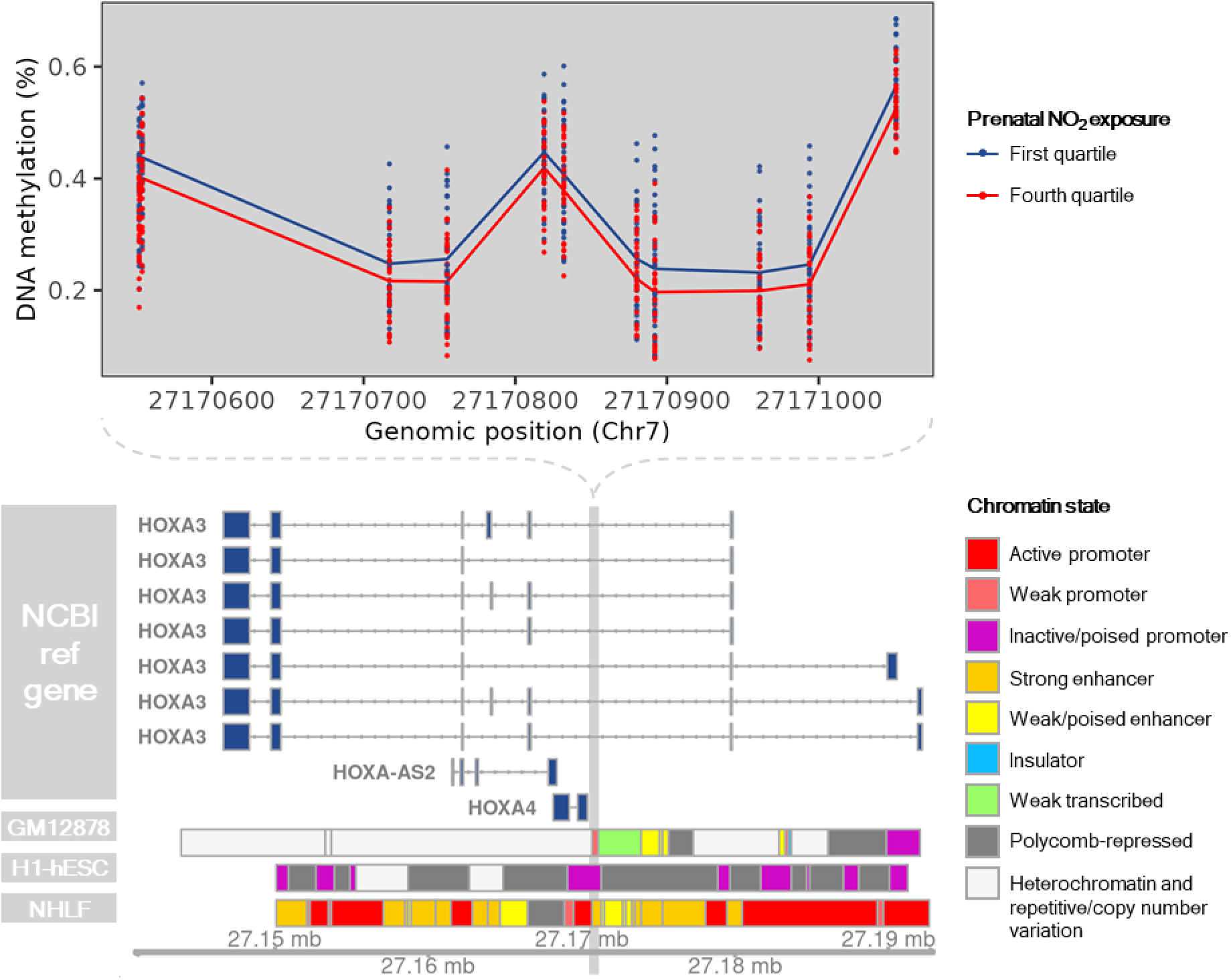
DNA methylation and chromatin state across the cord blood differentially methylated region (DMR) located on Chr7:27170552-27171052. DNA methylation at CpGs contained within Chr7:27170552-27171052 is shown for individuals with prenatal NO_2_ exposure < 7.3 ppb (first quartile; N=32; blue) or prenatal NO_2_ exposure > 16.3 ppb (fourth quartile; N=32; red) is displayed. Annotation of known genes and chromatin state were obtained from the University of California Santa Cruz using the *UcscTrack()* function from the *Gviz* R package. Chromatin states of GM12878, H1-hESC, and NHLF cell lines are displayed based on their relevance to blood monocytes, prenatal development, and lung health and function, respectively.

**Supplementary Figure 23.**
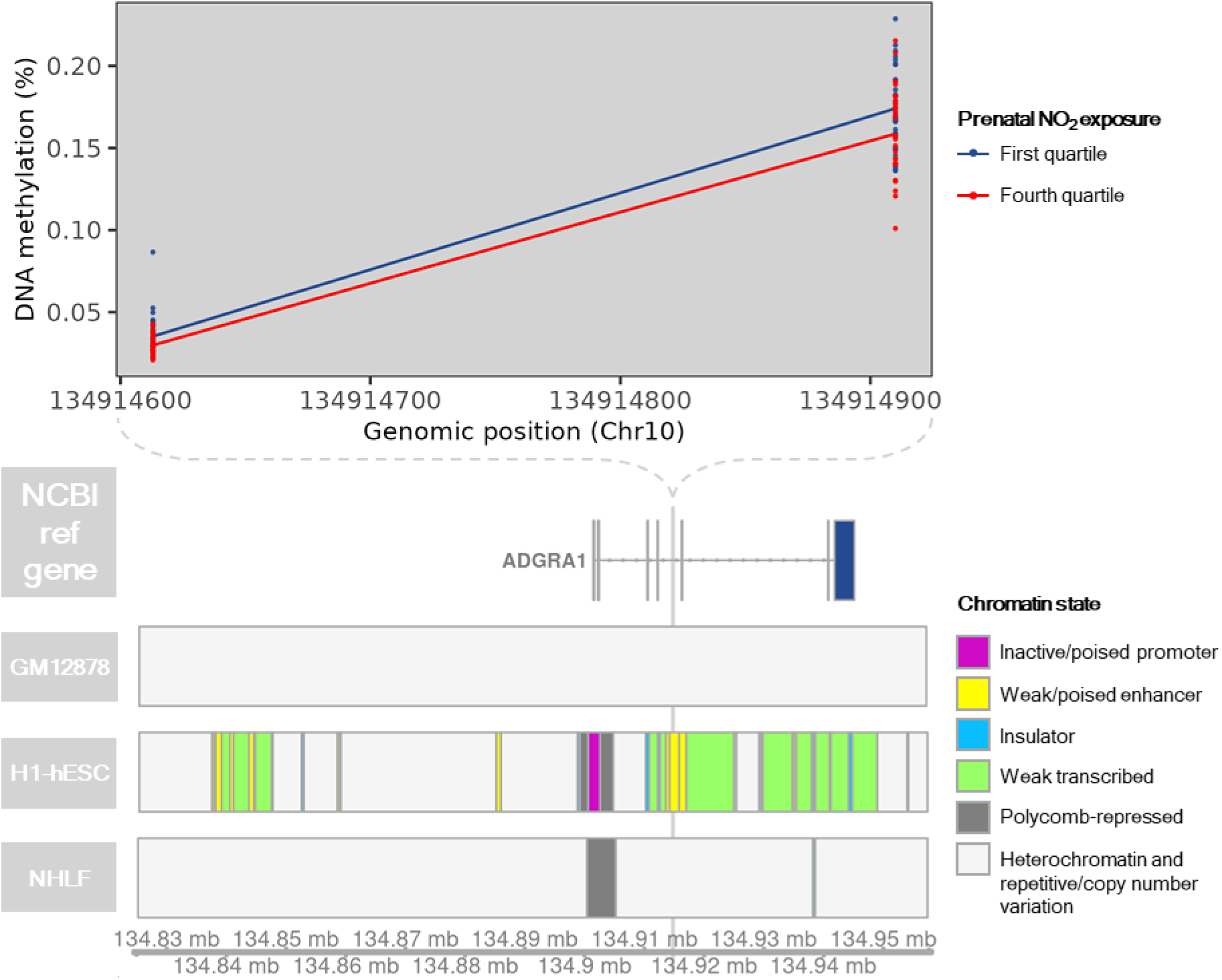
DNA methylation and chromatin state across the cord blood differentially methylated region (DMR) located on Chr10:134914613-134914911. DNA methylation at CpGs contained within Chr10:134914613-134914911 is shown for individuals with prenatal NO_2_ exposure < 7.3 ppb (first quartile; N=32; blue) or prenatal NO_2_ exposure > 16.3 ppb (fourth quartile; N=32; red) is displayed. Annotation of known genes and chromatin state were obtained from the University of California Santa Cruz using the *UcscTrack()* function from the *Gviz* R package. Chromatin states of GM12878, H1-hESC, and NHLF cell lines are displayed based on their relevance to blood monocytes, prenatal development, and lung health and function, respectively. *ADGRA1* is also known as *GPR123*.

**Supplementary Figure 24.**
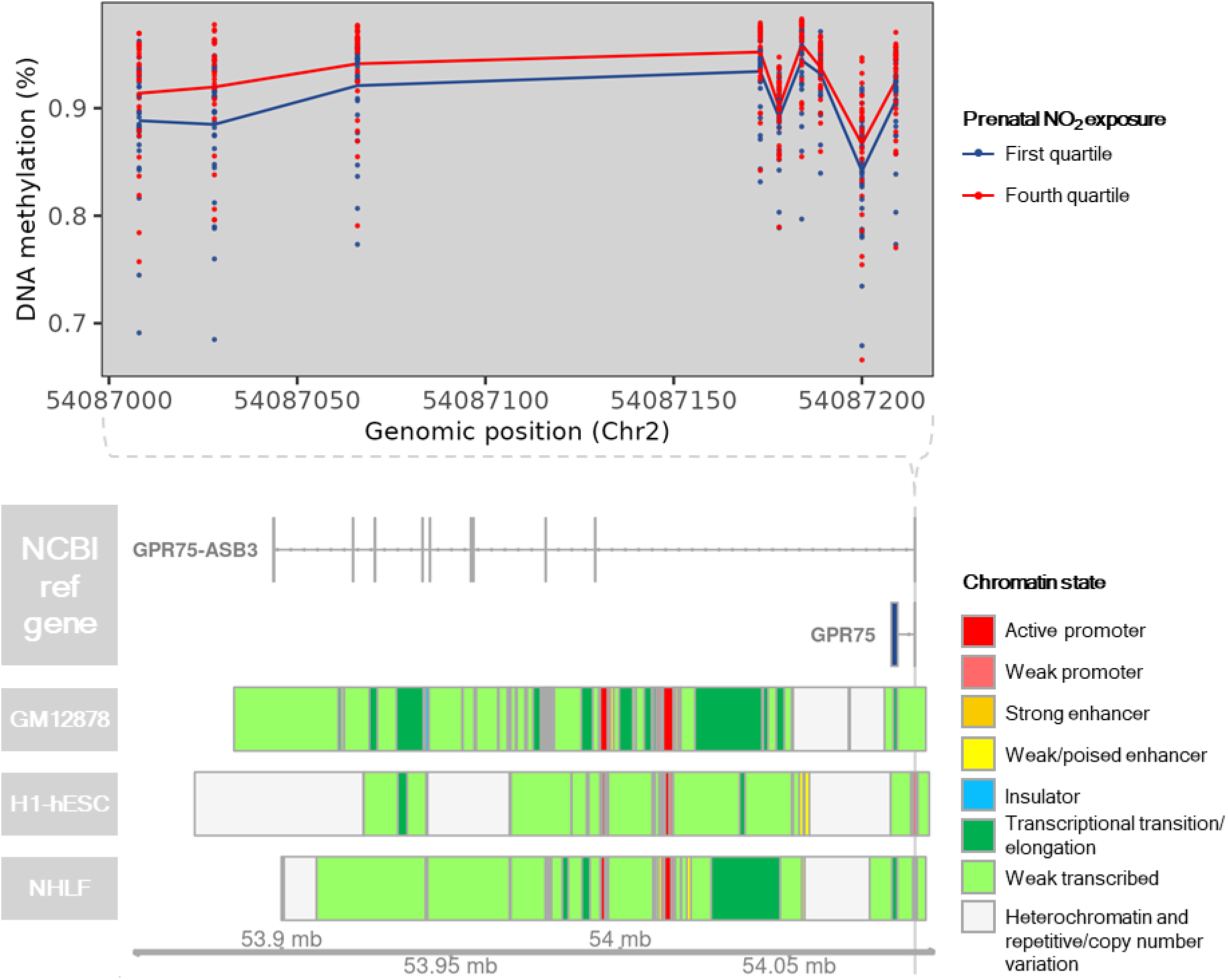
DNA methylation and chromatin state across the cord blood differentially methylated region (DMR) located on Chr2:54087008-54087210. DNA methylation at CpGs contained within Chr2:54087008-54087210 is shown for individuals with prenatal NO_2_ exposure < 7.3 ppb (first quartile; N=32; blue) or prenatal NO_2_ exposure > 16.3 ppb (fourth quartile; N=32; red) is displayed. Annotation of known genes and chromatin state were obtained from the University of California Santa Cruz using the *UcscTrack()* function from the *Gviz* R package. Chromatin states of GM12878, H1-hESC, and NHLF cell lines are displayed based on their relevance to blood monocytes, prenatal development, and lung health and function, respectively.

**Supplementary Figure 25.**
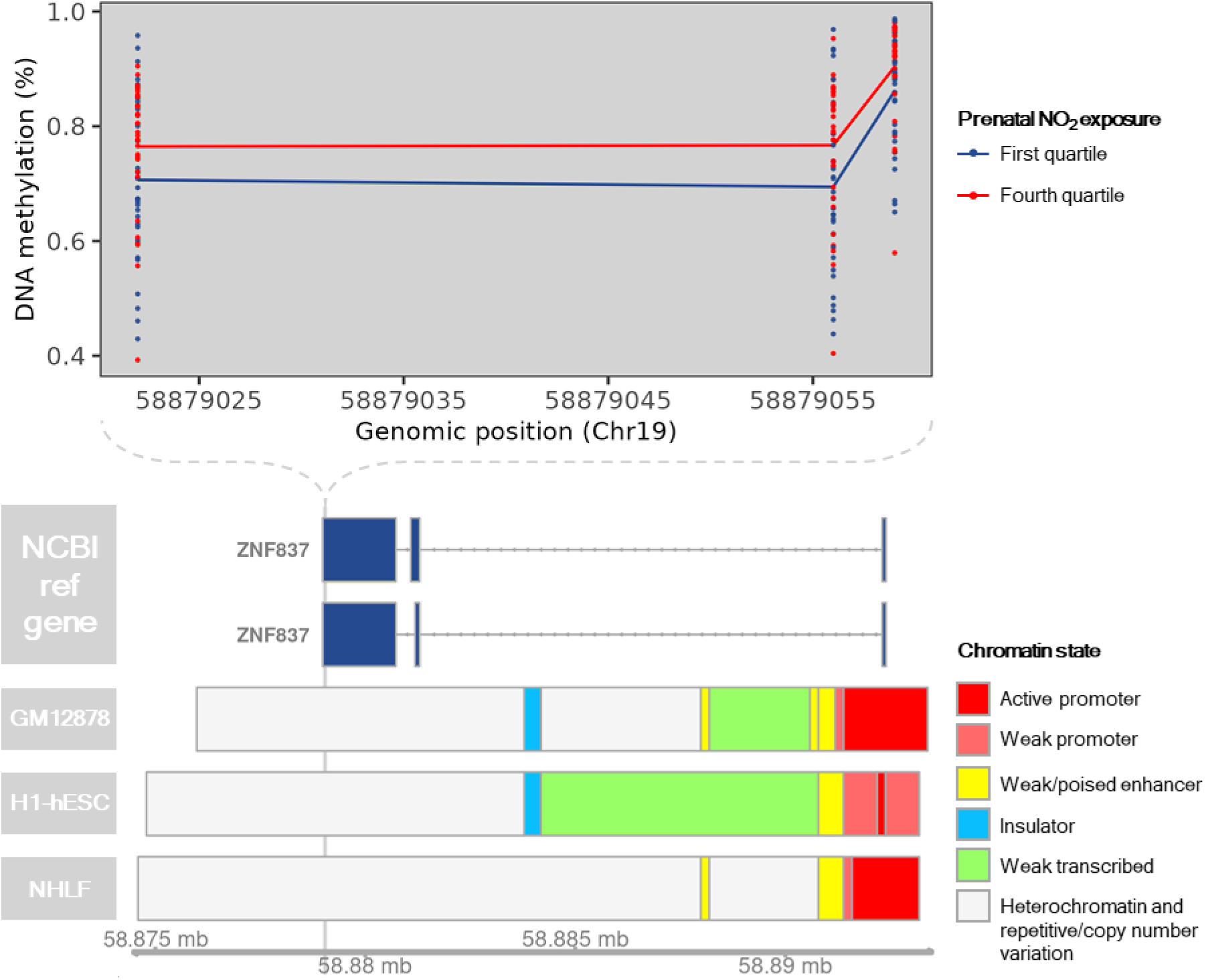
DNA methylation and chromatin state across the cord blood differentially methylated region (DMR) located on Chr19:58879022-58879060. DNA methylation at CpGs contained within Chr19:58879022-58879060 is shown for individuals with prenatal NO_2_ exposure < 7.3 ppb (first quartile; N=32; blue) or prenatal NO_2_ exposure > 16.3 ppb (fourth quartile; N=32; red) is displayed. Annotation of known genes and chromatin state were obtained from the University of California Santa Cruz using the *UcscTrack()* function from the *Gviz* R package. Chromatin states of GM12878, H1-hESC, and NHLF cell lines are displayed based on their relevance to blood monocytes, prenatal development, and lung health and function, respectively.

**Supplementary Figure 26.**
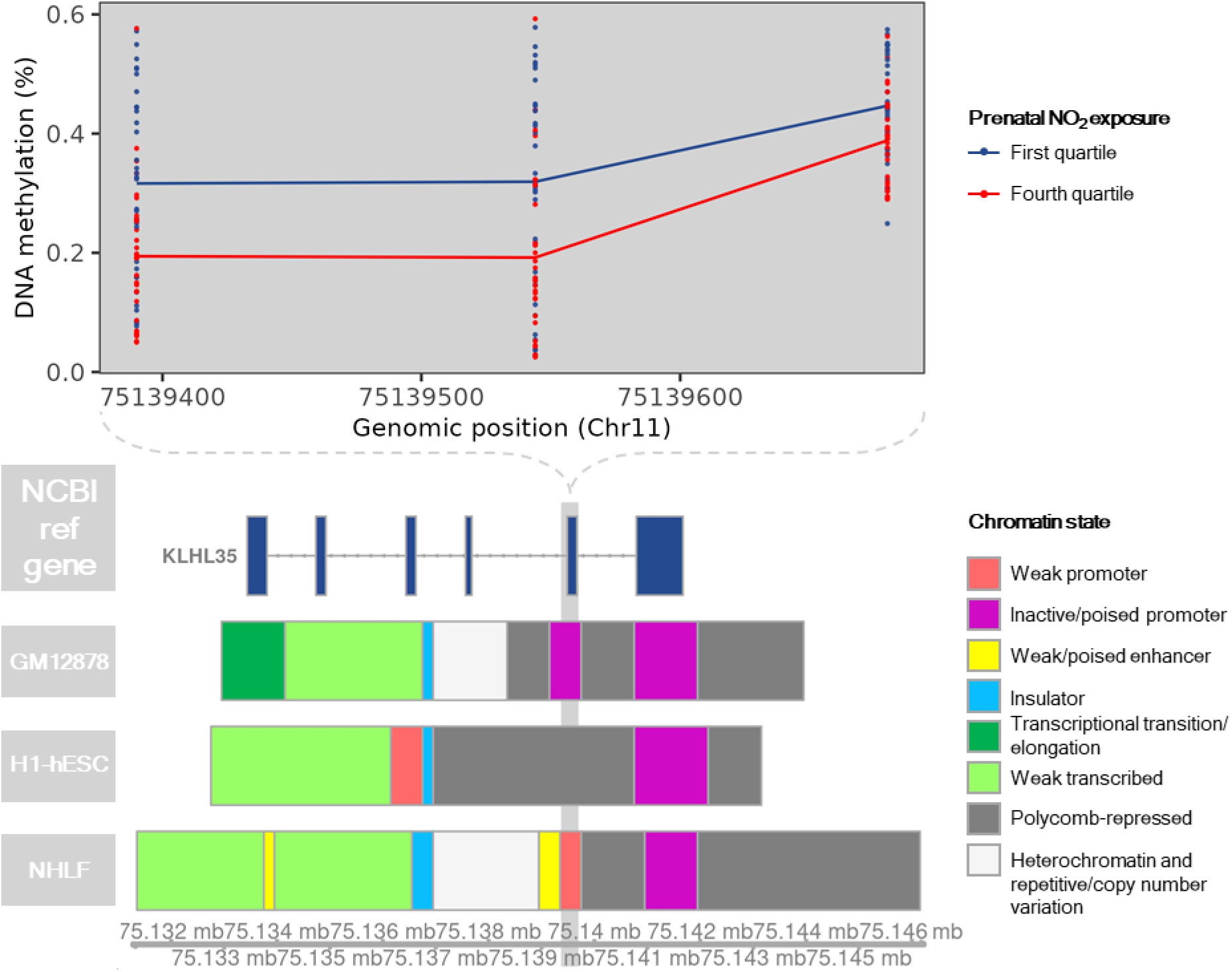
DNA methylation and chromatin state across the cord blood differentially methylated region (DMR) located on Chr11:75139390-75139681. DNA methylation at CpGs contained within Chr11:75139390-75139681 is shown for individuals with prenatal NO_2_ exposure < 7.3 ppb (first quartile; N=32; blue) or prenatal NO_2_ exposure > 16.3 ppb (fourth quartile; N=32; red) is displayed. Annotation of known genes and chromatin state were obtained from the University of California Santa Cruz using the *UcscTrack()* function from the *Gviz* R package. Chromatin states of GM12878, H1-hESC, and NHLF cell lines are displayed based on their relevance to blood monocytes, prenatal development, and lung health and function, respectively.

**Supplementary Figure 27.**
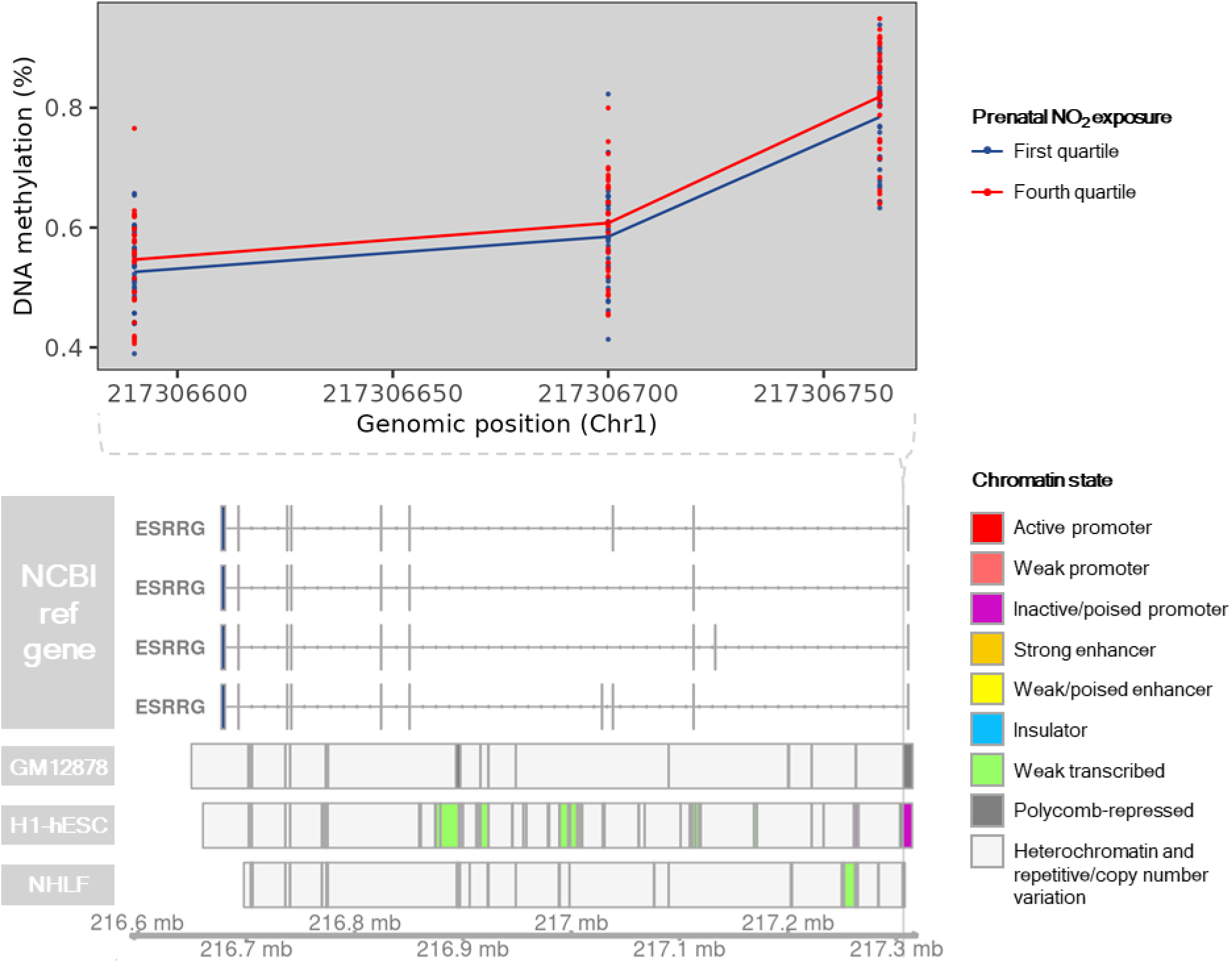
DNA methylation and chromatin state across the cord blood differentially methylated region (DMR) located on Chr1:217306590-217306764. DNA methylation at CpGs contained within Chr1:217306590-217306764 is shown for individuals with prenatal NO_2_ exposure < 7.3 ppb (first quartile; N=32; blue) or prenatal NO_2_ exposure > 16.3 ppb (fourth quartile; N=32; red) is displayed. Annotation of known genes and chromatin state were obtained from the University of California Santa Cruz using the *UcscTrack()* function from the *Gviz* R package. Chromatin states of GM12878, H1-hESC, and NHLF cell lines are displayed based on their relevance to blood monocytes, prenatal development, and lung health and function, respectively.

**Supplementary Figure 28.**
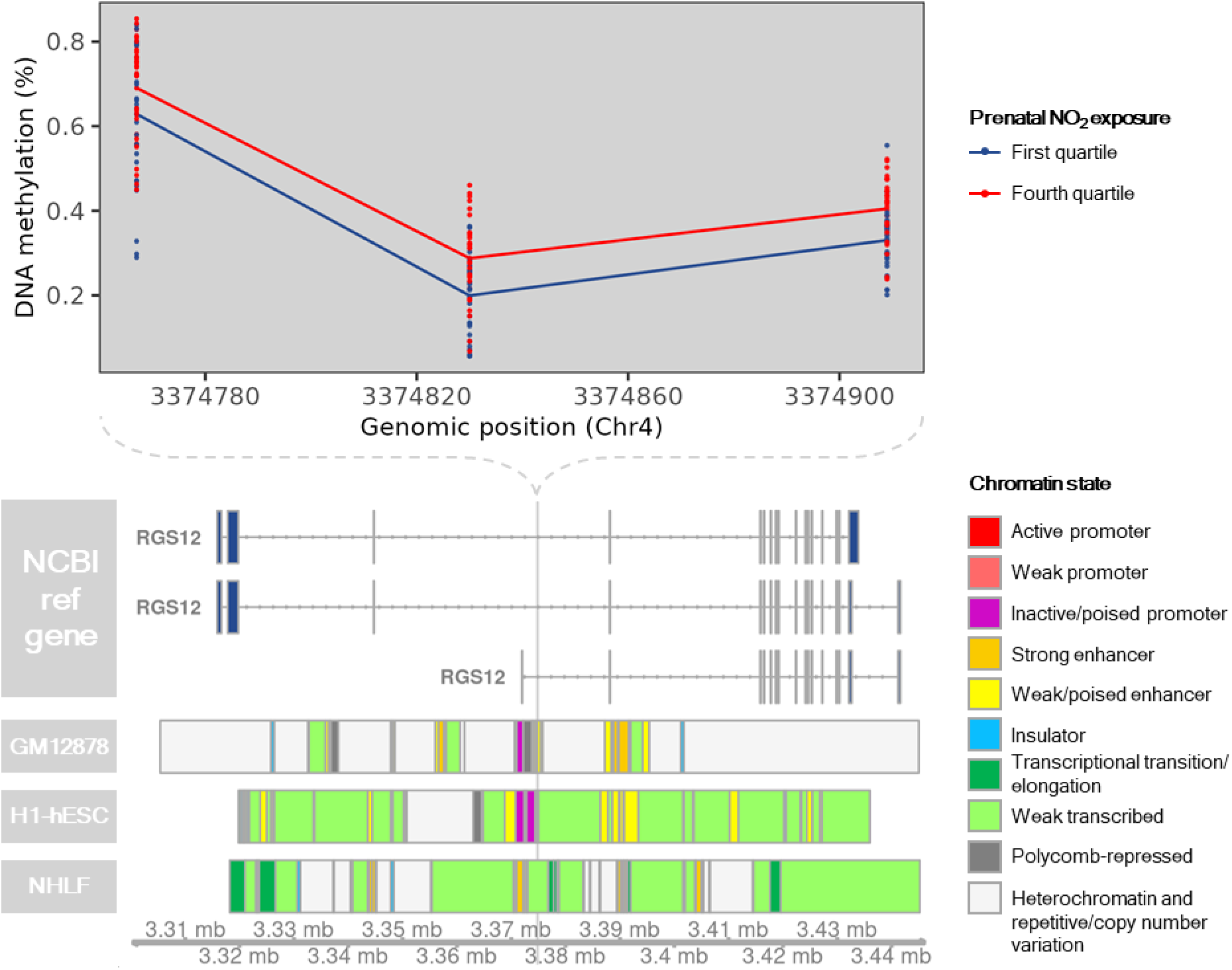
DNA methylation and chromatin state across the cord blood differentially methylated region (DMR) located on Chr4:3374767-3374910. DNA methylation at CpGs contained within Chr4:3374767-3374910 is shown for individuals with prenatal NO_2_ exposure < 7.3 ppb (first quartile; N=32; blue) or prenatal NO_2_ exposure > 16.3 ppb (fourth quartile; N=32; red) is displayed. Annotation of known genes and chromatin state were obtained from the University of California Santa Cruz using the *UcscTrack()* function from the *Gviz* R package. Chromatin states of GM12878, H1-hESC, and NHLF cell lines are displayed based on their relevance to blood monocytes, prenatal development, and lung health and function, respectively.

**Supplementary Figure 29.**
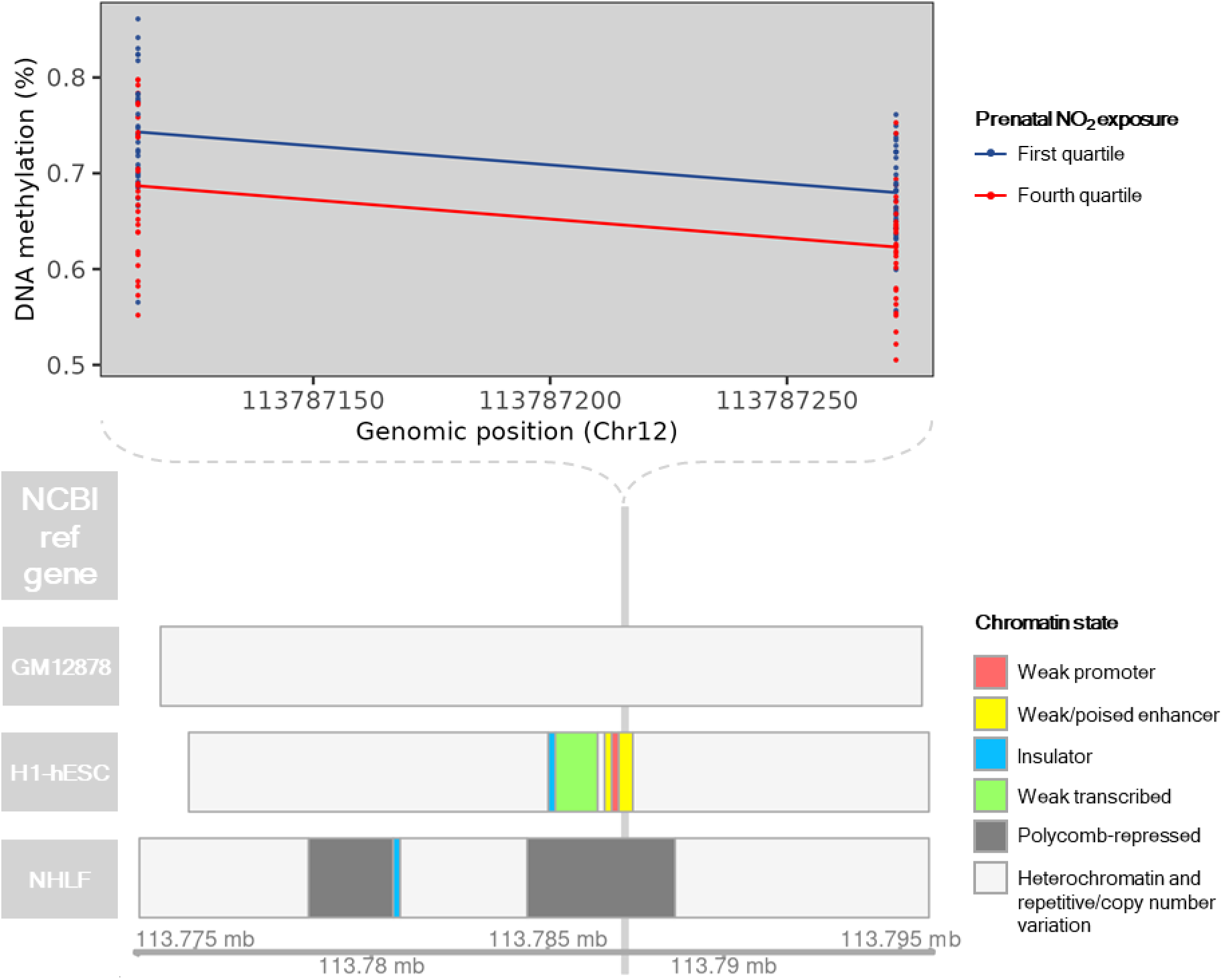
DNA methylation and chromatin state across the cord blood differentially methylated region (DMR) located on Chr12:113787113-113787274. DNA methylation at CpGs contained within Chr12:113787113-113787274 is shown for individuals with prenatal NO_2_ exposure < 7.3 ppb (first quartile; N=32; blue) or prenatal NO_2_ exposure > 16.3 ppb (fourth quartile; N=32; red) is displayed. Annotation of known genes and chromatin state were obtained from the University of California Santa Cruz using the *UcscTrack()* function from the *Gviz* R package. Chromatin states of GM12878, H1-hESC, and NHLF cell lines are displayed based on their relevance to blood monocytes, prenatal development, and lung health and function, respectively.

**Supplementary Figure 30.**
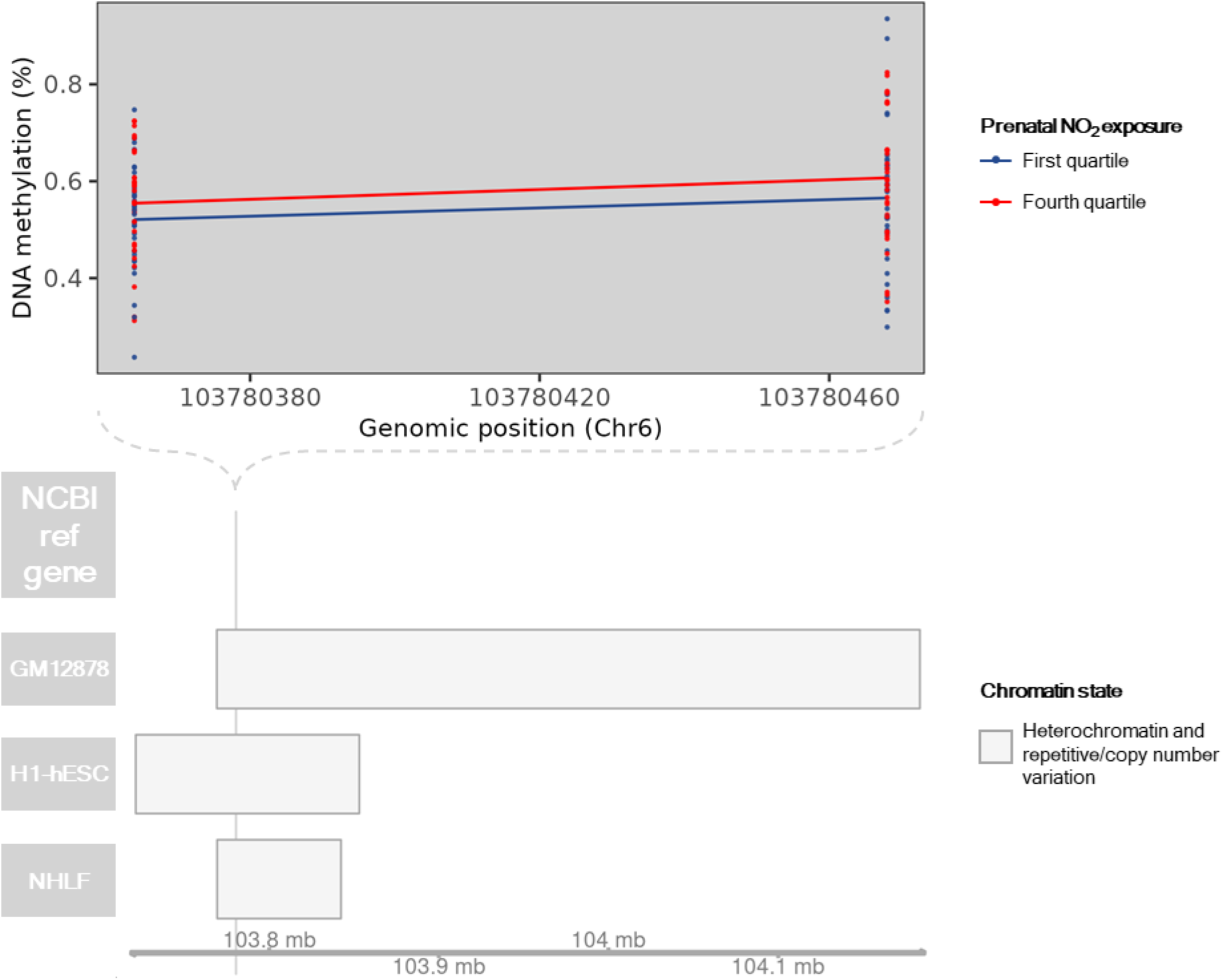
DNA methylation and chromatin state across the cord blood differentially methylated region (DMR) located on Chr6:103780364-103780469. DNA methylation at CpGs contained within Chr6:103780364-103780469 is shown for individuals with prenatal NO_2_ exposure < 7.3 ppb (first quartile; N=32; blue) or prenatal NO_2_ exposure > 16.3 ppb (fourth quartile; N=32; red) is displayed. Annotation of known genes and chromatin state were obtained from the University of California Santa Cruz using the *UcscTrack()* function from the *Gviz* R package. Chromatin states of GM12878, H1-hESC, and NHLF cell lines are displayed based on their relevance to blood monocytes, prenatal development, and lung health and function, respectively.

**Supplementary Figure 31.**
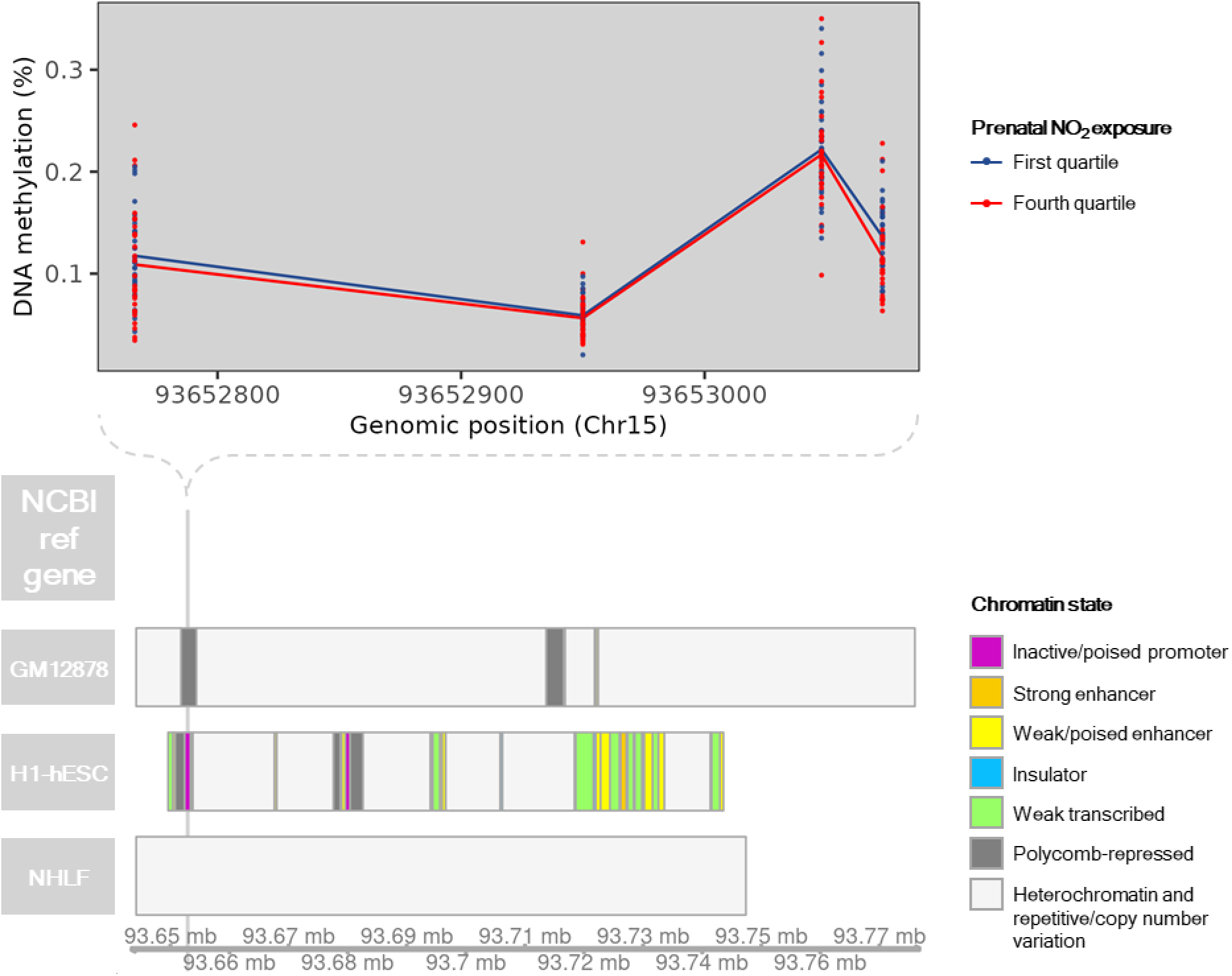
DNA methylation and chromatin state across the cord blood differentially methylated region (DMR) located on Chr15:93652766-93653074. DNA methylation at CpGs contained within Chr15:93652766-93653074 is shown for individuals with prenatal NO_2_ exposure < 7.3 ppb (first quartile; N=32; blue) or prenatal NO_2_ exposure > 16.3 ppb (fourth quartile; N=32; red) is displayed. Annotation of known genes and chromatin state were obtained from the University of California Santa Cruz using the *UcscTrack()* function from the *Gviz* R package. Chromatin states of GM12878, H1-hESC, and NHLF cell lines are displayed based on their relevance to blood monocytes, prenatal development, and lung health and function, respectively.

**Supplementary Figure 32.**
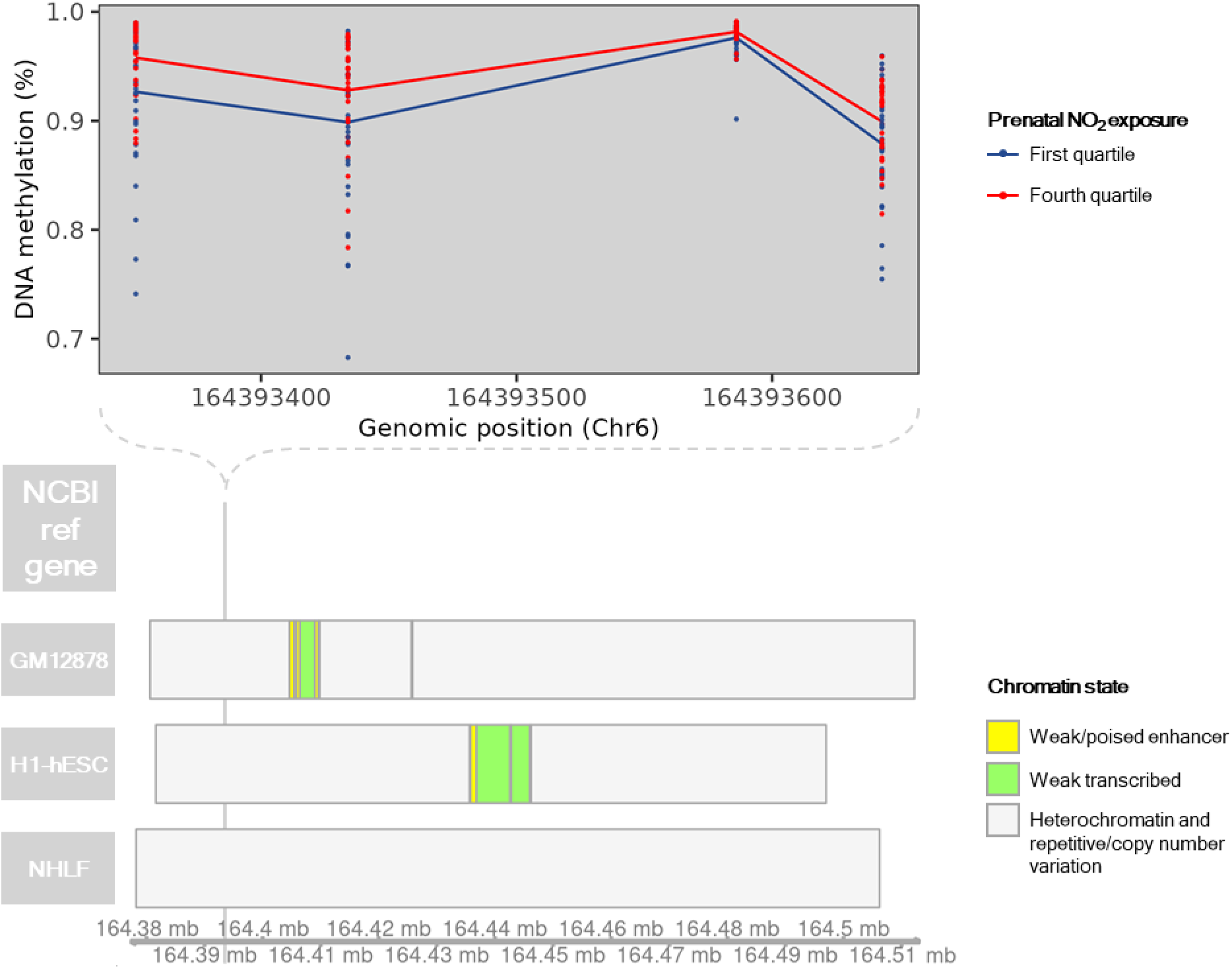
DNA methylation and chromatin state across the cord blood differentially methylated region (DMR) located on Chr6:164393351-164393644. DNA methylation at CpGs contained within Chr6:164393351-164393644 is shown for individuals with prenatal NO_2_ exposure < 7.3 ppb (first quartile; N=32; blue) or prenatal NO_2_ exposure > 16.3 ppb (fourth quartile; N=32; red) is displayed. Annotation of known genes and chromatin state were obtained from the University of California Santa Cruz using the *UcscTrack()* function from the *Gviz* R package. Chromatin states of GM12878, H1-hESC, and NHLF cell lines are displayed based on their relevance to blood monocytes, prenatal development, and lung health and function, respectively.

**Supplementary Figure 33.**
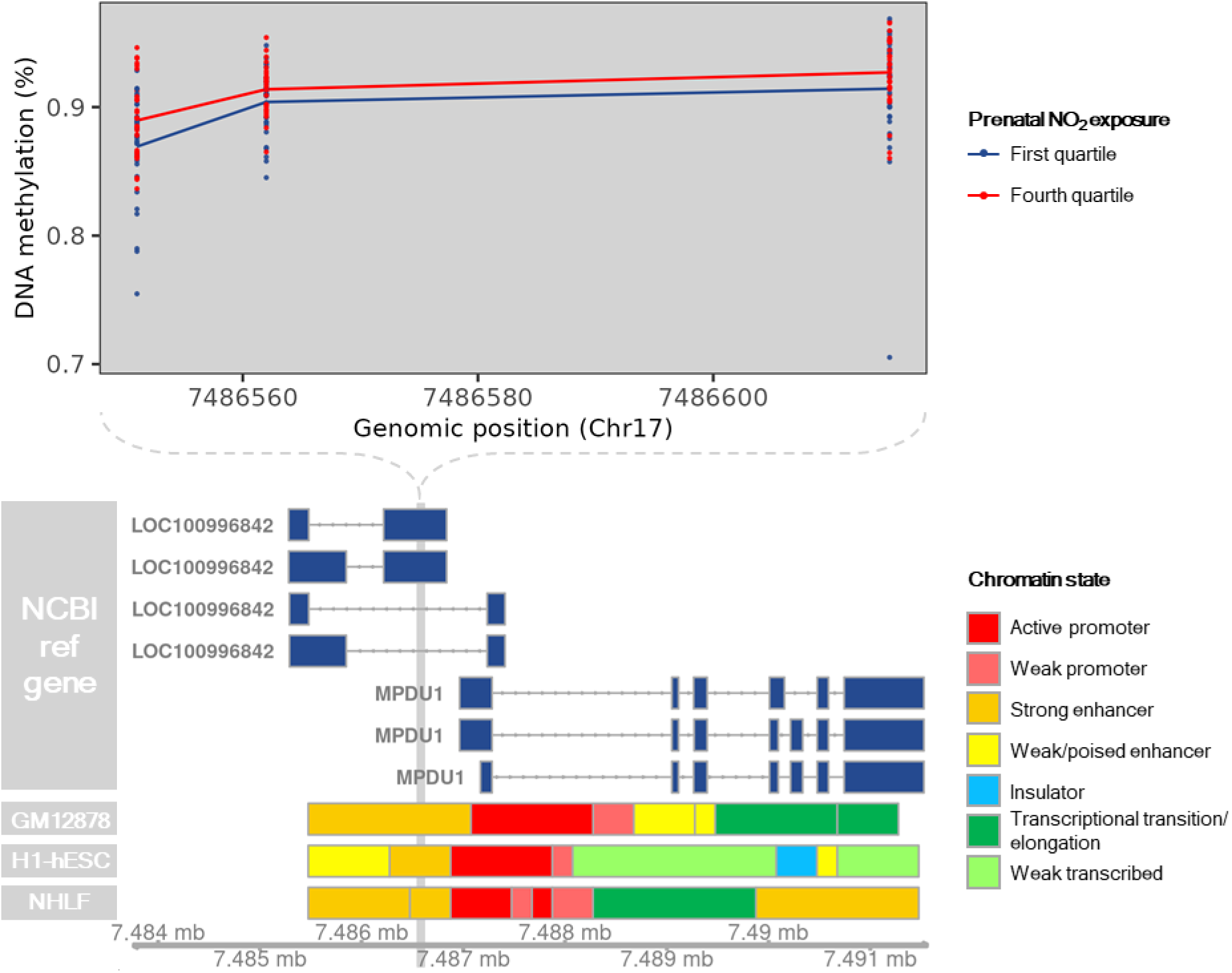
DNA methylation and chromatin state across the cord blood differentially methylated region (DMR) located on Chr17:7486551-7486616. DNA methylation at CpGs contained within Chr17:7486551-7486616 is shown for individuals with prenatal NO_2_ exposure < 7.3 ppb (first quartile; N=32; blue) or prenatal NO_2_ exposure > 16.3 ppb (fourth quartile; N=32; red) is displayed. Annotation of known genes and chromatin state were obtained from the University of California Santa Cruz using the *UcscTrack()* function from the *Gviz* R package. Chromatin states of GM12878, H1-hESC, and NHLF cell lines are displayed based on their relevance to blood monocytes, prenatal development, and lung health and function, respectively.

**Supplementary Figure 34.**
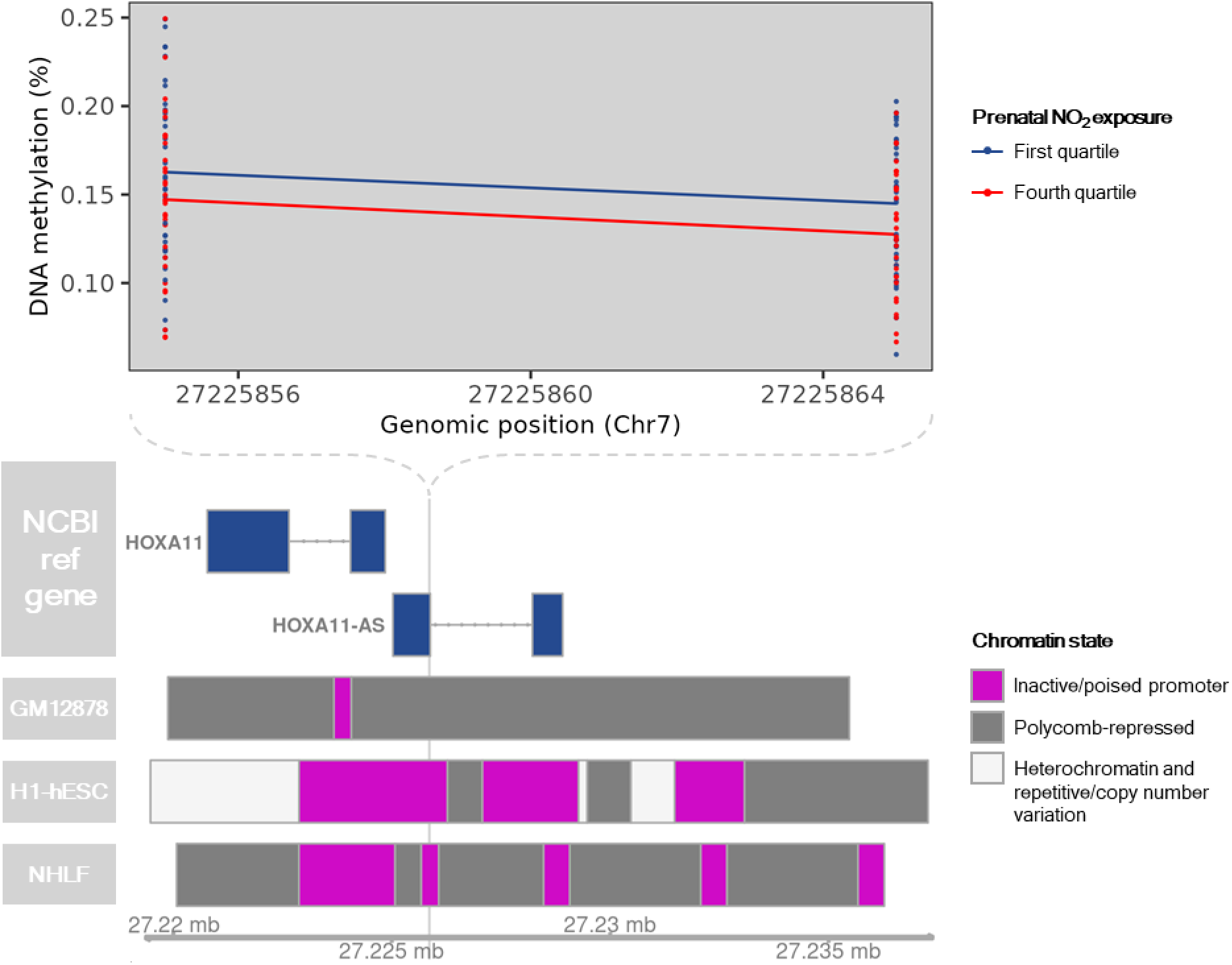
DNA methylation and chromatin state across the cord blood differentially methylated region (DMR) located on Chr7:27225855-27225866. DNA methylation at CpGs contained within Chr7:27225855-27225866 is shown for individuals with prenatal NO_2_ exposure < 7.3 ppb (first quartile; N=32; blue) or prenatal NO_2_ exposure > 16.3 ppb (fourth quartile; N=32; red) is displayed. Annotation of known genes and chromatin state were obtained from the University of California Santa Cruz using the *UcscTrack()* function from the *Gviz* R package. Chromatin states of GM12878, H1-hESC, and NHLF cell lines are displayed based on their relevance to blood monocytes, prenatal development, and lung health and function, respectively.

**Supplementary Figure 35.**
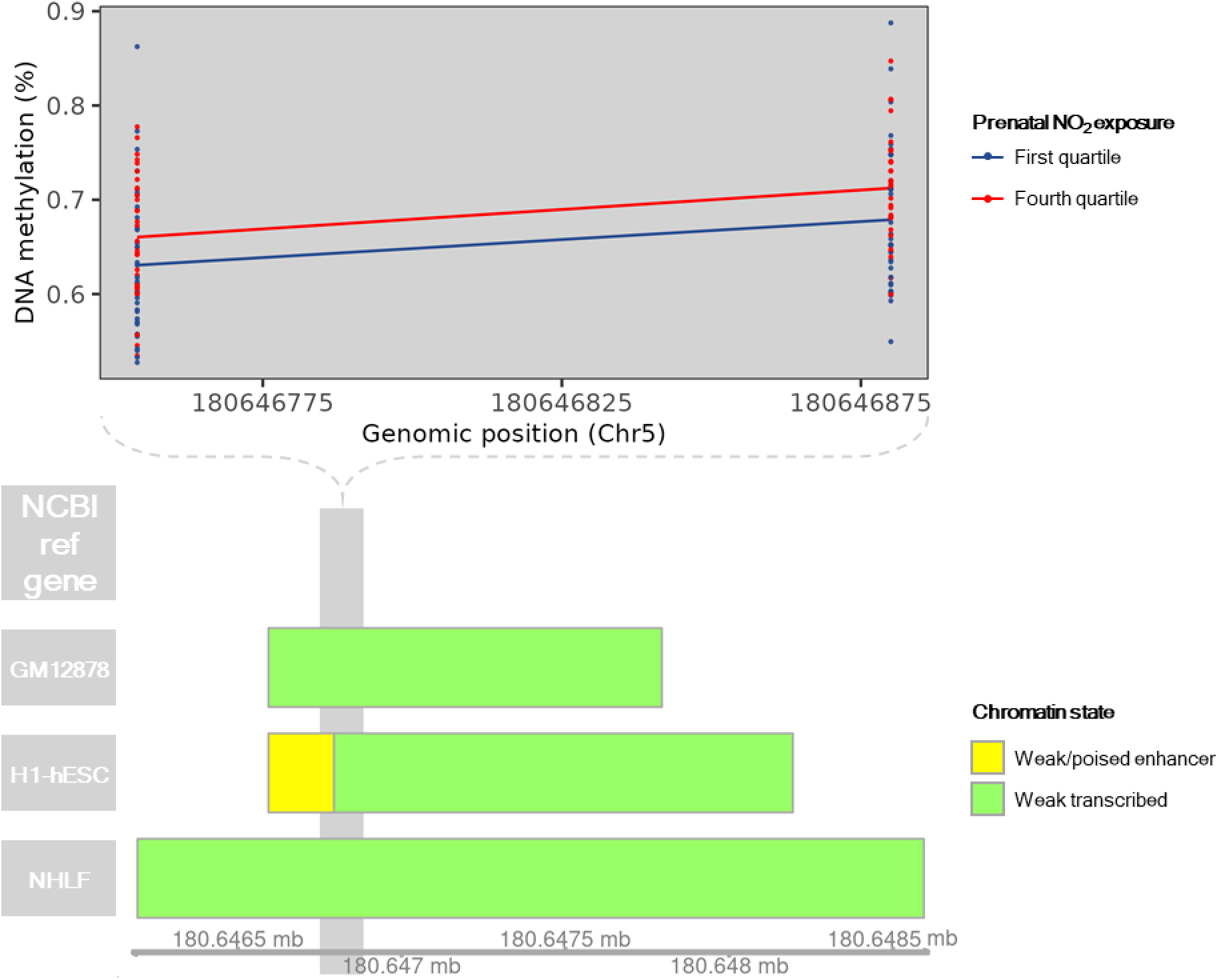
DNA methylation and chromatin state across the cord blood differentially methylated region (DMR) located on Chr5:180646754-180646881. DNA methylation at CpGs contained within Chr5:180646754-180646881 is shown for individuals with prenatal NO_2_ exposure < 7.3 ppb (first quartile; N=32; blue) or prenatal NO_2_ exposure > 16.3 ppb (fourth quartile; N=32; red) is displayed. Annotation of known genes and chromatin state were obtained from the University of California Santa Cruz using the *UcscTrack()* function from the *Gviz* R package. Chromatin states of GM12878, H1-hESC, and NHLF cell lines are displayed based on their relevance to blood monocytes, prenatal development, and lung health and function, respectively.

**Supplementary Figure 36.**
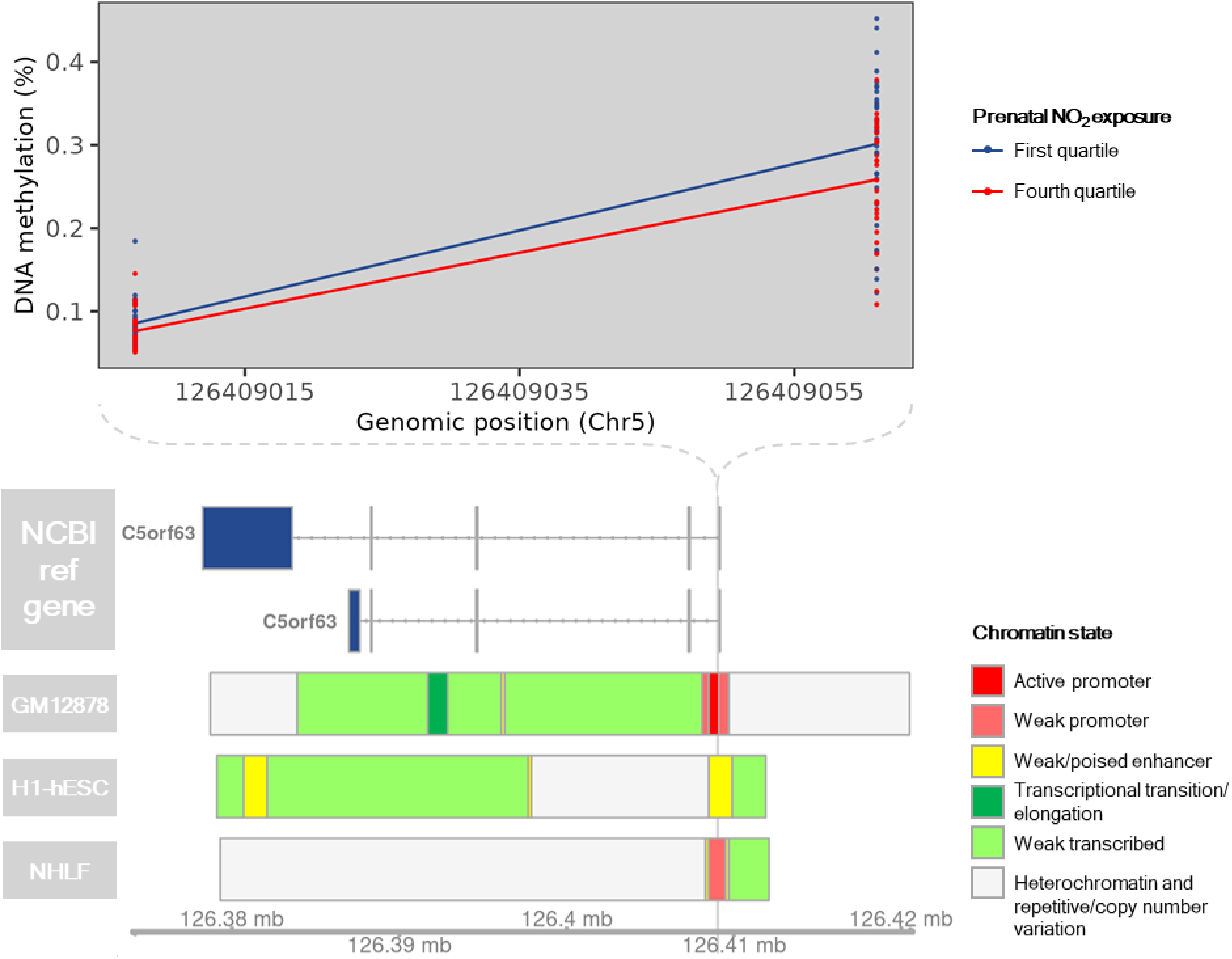
DNA methylation and chromatin state across the cord blood differentially methylated region (DMR) located on Chr5:126409007-126409062. DNA methylation at CpGs contained within Chr5:126409007-126409062 is shown for individuals with prenatal NO_2_ exposure < 7.3 ppb (first quartile; N=32; blue) or prenatal NO_2_ exposure > 16.3 ppb (fourth quartile; N=32; red) is displayed. Annotation of known genes and chromatin state were obtained from the University of California Santa Cruz using the *UcscTrack()* function from the *Gviz* R package. Chromatin states of GM12878, H1-hESC, and NHLF cell lines are displayed based on their relevance to blood monocytes, prenatal development, and lung health and function, respectively.

**Supplementary Figure 37.**
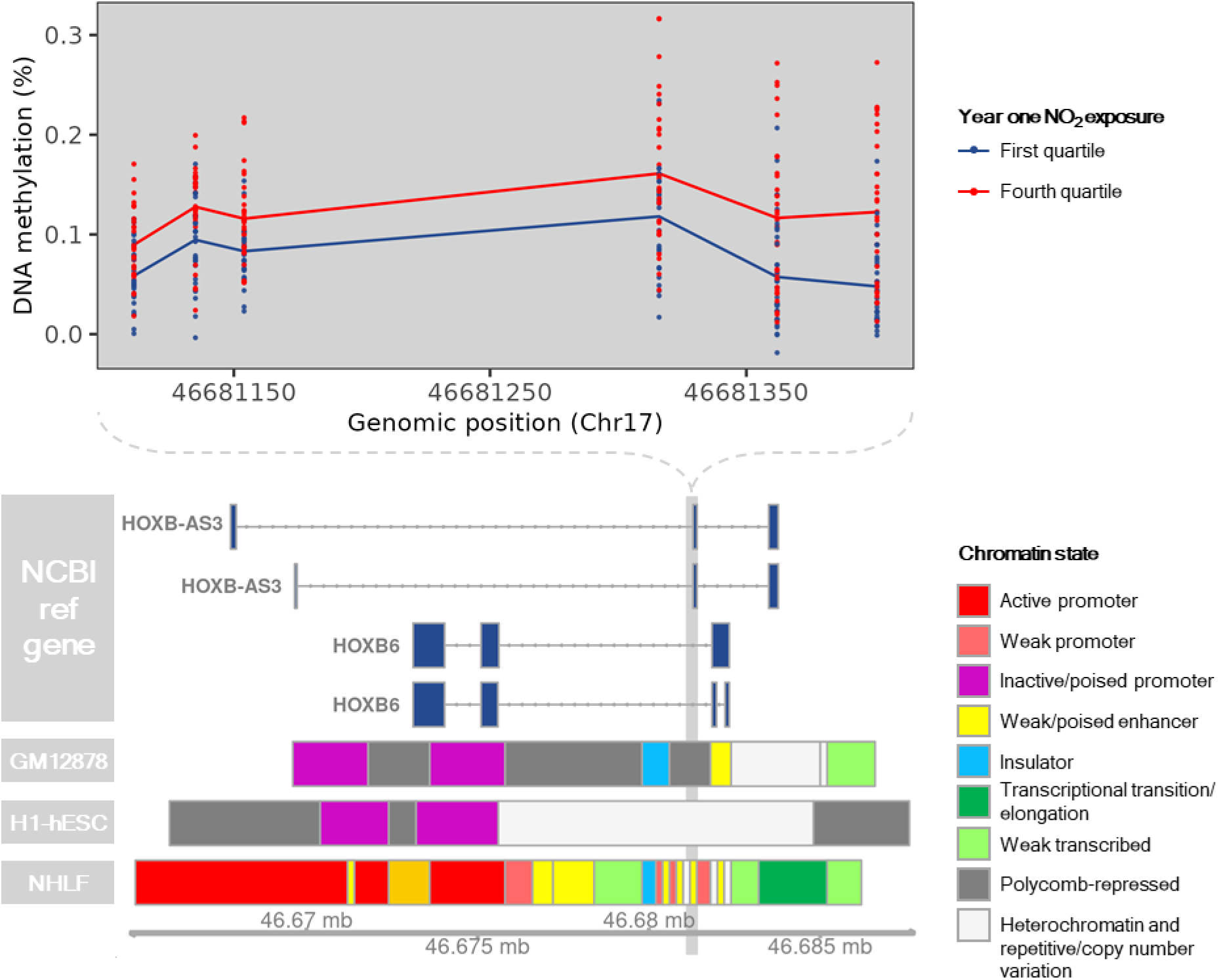
DNA methylation and chromatin state across the postnatal-specific differentially methylated region (DMR) located on Chr17:46681111-46681402. DNA methylation at CpGs contained within Chr17:46681111-46681402 is shown for individuals with prenatal NO_2_ exposure < 7.3 ppb (first quartile; N=32; blue) or prenatal NO_2_ exposure > 16.3 ppb (fourth quartile; N=32; red) is displayed. Annotation of known genes and chromatin state were obtained from the University of California Santa Cruz using the *UcscTrack()* function from the *Gviz* R package. Chromatin states of GM12878, H1-hESC, and NHLF cell lines are displayed based on their relevance to blood monocytes, prenatal development, and lung health and function, respectively.

**Supplementary Figure 38.**
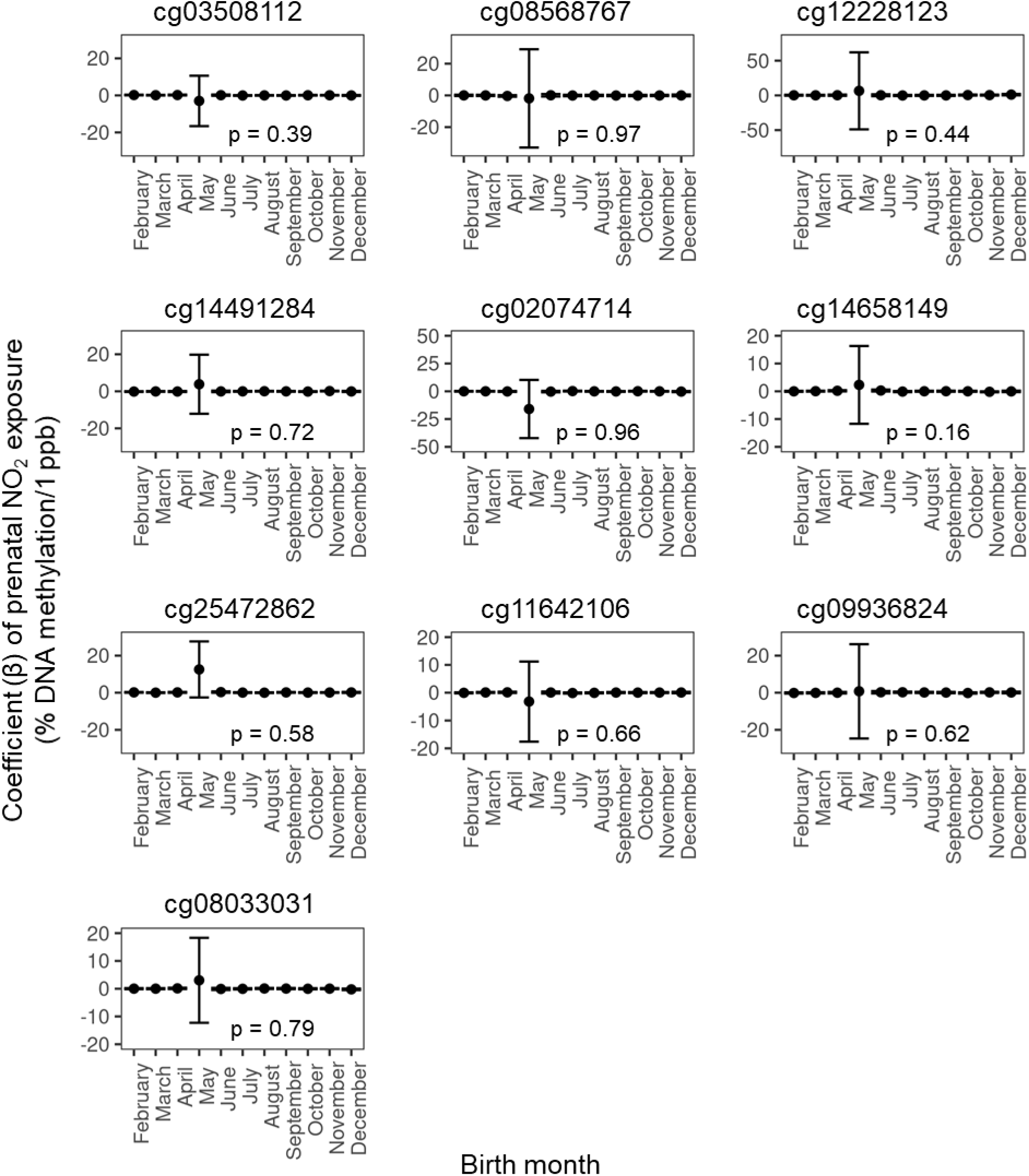
The effect of prenatal NO_2_ exposure on cord blood DNA methylation (DNAm) does not vary by birth month at the top 10 CpGs identified in the epigenome-wide analysis of cord blood (N=128). Based on ANOVA likelihood ratio tests, modelling the effect of prenatal NO_2_ exposure by birth month did not significantly improve model fit compared to models that did not include birth month. In all CpGs investigated, the month of May exhibited greater variation in the coefficient of prenatal NO_2_ exposure. This is likely spurious and related to a relatively small (N=3) number of CHILD participants born in May. The mean number of participants born in each month is 12 with a standard deviation of ±5.

**Supplementary Figure 39.**
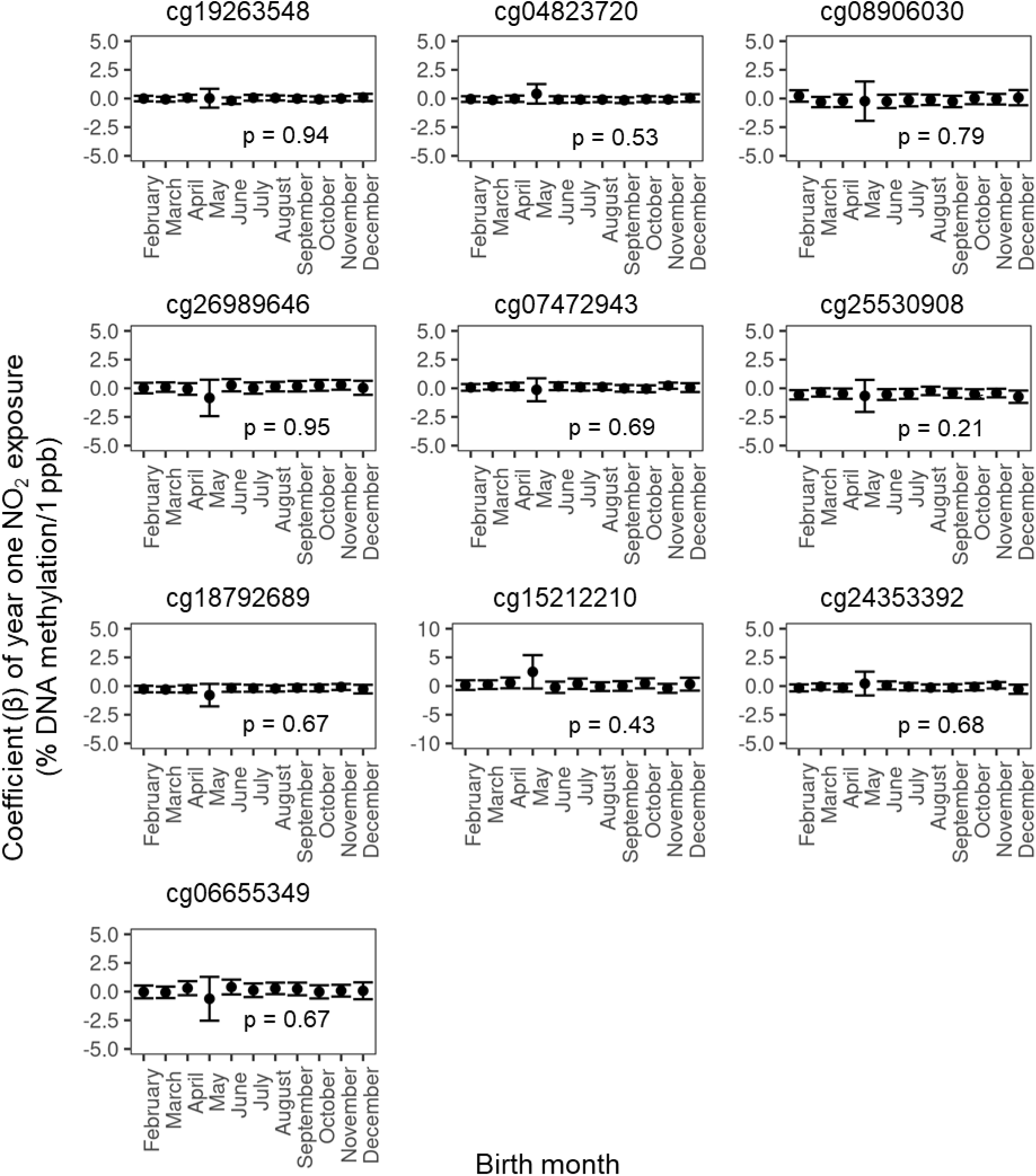
The effect of year one NO_2_ exposure on postnatal-specific DNA methylation (DNAm) does not vary by birth month at the top 10 CpGs identified in the epigenome-wide analysis of postnatal-specific DNAm changes. Based on ANOVA likelihood ratio tests, modelling the effect of year one NO_2_ exposure by birth month did not significantly improve model fit compared to models that did not include birth month. In all CpGs investigated, the month of May exhibited greater variation in the coefficient of year one NO_2_ exposure. This is likely spurious and related to a relatively small (N=3) number of CHILD participants born in May. The mean number of participants born in each month is 12 with a standard deviation of 5.

**Supplementary Table 1.**
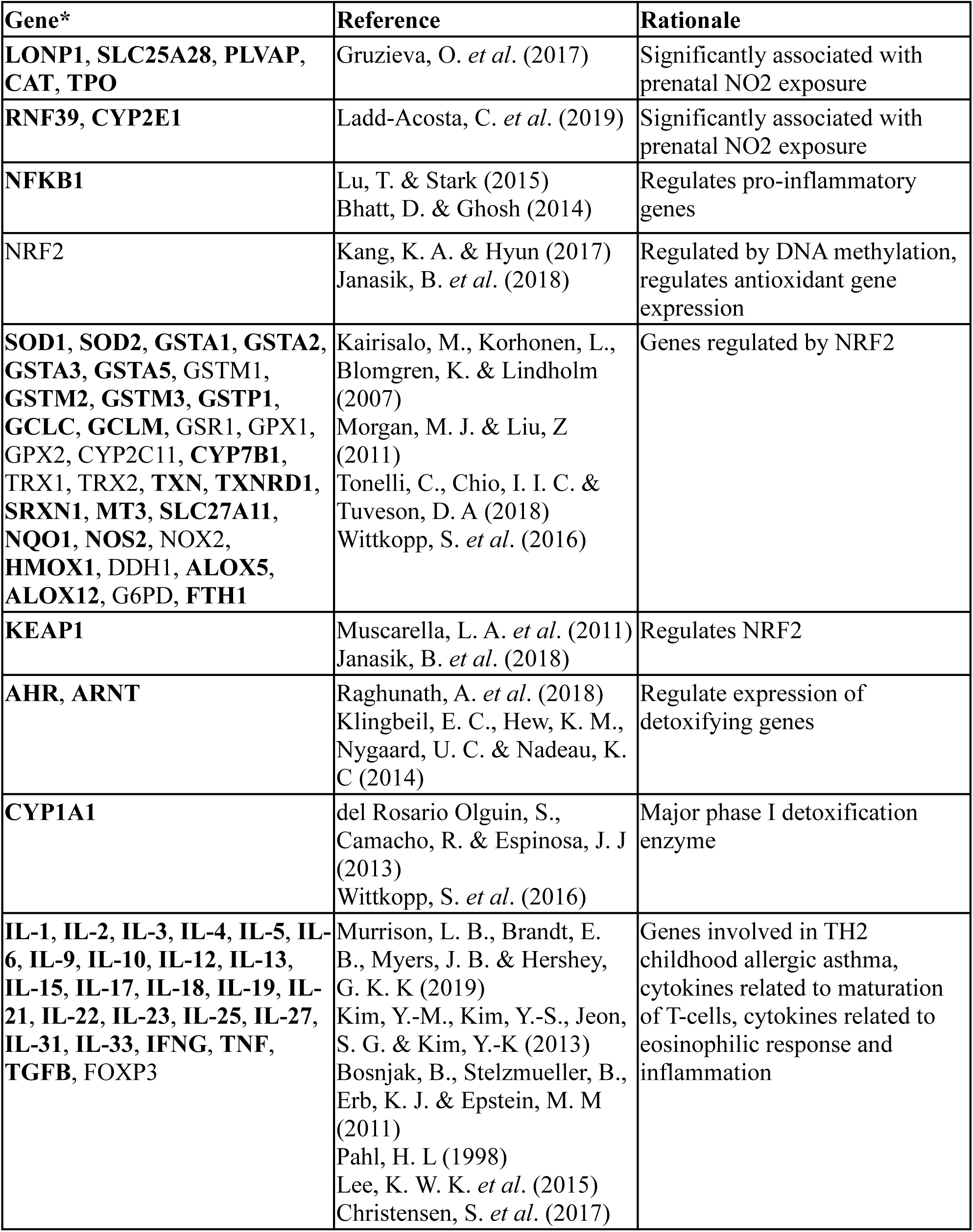
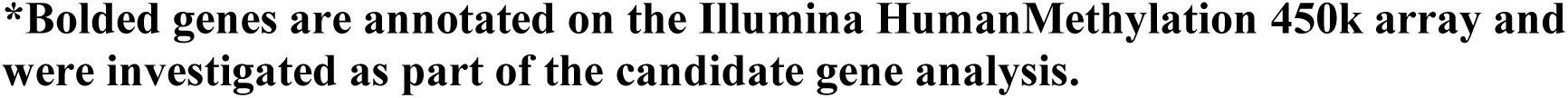
List of candidate genes investigated.

| Gene* | Reference | Rationale |
| --- | --- | --- |
| **LONP1, SLC25A28, PLVAP, CAT, TPO** | Gruzieva, O. *et al.* (2017) | Significantly associated with prenatal NO2 exposure |
| **RNF39, CYP2E1** | Ladd-Acosta, C. *et al.* (2019) | Significantly associated with prenatal NO2 exposure |
| **NFKB1** | Lu, T. & Stark (2015)<br>Bhatt, D. & Ghosh (2014) | Regulates pro-inflammatory genes |
| **NRF2** | Kang, K. A. & Hyun (2017)<br>Janasik, B. *et al.* (2018) | Regulated by DNA methylation, regulates antioxidant gene expression |
| **SOD1, SOD2, GSTA1, GSTA2, GSTA3, GSTA5, GSTM1, GSTM2, GSTM3, GSTP1, GCLC, GCLM, GSR1, GPX1, GPX2, CYP2C11, CYP7B1, TRX1, TRX2, TXN, TXNRD1, SRXN1, MT3, SLC27A11, NQO1, NOS2, NOX2, HMOX1, DDH1, ALOX5, ALOX12, G6PD, FTH1** | Kairisalo, M., Korhonen, L., Blomgren, K. & Lindholm (2007)<br>Morgan, M. J. & Liu, Z (2011)<br>Tonelli, C., Chio, I. I. C. & Tuveson, D. A (2018)<br>Wittkopp, S. *et al.* (2016) | Genes regulated by NRF2 |
| **KEAP1** | Muscarella, L. A. *et al.* (2011)<br>Janasik, B. *et al.* (2018) | Regulates NRF2 |
| **AHR, ARNT** | Raghunath, A. *et al.* (2018)<br>Klingbeil, E. C., Hew, K. M., Nygaard, U. C. & Nadeau, K. C (2014) | Regulate expression of detoxifying genes |
| **CYP1A1** | del Rosario Olguin, S., Camacho, R. & Espinosa, J. J (2013)<br>Wittkopp, S. *et al.* (2016) | Major phase I detoxification enzyme |
| **IL-1, IL-2, IL-3, IL-4, IL-5, IL-6, IL-9, IL-10, IL-12, IL-13, IL-15, IL-17, IL-18, IL-19, IL-21, IL-22, IL-23, IL-25, IL-27, IL-31, IL-33, IFNG, TNF, TGFB, FOXP3** | Murrison, L. B., Brandt, E. B., Myers, J. B. & Hershey, G. K. K (2019)<br>Kim, Y.-M., Kim, Y.-S., Jeon, S. G. & Kim, Y.-K (2013)<br>Bosnjak, B., Stelzmueller, B., Erb, K. J. & Epstein, M. M (2011)<br>Pahl, H. L (1998)<br>Lee, K. W. K. *et al.* (2015)<br>Christensen, S. *et al.* (2017) | Genes involved in TH2 childhood allergic asthma, cytokines related to maturation of T-cells, cytokines related to eosinophilic response and inflammation |

**\*Bolded genes are annotated on the Illumina HumanMethylation 450k array and** **were investigated as part of the candidate gene analysis.**

**Supplementary Table 2.**
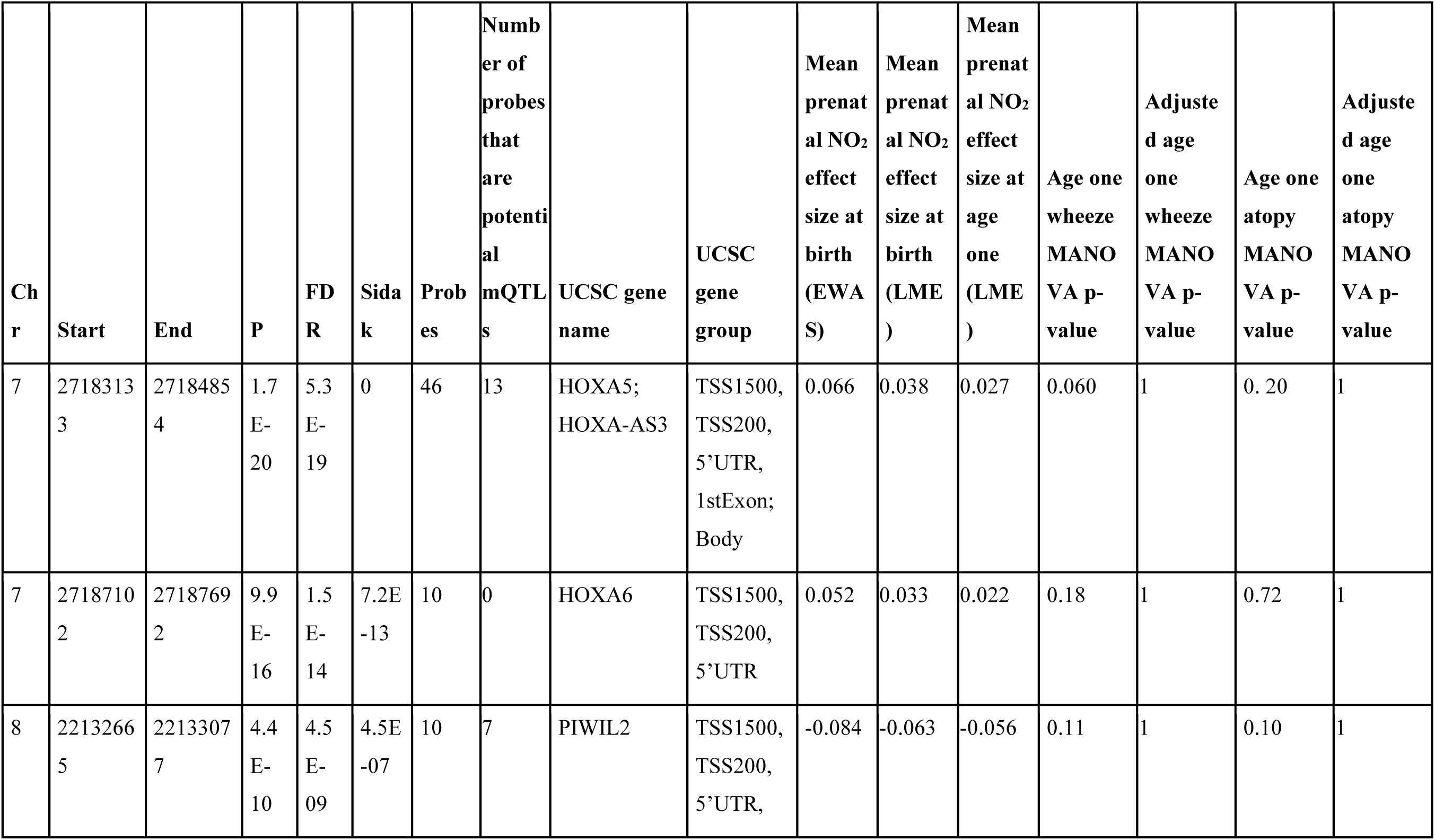
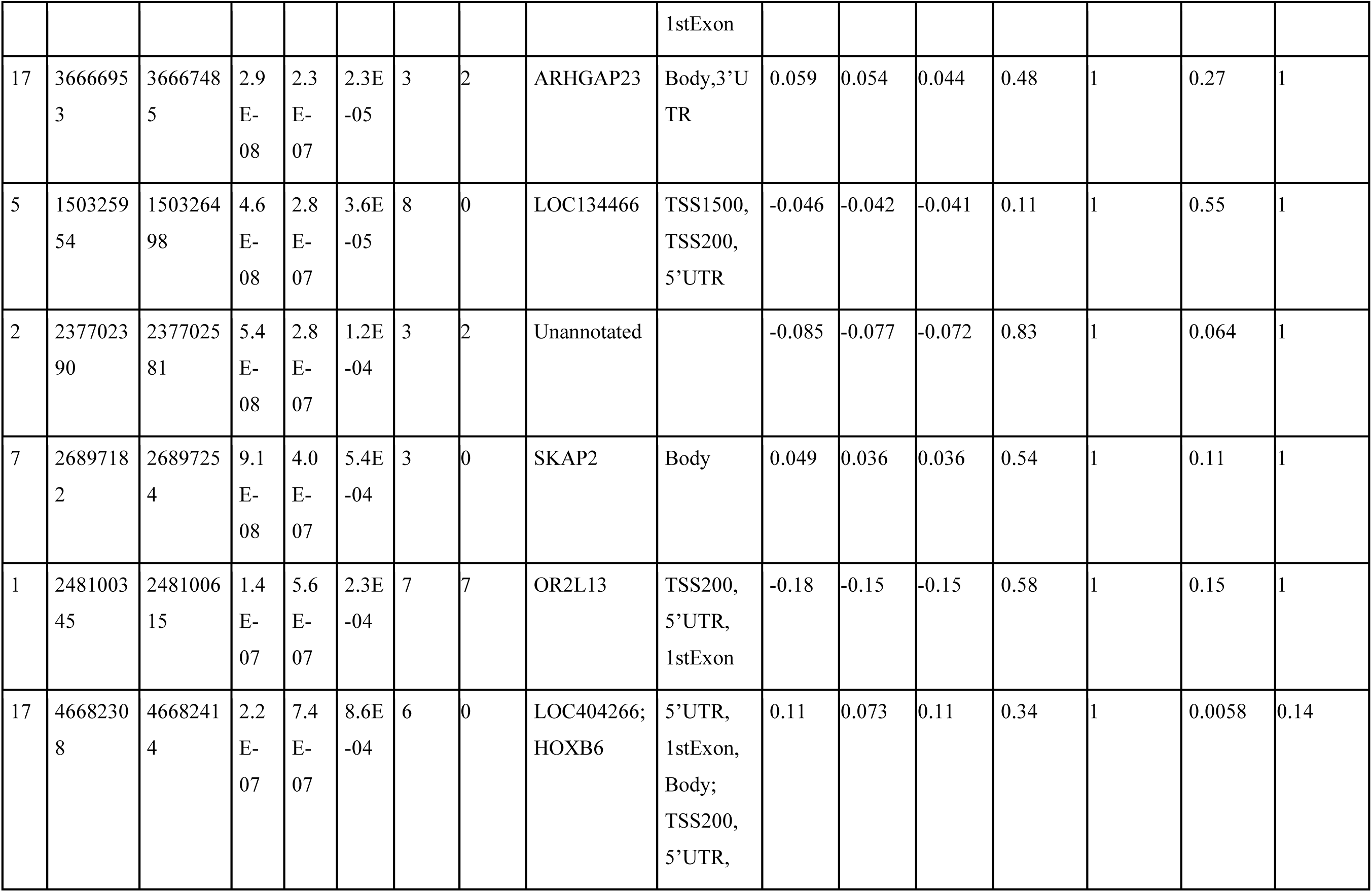
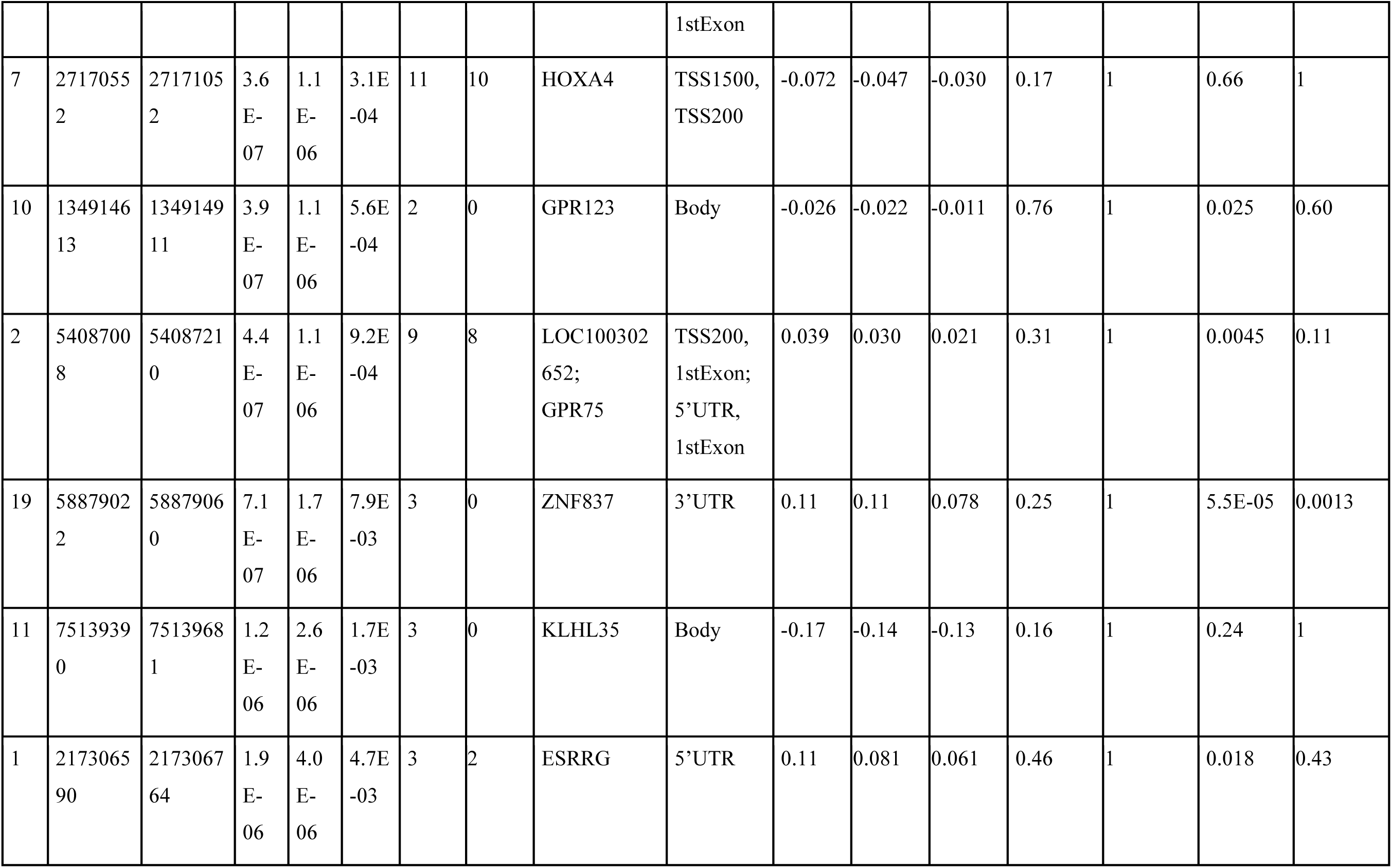
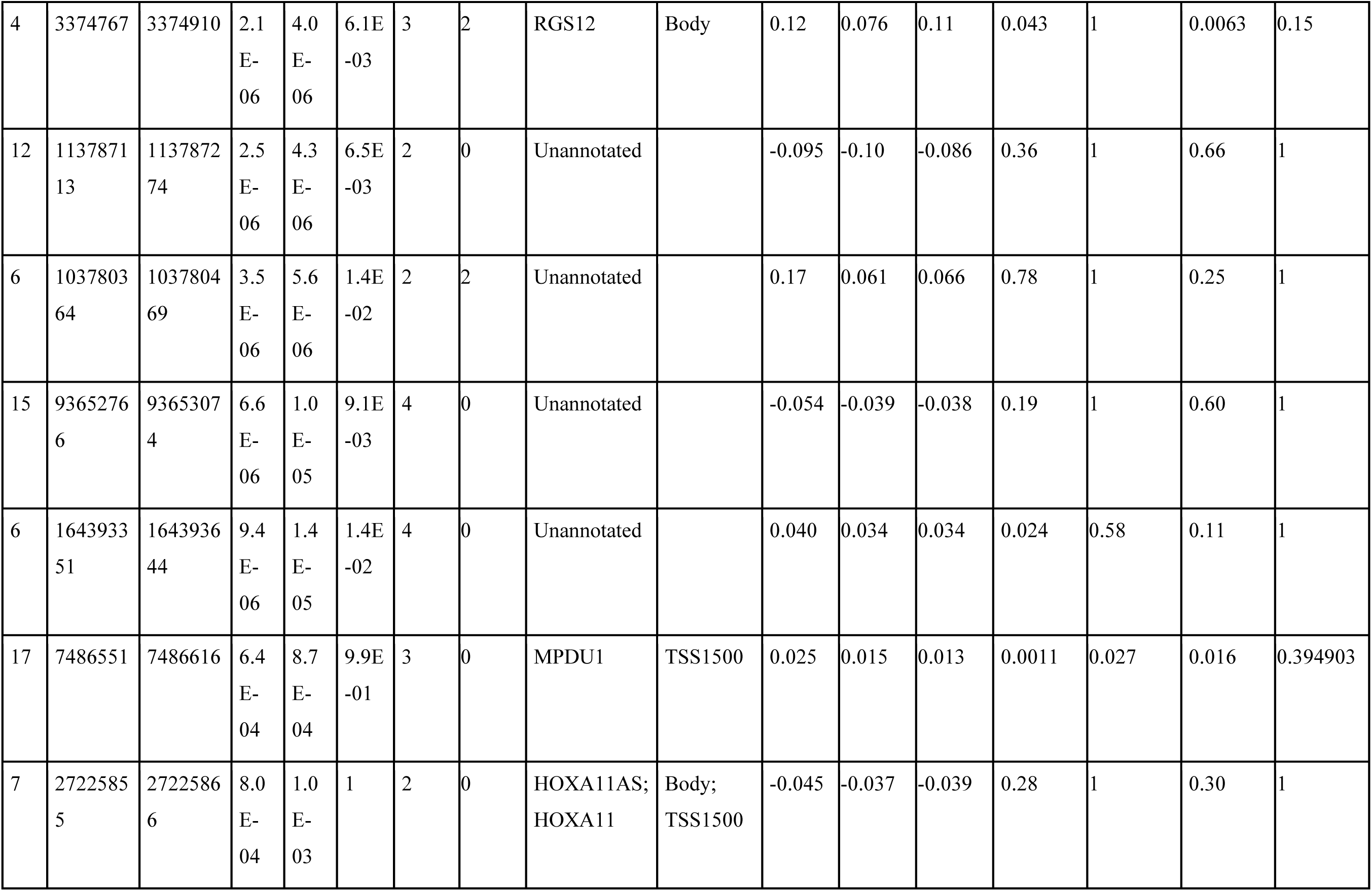
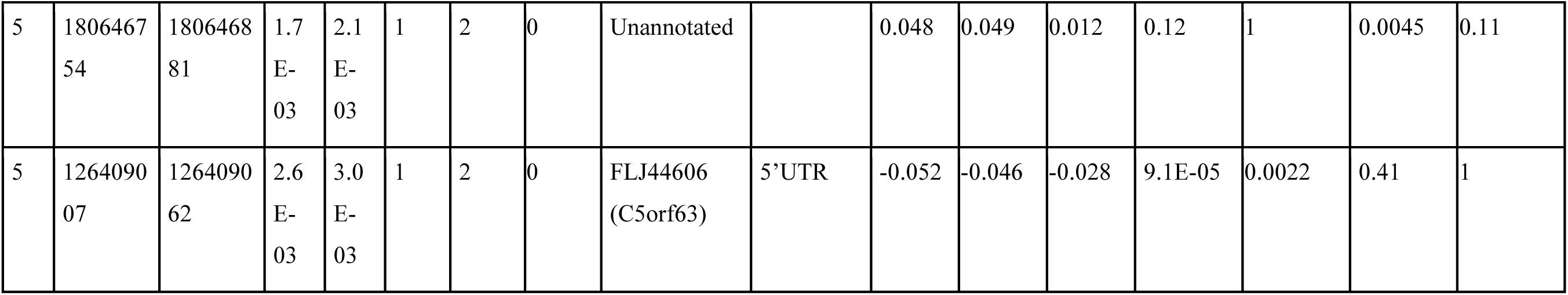
Genomic information of cord blood differentially methylated regions (N=24) associated with prenatal NO_2_ exposure.

